# Characterizing the metabolomes of microglia, astrocytes, and neurons in aging and Alzheimer’s brains

**DOI:** 10.64898/2026.02.06.703980

**Authors:** Jie Yu, Fengzhi Li, Xing-jun Chen, Chenye Mou, Di Yao, Zhanying Bi, Xiaoli Chen, Lulu Du, Ziyan Feng, Xinshuang Zhang, Xiaoqian Yu, Lauren G. Zacharias, Ralph J. DeBerardinis, Li Zhang, Zhen Li, Benyan Luo, Xiaoling Hu, Woo-ping Ge

## Abstract

Neurons and glia are distinct in their morphology, development, and function, possessing unique transcriptomes and proteomes, but little is known about their metabolomes. The challenge of brain cell metabolic profiling is to obtain a large number of cells for reliable analysis. Here, we purified microglia, astrocytes, and neurons from mouse brains, identifying >70 metabolites through targeted metabolomics and 9,854 metabolite features via untargeted metabolomics. We systematically characterized cell type–enriched metabolites and metabolic pathways, revealing an enrichment of glutathione (GSH) and polyamine metabolism in microglia. This enrichment was validated *in vivo* and showed significant decreases with aging and in an Alzheimer’s disease (AD) model. Notably, GSH and polyamine metabolism correlated strongly with chemokine-related gene expression. Disrupting the GSH pathway in microglia resulted in downregulation of chemokine-related genes, aberrant morphogenesis, and β-amyloid deposition. Our results provide a valuable resource (https://metabolismocean.org/braincell) for metabolic studies related to aging, AD, and other neurological diseases.

## Introduction

The brain is an organ with an especially high energy requirement ^1–4^. Glial cells and the brain vasculature form a complex network with neurons and provide neurons with nutritional support to maintain their physiological functions ^5^. Astrocytes are the most abundant glial population in the brain. They take up glutamate released from synapses via excitatory amino acid transporters and in return supply neurons with glutamine ^2,3^. Their endfeet cover ∼99% of the abluminal vascular surface in the adult brain ^6,7^ and ferry glucose from the blood to neurons, which is vital for neuronal survival and function ^8,9^. Microglia represent 5–10% of the total brain cell population ^10,11^. They are the resident immune cells of the central nervous system^12–16^. As critical players in neuroinflammatory and neurodegenerative disease, microglia have become an important target for neurological diseases ^11,17^. In addition, microglia actively respond to brain metabolism alterations, especially under pathological conditions or during aging ^18,19^. However, to date, little is known about their own metabolism.

Metabolic abnormalities in the brain can lead to many disorders, including neurodegenerative diseases ^20,21^, neuroimmune defects ^22^, and epilepsy ^23^. Abnormal metabolic communications among various brain cells result in the collapse of entire metabolic networks, eventually leading to neuronal degeneration or death ^24–26,27–30^. For example, TREM2-deficient microglia show defects in glycolysis, ATP production, and cholesterol metabolism, exacerbating the progression of Alzheimer’s disease (AD) ^31–33^. Dysfunctions that affect glutamate transporters and the glutamate-converting enzyme glutamine synthetase (GS) in astrocytes are important causative factors for seizures and neurotoxicity ^34,35^. The glutamate-glutamine cycle between astrocytes and neurons is a notable contributor to GABA (γ-aminobutyric acid) and glutamate release from synapses ^36^. Therefore, characterizing the metabolomes of brain cells will help us to better understand their physiological functions and the mechanisms underlying various neurological diseases.

Metabolomics has been widely used to study the brain and various neurological diseases ^37–39,4,40–42^. However, these studies have been limited to a block of brain tissue or to the cerebral spinal fluid ^29,43–45^, which makes it tremendously challenging to elucidate the contributions of certain types of brain cells to the metabolites. Other studies have focused on one particular metabolite in one or more cell types in the brain, such as the lactate and pyruvate that are provided to neurons by astrocytes for energy metabolism ^46^ or the increase in taurine levels during the differentiation and maturation of oligodendrocyte precursor cells ^47^. Nevertheless, metabolic disorders of brain cells usually involve the dysfunction of various pathways or networks rather than the alteration of individual metabolites ^48,49^. The field still lacks a comprehensive metabolome dataset from various brain cells as a reference that would allow researchers to directly compare metabolic profiles among neurons and glial cells for mechanistic studies.

Obtaining high-quality metabolomic data depends on the high purity, viability, and yield of cells isolated from a brain. In contrast to the metabolic profiling of cells *in vitro*, a metabolic study requires purifying neurons and glial cells from adult mouse brains, which is difficult ^50–52^. During the process of purification, excitatory toxicity and mechanical dissociation lead to the death of most brain cells ^53–59^. Here, we constructed a metabolic database for these three types of cells in aging and AD models. By integrating the transcriptome data from isolated neurons, astrocytes, and microglia with our metabolomic analyses, we discovered the crucial role of microglial glutathione (GSH) metabolism in the pathological advancement of AD.

## Results

To obtain a large number of brain cells, we used different transgenic mouse strains to label neurons and glial cells with EGFP or EYFP. We then isolated these cells from the cerebral cortex and hippocampus before we performed metabolomics and bulk RNA-seq for metabolome and transcriptome analyses (**Supplementary Fig. 1**).

### Targeted metabolomic profiling of microglia

In brain sections from adult *Cx3cr1*^+/EGFP^ mice^60^, EGFP fluorescence was uniquely expressed in microglia (**Fig. 1a**). We stained brain sections with anti-Iba1, an antibody for microglia. All EGFP^+^ cells in the cerebral cortex and hippocampus were Iba1^+^ (100.0%, n = 2,281 cells; **Figs. 1b, c**). Thus, EGFP^−^ cells from the same brain were used as the control group.

**Fig. 1.**
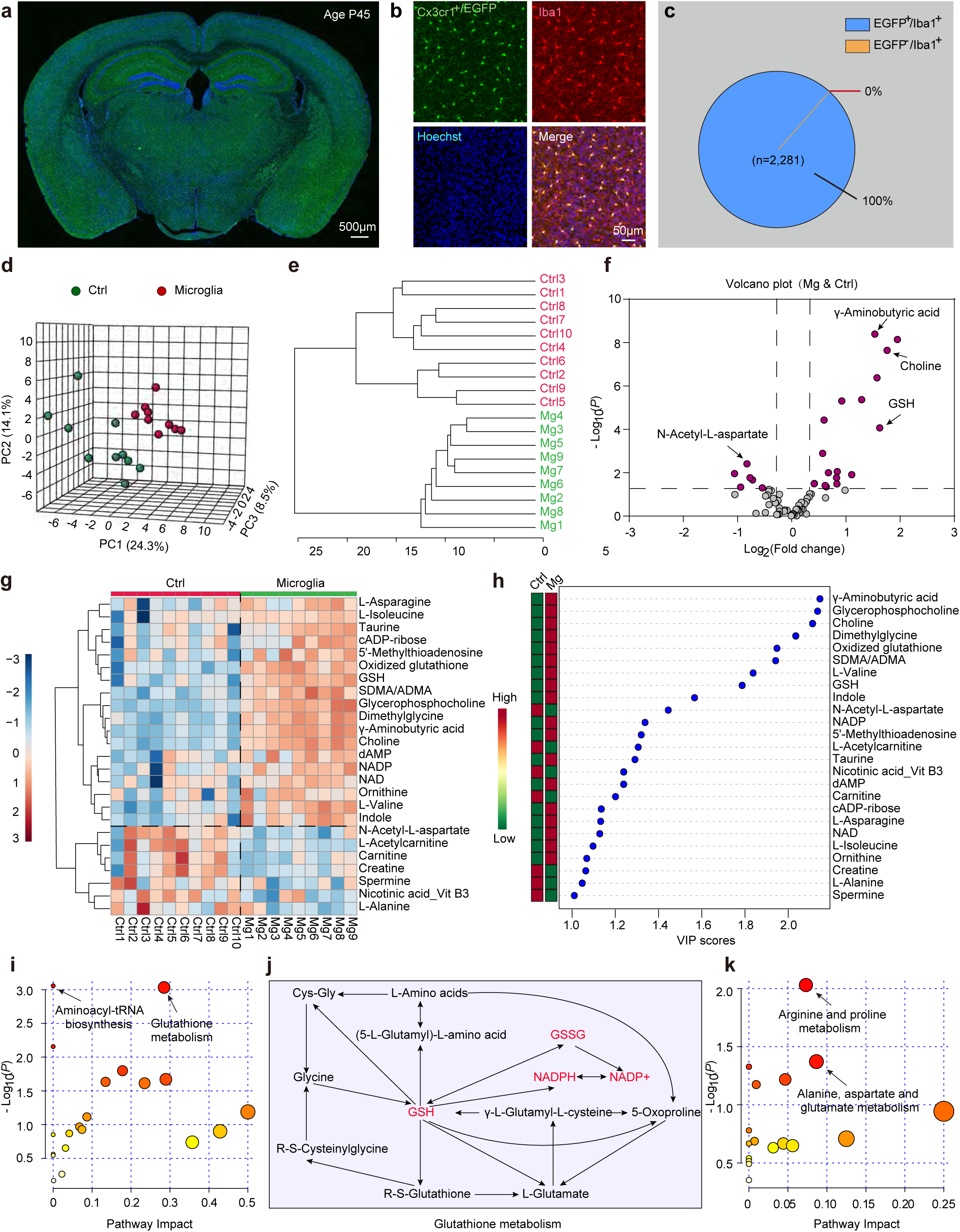
The metabolome of microglia identified by targeted metabolomics. **(a, b)** Image of a coronal section (a) and microglia (green, EGFP) in the in the cerebral cortex (b) from the brain of a postnatal(P)45 *Cx3cr1*^+/EGFP^tg mouse. microglia, anti-Iba1(red); nuclei, Hoechst 33342 (Hoechst, blue). **(c)** Percentage of EGFP^+^ cells (Iba1^+^EGFP^+^ cells/Iba1^+^ cells×100%) in microglia from (b). **(d)** Principal component analysis (PCA) of the metabolomes of microglia (n = 9 samples) and the control group (i.e., EGFP^−^ cells, n = 10 samples). Each dot represents a sample from a different mouse. **(e)** Dendrogram showing the clustering of all samples from the control (Ctrl1–10) and microglial groups (Mg1–9). **(f)** Volcano plot of differential metabolites (fold change > 1.2 and *P* **<** 0.05, purple, a two-tailed unpaired *t*-test). Non-differential, gray. A data point with a positive value indicates a high level of a metabolite in microglia. **(g)** Heatmap representation of 25 of 73 metabolites in the control (n = 10 samples) and microglia (n = 9 samples) groups, as identified by VIP scores from a PLS-DA. Each column represents a sample from a different mouse. **(h)** Top 25 metabolites based on VIP scores for differentiating between microglia and the control group. The colored boxes on the left indicate the relative concentrations of corresponding metabolites averaged across each group. **(i, k)** A summary of the pathway analysis using metabolites found at high levels (fold change > 1.2) in microglia as compared with the control group (i) or in the control group as compared with microglia (k). Pathway enrichment was performed using a hypergeometric test and pathway topology analysis. **(j)** A portion of the glutathione metabolism pathway in microglia and the control group. Metabolites in red were present at a high level in microglia as compared with the control group. The size of the circle represents pathway impact, and the color represents the *P*-value. Smaller *P*-values and larger pathway impacts indicate more influential pathways. NAD, nicotinamide adenine dinucleotide; NADP, nicotinamide adenine dinucleotide phosphate; SDMA/ADMA, symmetric dimethylarginine/asymmetric dimethylarginine; Vit, vitamin; GSH, glutathione (reduced); Ctrl, control; Mg, microglia.

We used ∼50,000 microglia per sample for the targeted metabolomic analysis. Among 204 metabolites in our targeted library, 73 metabolites were reliably identified from these samples. The metabolic profiles of the microglial and control groups were different (**Fig. 1d, e**). Of the 73 metabolites we identified, 18 metabolites, including γ-aminobutyric acid, choline, and GSH, were present at substantially higher concentrations in microglia than in the control group, whereas 6 other metabolites, including *N*-acetyl-L-aspartate, were present at a lower concentration in microglia **(Fig. 1f**). We further performed a partial least-squares discriminant analysis (PLS-DA) to identify the metabolites that maximized the separation between microglia and the control group. We identified 25 metabolites that maximized the separation, of which L-asparagine, L-isoleucine, taurine, cADP-ribose, 5’-methylthioadenosine, oxidized glutathione (GSSG), GSH, symmetric dimethylarginine/asymmetric dimethylarginine (SDMA/ADMA), glycerophosphocholine, dimethylglycine, choline, dAMP, NADP, NAD, ornithine, L-valine, and indole were enriched in microglia (**Figs. 1g, h**).

To identify the dominant metabolic pathways in microglia, we analyzed the global metabolic pathway using the metabolites with a fold change > 1.2. Among 20 metabolic pathways, the GSH metabolism pathway and the Aminoacyl-tRNA biosynthesis pathway showed the most substantial differences between microglia and the control group (**Fig. 1i**). In the GSH metabolism pathway, the levels of metabolites GSH, NADPH, and NADP^+^ were higher in microglia as compared with the control group (**Fig. 1j**). We also analyzed those pathways in which the metabolites were at lower levels in microglia and observed that the Arginine and proline metabolic pathways and Alanine, aspartate, and glutamate metabolic pathways differed notably between these two groups (**Fig. 1k**). To further determine the most dominantly enriched metabolites in microglia, we identified 10 metabolites with significant differences based on variable importance in projection (VIP) > 1 and *P* < 0.05. Of these, 9 metabolites, glycerophosphocholine, choline, dimethylglycine, GSH, SDMA/ADMA, γ-aminobutyric acid, L-valine, GSSG, and indole, were present at significantly higher levels in microglia (**Extended Data Fig. 1a–i**). In contrast, the level of *N*-acetyl-L-aspartate was dramatically lower in microglia (**Extended Data Fig. 1j**).

### Targeted metabolomic profiling of astrocytes

To collect astrocytes from the mouse brain, we used *Aldh1l1-EGFP* transgenic (*Aldh1l1^EGFP^tg*) mice^61,62^. We verified that EGFP^+^ cells in the cerebral cortex and hippocampus were astrocytes through labeling with anti-GFAP (99.0%, n=1,215 cells; **Extended Data Fig. 2a–c**) or anti-Sox9 (99.0%, n = 1,186 cells in the cortex; **Fig. 2a and b**; 87.5%, n = 1,922 cells in the hippocampus; **Fig. 2c**), two antibodies specific for astrocytes. After extracting metabolites from ∼50,000 sorted astrocytes (i.e., EGFP^+^ cells) in the cerebral cortex and hippocampus, we performed targeted metabolomics on both the astrocyte group and the control group (i.e., EGFP^−^ cells, representing non-astrocyte cells). Among 204 metabolites in our targeted measurements, 82 metabolites were reliably identified in both groups. Based on principal component analysis (PCA) and clustering analysis, 12 of the obtained samples were well separated into two clusters (**Fig. 2d, 2e**). Compared to their relative concentrations in the control group, 12 metabolites, including putrescine, L-histidine, and 7-methylguanosine, had higher levels in astrocytes (fold change > 1.2 and *P* < 0.05; **Fig. 2f**). In contrast, the levels of 32 metabolites, including GSH, GSSG, taurine, carnitine, and 2-isopropylmalate, were lower in astrocytes (**Fig. 2f**). With PLS-DA analysis, we isolated 12 metabolites that were enriched in astrocytes (**Fig. 2g, h**).

**Fig. 2.**
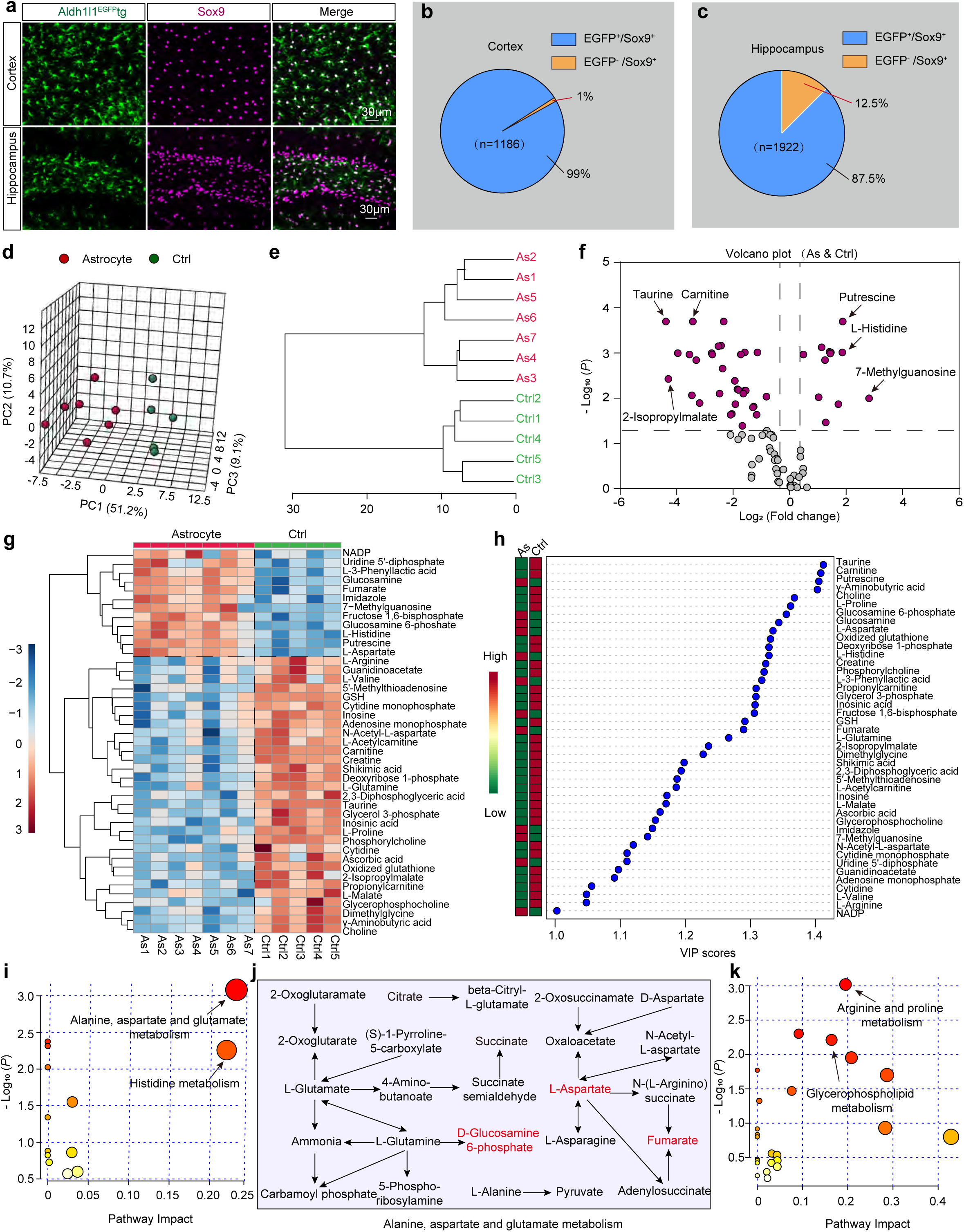
The metabolome of astrocytes identified by targeted metabolomics. **(a–c)** Image of astrocytes in the hippocampus and cortex of an *Aldh1l1^EGFP^*tg mouse (a). Astrocytes (Sox9^+^, purple) with EGFP (EGFP^+^Sox9^+^) or without EGFP (EGFP^−^Sox9^+^) expression were included for analysis in (b and c). Percentage was calculated as 100% × (Sox9^+^EGFP^+^ cells/Sox9^+^ cells). **(d)** PCA of the metabolomes of astrocytes and the control group (i.e., EGFP^−^ cell group). In total, 82 metabolites identified from each sample were used for PCA. Circles indicate individual samples from astrocytes (n = 7 samples) and the control group (n = 5 samples). **(e)** Dendrogram showing the clustering of all samples. Ctrl1–5, control group (n = 5 samples); As1–7, astrocyte group (n = 7 samples). **(f)** Volcano plot of differential metabolites between astrocytes and the control group as described in Figure 1. A data point with a positive value indicates a high level of a metabolite in astrocytes. **(g)** Heatmap representation of 43 of 82 metabolites in the control (n = 5 samples) and astrocyte (n = 7 samples) groups, as identified by VIP scores from a PLS-DA. Each column represents a sample from a different mouse. Red indicates a greater abundance of the metabolite. **(h)** Top 43 metabolites based on VIP scores for differentiating between astrocytes and the control group. The colored boxes on the left indicate the relative concentrations of the corresponding metabolites averaged across each group. **(i, k)** A summary of the pathway analysis using metabolites found at high levels (fold change > 1.2) in astrocytes as compared with the control group (i) or in high levels in the control group as compared with astrocytes (k). Pathway enrichment was performed using a hypergeometric test and pathway topology analysis. The size of the circle represents pathway impact, and the color represents the *P*-value. Smaller *P*-values and larger pathway impacts indicate more influential pathways. **(j)** A portion of the Alanine, aspartate, and glutamate metabolic pathway in astrocytes and the control group. Metabolites in red were present at high levels in astrocytes as compared with the control group. GSH, glutathione (reduced).

To identify the metabolic pathways that were uniquely dominant in astrocytes, we carried out a global metabolic pathway analysis using those metabolites with a fold change > 1.2 and *P* < 0.05. Among the pathways, the Alanine, aspartate, and glutamate metabolic pathway and Histidine metabolic pathway showed the most notable differences between astrocytes and the control group (**Fig. 2i**). In the Alanine, aspartate, and glutamate metabolic pathway, the metabolites L-aspartate, fumarate, and D-glucosamine 6-phosphate were present at substantially higher concentrations in astrocytes than in the control group (**Fig. 2j**). Those pathways that involved metabolites present at lower levels in astrocytes were also analyzed. The Arginine and proline metabolic pathway and Glycerophospholipid metabolic pathway were notably lower in astrocytes as compared with the control group (**Fig. 2k**). Among all differential metabolites (highest VIP, *P* < 0.05), taurine, carnitine, L-proline, γ-aminobutyric acid, choline, and GSSG were present at significantly higher levels in the control group (**Extended Data Fig. 2d-i**), whereas glucosamine 6-phosphate, glucosamine, L-aspartate, and putrescine were present at significantly higher levels in astrocytes (**Extended Data Fig. 2j–m**).

### Targeted metabolomic profiling of neurons

To collect neurons from the brain, we used *Thy1-YFP-16J tg* (*Thy1^YFP^tg*) mice^63^. After staining brain sections with an antibody against the neuronal nuclear protein (NeuN), we observed a high density of neurons with YFP expression in the cerebral cortex and hippocampus (**Extended Data Fig. 3a, b**). Although YFP^+^NeuN^+^ cells made up the majority among all NeuN^+^ neurons in the cerebral cortex and hippocampus (YFP^+^NeuN^+^ cells/all NeuN^+^ cells, 79.3%, n = 2,275 cells), all YFP^+^ cells in these two regions were NeuN^+^ (100.0%, n = 2,275 cells; **Extended Data Fig. 3c**).

We then performed targeted metabolomic analysis of ∼50,000 neurons. Based on a PCA and cluster analysis, the metabolomes of neurons and the control group were not well separated, although most of the neuronal samples were clustered together (**Extended Data Fig. 3d, e**). This is likely because YFP^+^ cells did not represent all neurons in *Thy1^YFP^tg* mice; ∼20% of NeuN^+^ neurons were YFP^−^ and were included in the control samples during sorting. Also, some inhibitory neurons in the cerebral cortex and hippocampus are YFP^−^ in *Thy1^YFP^tg* mice, which is consistent with the result from the previous report ^63^. Among the 204 metabolites in our targeted measurements, 82 metabolites were reliably identified in both groups. The levels of 11 metabolites, including glucosamine 6-phosphate, L-histidine, and L-3-phenylacetic acid, were present at higher levels in neurons, and 8 metabolites, including carnitine, creatine, and L-proline, were found at lower levels in neurons (fold change > 1.2 and *P* < 0.05; **Extended Data Fig. 3f**). Using a PLS-DA analysis, we found that 15 of 34 metabolites—gluconic acid, adenosine, guanosine monophosphate (GMP), uridine 5’-diphosphate (UDP), acetylcholine, imidazole, glucosamine 6-phosphate (G6P), 2-phosphoglyceric acid, glucosamine, putrescine, L-3-phenylacetic acid, methionine sulfoxide, 2-hydroxyglutaric acid, L-histidine, and L-aspartate—were enriched in neurons (**Extended Data Fig. 3g, h**).

We further performed metabolic pathway analysis with these neuron-enriched metabolites. Four pathways, namely the Alanine, aspartate, and glutamate metabolic pathway; the Histidine metabolic pathway; the Tricarboxylic acid (TCA) cycle; and the Arginine biosynthesis metabolic pathway, were highly enriched in neurons (**Extended Data Fig. 3i**). Among the metabolites in the Alanine, aspartate, and glutamate metabolic pathway, citrate, succinate, L-aspartate, fumarate, and glucosamine 6-phosphate showed a higher level in neurons than in the control group (**Extended Data Fig. 3j**). The metabolic pathways enriched in the control group included the Arginine and proline metabolic pathway and the glutathione metabolic pathway (**Extended Data Fig. 3k**). Additionally, the levels of creatine, carnitine, and L-proline were significantly higher in the control group (**Extended Data Fig. 4a–c**), whereas L-aspartate, glucosamine, L-histidine, L-3-phenylacetic acid, 2-hydroxyglutaric acid, putrescine, and glucosamine 6-phosphate were significantly higher in neurons (**Extended Data Fig. 4d–j**).

### Comparison of the targeted metabolomes of microglia, astrocytes, and neurons

To identify unique metabolites in different brain cells, we compared the metabolites identified in neurons and glial cells. Based on PCA and cluster analysis, the metabolomes from the same cell type clustered together under a correlation analysis, indicating a high correlation among samples of the same cell type (**Extended Data Fig. 5a–c**). With a PLS-DA analysis, we identified 35 metabolites enriched in distinct cell types (**Extended Data Fig. 5d, 5e**). Of these metabolites, NADP, 7-methylguanosine, imidazole, and fumarate were enriched in astrocytes, and metabolites including betaine, adenosine monophosphate (AMP), cytidine, 4-hydroxyproline, and L-lactate were enriched in neurons. Interestingly, some metabolites that were enriched in neurons, including GSSG, betaine, deoxyribose 1-phosphate, carnitine, taurine, GSH, cytidine monophosphate (CMP), dimethylglycine, γ-aminobutyric acid, choline, L-proline, L-arginine, L-valine, 4-hydroxyproline, inosinic acid, and phosphorylcholine, were also enriched in microglia (**Extended Data Fig. 5d**).

We additionally performed comparisons of the metabolites among the three analyzed types of cells. The levels of L-palmitoylcarnitine, pantothenic acid, and adenosine monophosphate were higher in neurons than in microglia (**Extended Data Fig. 5f–h**). L-Histidine, putrescine, and 7-methylguanosine were found at significantly higher levels in astrocytes than in microglia or neurons (**Extended Data Fig. 5i–k**). Fructose 1,6-bisphosphate, guanidinoacetate, and spermine were present at significantly higher levels in microglia than in neurons or astrocytes (**Extended Data Fig. 5l–n**). Choline, betaine, and L-valine were found at significantly higher levels in microglia and neurons than in astrocytes (**Extended Data Fig. 5o-q**).

### Comparison of untargeted metabolomes of microglia, astrocytes, and neurons

To systematically identify metabolites in various brain cells, we performed untargeted metabolomics in different cell samples. We detected 9,854 metabolite features for annotation with the mzCloud libraries and **[for]** categorization using the Human Metabolome Database (HMDB) (**Supplementary Fig. 2a and b**). Based on a PCA of the metabolite features, we found that neurons, astrocytes, and microglia were well separated (**Fig. 3a**). We also detected a substantial number of metabolites enriched in specific cell types (**Fig. 3a**), such as palmiticamide, malate, and sphingosine (d17:0) in astrocytes; trigonelline, lysophosphatidylserine, and phosphorylcholine in microglia; and creatine and D-pipecolic acid in neurons (**Fig. 3b**). For each metabolite, we compared its abundance in a given cell type to its average abundance in the other two cell types. Metabolites were considered enriched if they met the criteria of fold change > 2 and *P* < 0.01. Using these thresholds, we identified 250 metabolite features enriched in microglia, 856 in astrocytes, and 495 in neurons. A more stringent threshold (fold change > 10, *P* < 0.01) yielded 24 metabolites enriched in microglia, 286 in astrocytes, and 40 in neurons (**Supplementary Figs. 3a, 3b**).

**Fig. 3.**
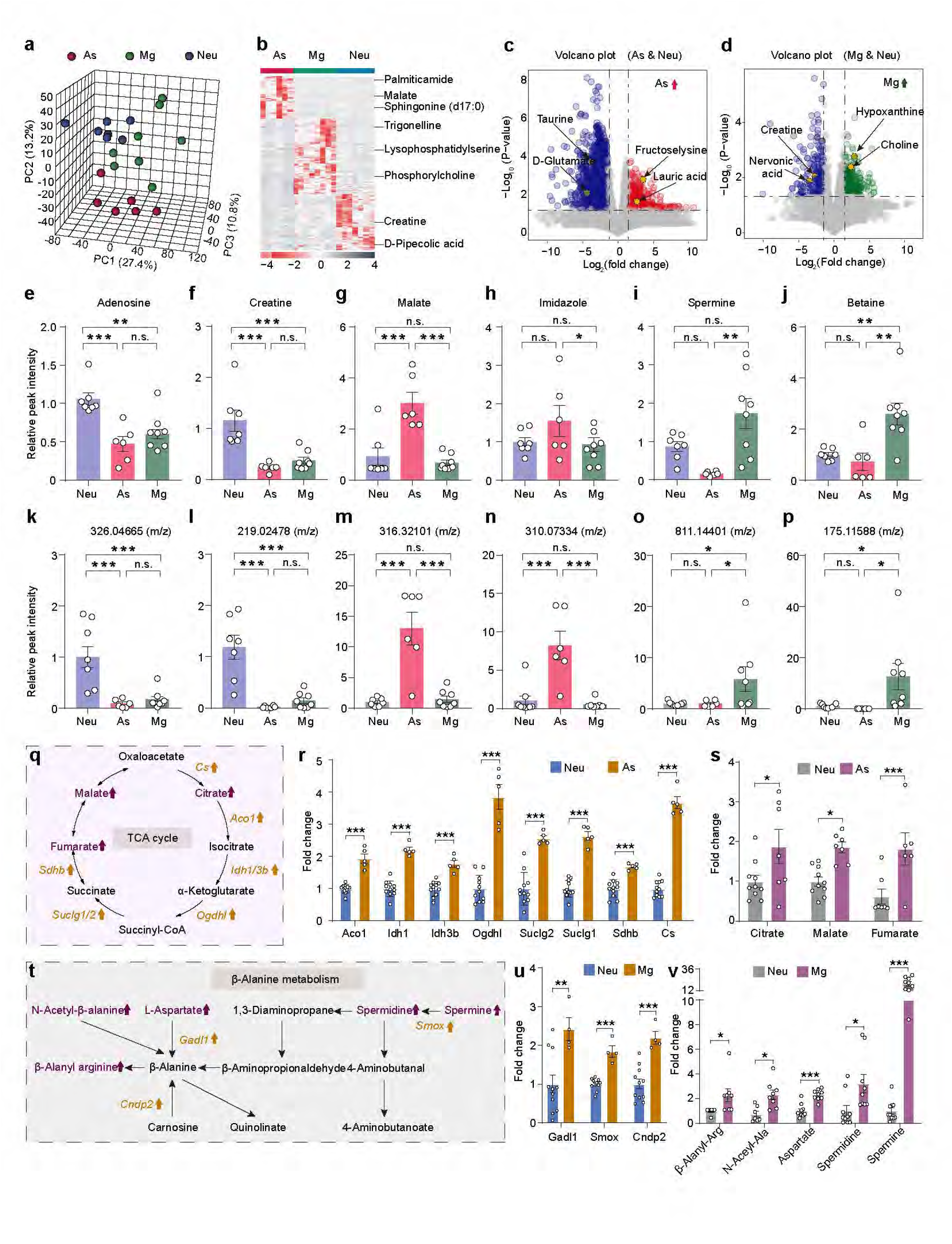
The metabolomes of neurons, astrocytes, and microglia based on untargeted metabolomics. **(a)** PCA of the metabolomes of neurons (Neu), astrocytes (As), and microglia (Mg) obtained through untargeted metabolomics analysis. In total, 12,166 features were detected from each sample and used for PCA. **(b)** Heatmap representation of top metabolites of the three groups, as identified by fold change, *P*-value, and abundance. Each column represents a sample. The red color indicates the greater abundance of a metabolite. **(c, d)** Volcano plot of differential metabolites between astrocytes and neurons (c), or between microglia and neurons (d). Differential (red and blue) and non-differential (gray) metabolites were defined by the criteria of fold change > 2 and *P* < 0.05 (two-tailed unpaired *t*-test). **(e–p)** Relative peak intensities of representative identified metabolites among neurons, astrocytes, and microglia, including adenosine (e), creatine (f), malate (g), imidazole (h), spermine (i), and betaine (j), 326.04665 (*m/z*) (k), 219.02478 (*m/z*) (l), 316.32101 (*m/z*) (m), 310.07334 (*m/z*) (n), 811.14401 (*m/z*) (o), and 175.11588 (*m/z*) (p). **(q)** The expression of genes and the level of metabolites in the TCA cycle pathway in neurons and astrocytes. Genes in yellow and metabolites in purple were expressed at a higher level in astrocytes than in neurons. **(r, s)** Histogram showing a comparison of the expression of TCA cycle–related genes (r) and the abundance of metabolites from the TCA cycle (s) in astrocytes and neurons. **(t)** A portion of the β-Alanine metabolism pathway in microglia and neurons. Genes in yellow and metabolites in purple were expressed at a high level in microglia when compared with neurons. **(u, v)** Comparisons of the genes expression (u) and the abundance of metabolites (v) in the β-Alanine metabolism pathway in microglia and neurons. Data are represented as the mean ± SEM; \**P* < 0.05, \*\**P* < 0.01, \*\*\**P* < 0.001. One-way ANOVA. In (a, b, e–p, s, v): As, n = 6 samples; Mg, n = 8 samples; Neu, n = 7 samples. In (r, u): As, n = 5 samples; Mg, n = 4 samples; Neu, n =11 samples.

However, although over 5,000 metabolite features have been found in brain tissues, only a few have been investigated ^38,39,64^. To systematically characterize metabolites that might be involved in the unique function of a particular brain cell type, we compared metabolic profiles obtained from non-targeted metabolomic measurements between astrocytes and neurons and between microglia and neurons. Of the metabolites we detected, taurine and glutamate showed high levels in neurons (fold change > 4, *P* < 0.05). In addition to 7-methylguanosine and putrescine, which we detected as being enriched in astrocytes through targeted metabolomics, we also found other metabolites, including fructoselysine and lauric acid, with higher abundance in astrocytes when compared with neurons (**Fig. 3c**). These metabolites have been reported to stimulate ketone body production in astrocytes and then promote neuronal maturation ^65,66^. Compared with microglia, the levels of creatine and nervonic acid were significantly higher in neurons (**Fig. 3d**). In contrast, hypoxanthine and choline levels were higher in microglia (**Fig. 3d**). This result is supported by a previous study in which nervonic acid was found to be required to synthesize many sphingolipids and gangliosides in neurons ^67^. In addition, creatine is abundant in neurons. It may be involved in regulating cellular energy and replenishing cellular ATP without oxygen ^68^.

We further compared our results from non-targeted and targeted metabolomics. Representative metabolites including adenosine, creatine, imidazole, spermine, and betaine in the three studied brain cell types showed similar cell type–specific tendencies across both analyses (**Figs. 3e-j**, also see **Extended Data Fig. 5**), suggesting that the untargeted metabolic measurements were reliable. In addition, some metabolites that had not yet been annotated were highly enriched in a certain cell type (**Fig. 3k-p**), such as 326.04665 (*m/z*) and 219.02478 (*m/z*) in neurons, 316.32101 (*m/z*) and 310.07334 (*m/z*) in astrocytes, and 811.14401 (*m/z*) and 175.11588 (*m/z*) in microglia. Given the challenges in performing verification and quantification of a large number of unknown metabolites through non-targeted measurements, more experiments are required for their annotation in the future.

Of the metabolites we detected in microglia and astrocytes, sphingosine (d18:1), 9-oxononanoic acid, and 2-methylcholine showed higher levels in microglia than in astrocytes (fold change > 1.5, *P* adj < 0.05) (**Supplementary Fig. 4a-b**). Other metabolites, including adenine, malate, and carnosine, showed lower abundance in microglia than in astrocytes (fold change > 1.5, *P* adj < 0.05; **Supplementary Fig. 4b**). With a Kyoto Encyclopedia of Genes and Genomes (KEGG) pathway analysis, we found that the Arginine and proline metabolism pathway and the Glycerophospholipid metabolism pathway showed the most substantial differences between microglia and astrocytes (**Supplementary Fig. 4c).** In the Arginine and proline metabolism pathway (**Supplementary Fig. 4d**), *Sms* and *Gatm* showed higher expression in microglia than in astrocytes **(Supplementary Fig. 4e**). Interestingly, levels of L-arginine, creatine, guanidinoacetate, *S*-adenosylmethionine, spermidine, and spermine of the Arginine and proline metabolism pathway were dramatically higher in microglia than in astrocytes (**Supplementary Fig. 4f**).

### Comparison of metabolism-related transcriptomes in microglia, astrocytes, and neurons

To better understand the uniqueness of cell metabolism at both metabolomic and transcriptomic levels in different brain cells, we collected neurons, astrocytes, and microglia from age-matched fluorescently labeled transgenic mice for bulk RNA-seq. Based on PCA and cluster analysis, the samples corresponding to a single cell type clustered together, whereas sample groups corresponding to neurons, astrocytes, and microglia were well separated (**Extended Data Fig. 6a-c**). The featured metabolic enzymes of the three types of brain cells were also easily detected, including *Sat1* in microglia, *Aldh1l1* and *Slc1a2* in astrocytes, and *Vglut1* in neurons, indicating the high quality of our RNA-seq results (**Extended Data Fig. 6d**).

Comparing the transcriptomes of astrocytes and neurons, we observed that the majority of the enriched metabolic genes in astrocytes encoded cellular components related to the mitochondrial inner membrane or matrix (**Extended Data Fig. 6e**). We further detected a notable difference in the expression of genes encoding TCA cycle enzymes between astrocytes and neurons (**Fig. 3q**). *Cs*, *Aco1*, *Idh1/3b*, *Ogdhl*, *Suclg1/2*, and *Sdhb*, which encode the key enzymes in the TCA cycle for metabolite synthesis, showed significant differences in their expression (**Fig. 3r**). High glycolytic activity and oxidative phosphorylation can be detected in astrocytes ^69–71^. Consistently, the critical components of the TCA cycle, namely citrate, fumarate, and malate, identified in our targeted and untargeted metabolomic analyses were more abundant in astrocytes as compared with neurons (**Fig. 3s**).

We then performed Gene Set Enrichment Analysis (GSEA) to compare gene expression between microglia and neurons. In addition to known differential gene sets in microglia, we observed that multiple metabolism-related gene sets, including Regulation of transport and Response to organic substance, were significantly more highly expressed in microglia than in neurons (**Extended Data Fig. 6f**). With KEGG pathway analysis, we found that *Gadl1*, *Cndp2*, and *Smox* in the β-Alanine metabolism pathway (**Fig. 3t**) showed higher expression in microglia than in neurons (**Fig. 3u**). Interestingly, levels of the precursors (*N*-acetyl-β-alanine, L-aspartate) and the product (β-alanyl-arginine) of the β-Alanine metabolism pathway were dramatically higher in microglia than in neurons (**Fig. 3v**). In the same pathway, the levels of spermine and spermidine were also higher in microglia than in neurons (**Fig. 3v**), indicating that the synthesis of spermidine from spermine was more active in microglia.

### Characterizing microglia-enriched metabolites *in vivo*

To determine to what extent the metabolites we found from sorted brain cells reflect the existence of metabolites *in vivo*, we established a microglial depletion model for metabolomic analysis using PLX3397, a selective inhibitor of colony-stimulating factor 1 receptor, CSF1R ^72,73^. We compared the metabolic profiles of cortical samples from control and microglial-depleted mice (**Fig. 4a-c)**. In total, 10,676 metabolite features were identified in both groups (**Fig. 4d**). Of these detected features, 387 metabolites were decreased in the PLX3397-treated group (fold change > 1.5, *P* < 0.05; **Fig. 4e, f**), which may represent unique metabolites enriched in microglia. The identities of these decreased metabolites were consistent with those of metabolites found in microglia through fluorescence-activated cell sorting (FACS)—for example, spermidine, γ-aminobutyric acid, GSH, hypoxanthine, *P*-guanidinoacetate, L-citrulline, and trigonelline (**Fig. 4g-m**). Additionally, some metabolites related to choline metabolism, e.g., choline, glycerophosphocholine (GPC), and betaine (**Figs. 4n-p**), decreased when microglia were depleted, which is consistent with the high levels of these metabolites detected in sorted microglia via FACS.

**Fig. 4.**
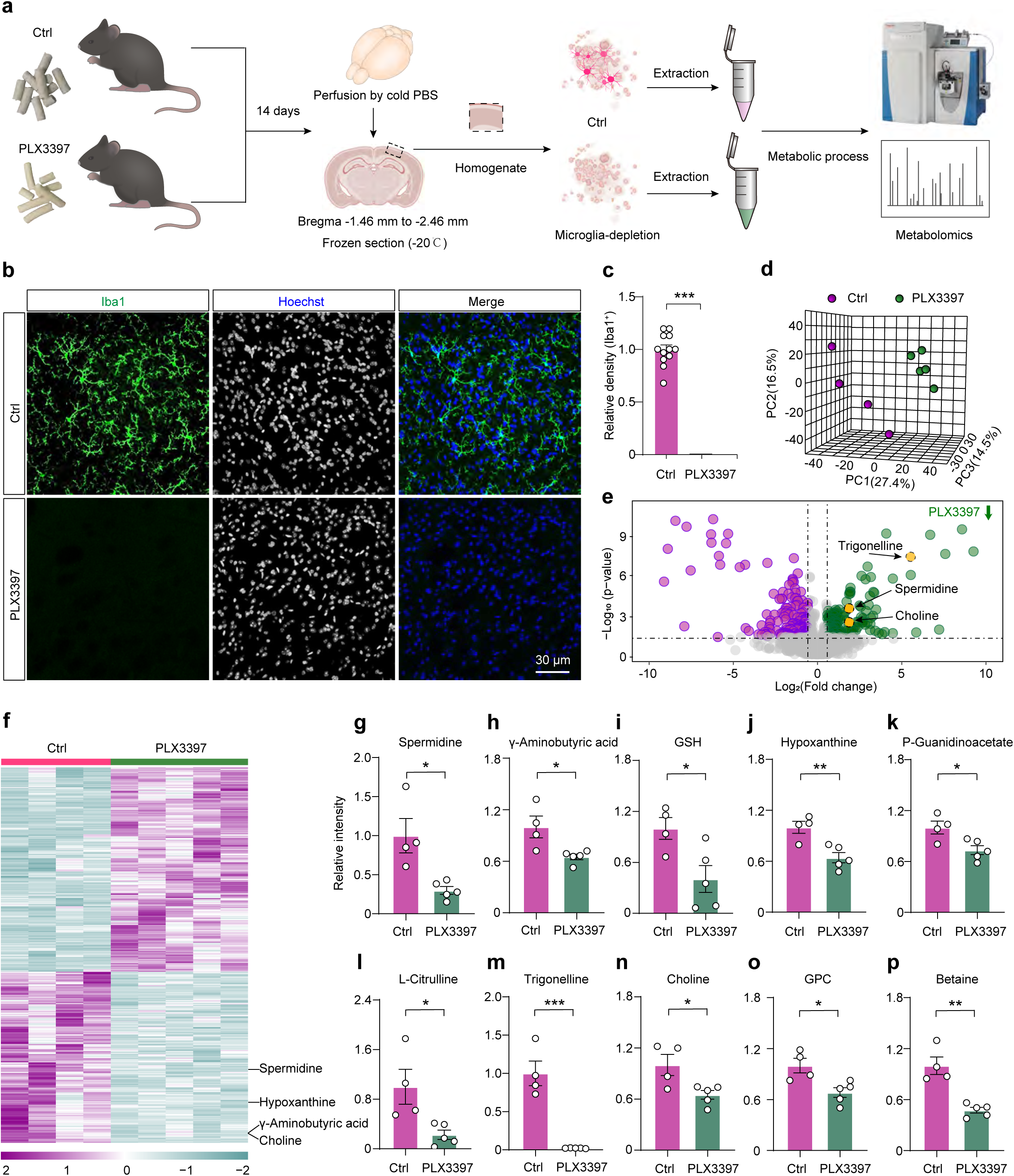
Characterizing microglia-specific metabolites *in vivo*. **(a)** Untargeted metabolomic analysis of brain tissues with microglial ablation in mice given food formulated with a CSF1-R blocker, PLX3397. Control mice (Ctrl) were given food without PLX3397. **(b)** Depletion of microglia with PLX3397 treatment. Representative images of the region in the cerebral cortex from control (upper panel) and PLX3397-treated (lower panel) mice. Microglia, anti-Iba1 (green); nuclei, Hoechst 33342 (Hoechst, blue). **(c)** Relative density of Iba1^+^ cells in the cerebral cortex from control and PLX3397-treated groups (*P* = 8.55e-17, n = 12 areas for each group, two-tailed unpaired *t*-test). **(d)** PCA of the metabolomes of the control and PLX3397-treated groups analyzed with untargeted metabolomics. In total, 10,676 features were detected from each sample and used for PCA. Circles indicate individual samples from each group. **(e)** Volcano plot of differential metabolites between the control and PLX3397-treated groups. Differential (purple and green) and non-differential (gray) metabolites were defined by the criteria of fold change >1.5 and *P* < 0.05, with *P*-values calculated using a two-tailed unpaired *t*-test. A data point with a negative fold change indicates a low level of a metabolite in the control group, whereas a positive value indicates a low level of a metabolite in the PLX3397-treated group. **(f)** Heatmap representation of top metabolites of the control group and PLX3397-treated group, as identified by fold change, *P*-values, and metabolite abundance. Each column represents a sample. The purple color indicates a greater abundance of a metabolite. **(g–p)** Relative peak intensity of representative metabolites between control and PLX5622-treated groups, including spermidine (g, *P* = 0.010), γ-aminobutyric acid (h, *P* = 0.021), GSH (i, *P* = 0.026), hypoxanthine (j, *P* = 0.006), P-guanidinoacetate (k, *P* = 0.021), L-citrulline (l, *P* = 0.021), trigonelline (m, *P* < 0.001), choline (n, *P* = 0.025), GPC (o, *P* = 0.015), betaine (p, *P* = 0.001). Data are represented as the mean ± SEM; \**P* < 0.05, \*\**P* < 0.01, \*\*\**P* < 0.001, two-tailed unpaired *t*-test. In (a, d, f–p): Ctrl, n = 4; PLX3397, n = 5.

To test whether there are potential off-target effects of PLX3397, we used a genetic approach, *Cx3cr1-CreER;DTA* to deplete microglia (**Extended Data Fig. 7a-d**). Metabolomic profiling of cortical tissues of *Cx3cr1-CreER;DTA* mice indicated metabolic alterations relative to controls, closely resembling those observed in PLX3397-treated mice. Specifically, levels of choline, betaine, GSH, L-citrulline, and γ-aminobutyric acid all showed decreasing trends after microglial ablation (**Extended Data Fig. 7e**). Among these, betaine, GSH, and L-citrulline were significantly reduced, whereas choline and γ-aminobutyric acid exhibited downward trends that did not reach statistical significance (**Extended Data Fig. 7f**).

### Characterizing changes in the microglial, astrocyte, and neuronal metabolomes during aging

With age, significant changes occur in brain metabolism accompanied by alterations in the functions of neurons and glial cells ^74,75^. Therefore, we carried out metabolomic profiling of aging microglia, astrocytes, and neurons. We isolated them from the cerebral cortex and hippocampus at different ages (1 month, 12 months, 18 months, and 30 months). Untargeted metabolomics and bulk RNA-seq were carried out on all samples for simultaneous metabolome and transcriptome analyses (**Fig. 5a**). In total, 13,700 metabolite features were detected, and 7,122 (51.9%) of them were annotated (**Fig. 5b**). On the basis of a PLS-DA model of the metabolite features, we found that the metabolomic profiles of microglia from mice at different ages were significantly separated (**Fig. 5c**). To better understand the quality of these changes over time, we clustered all measured metabolites based on their time trajectories and divided them into eight clusters (**Fig. 5d**). Cluster 7 and cluster 8 comprise metabolites with declining and enhancing trajectories over time, respectively. Some of the metabolites in these clusters were enriched in microglia, such as GSH and spermine (**Fig. 5e**). Of the metabolic pathways that were declining and enhancing over time, the glutathione metabolism and Aminoacyl-tRNA biosynthesis pathways were the most notable as they were also enriched in microglia (**Fig. 5f-g**, **Fig. 1i**). Levels of GSH and GSH/GSSG decreased significantly at 12 months, 18 months, and 30 months as compared with those at 1 month (**Fig. 5h-j**), which may directly lead to the gradual decline of the antioxidant capacity of microglia during aging. Other representative metabolites, such as diethanolamine, adenine, and uridine, decreased in cluster 7 (**Fig. 5k-m**), and CDP-ethanolamine, spermine, and histamine increased in cluster 8 (**Fig. 5n-p**). With a targeted metabolomic approach, we observed that GSH levels were higher in 1-month-old microglia compared with those from 12- and 30-month-old mice, with a significant difference between 1 and 12 months (**Supplementary Fig. 5a, b**).

**Fig. 5.**
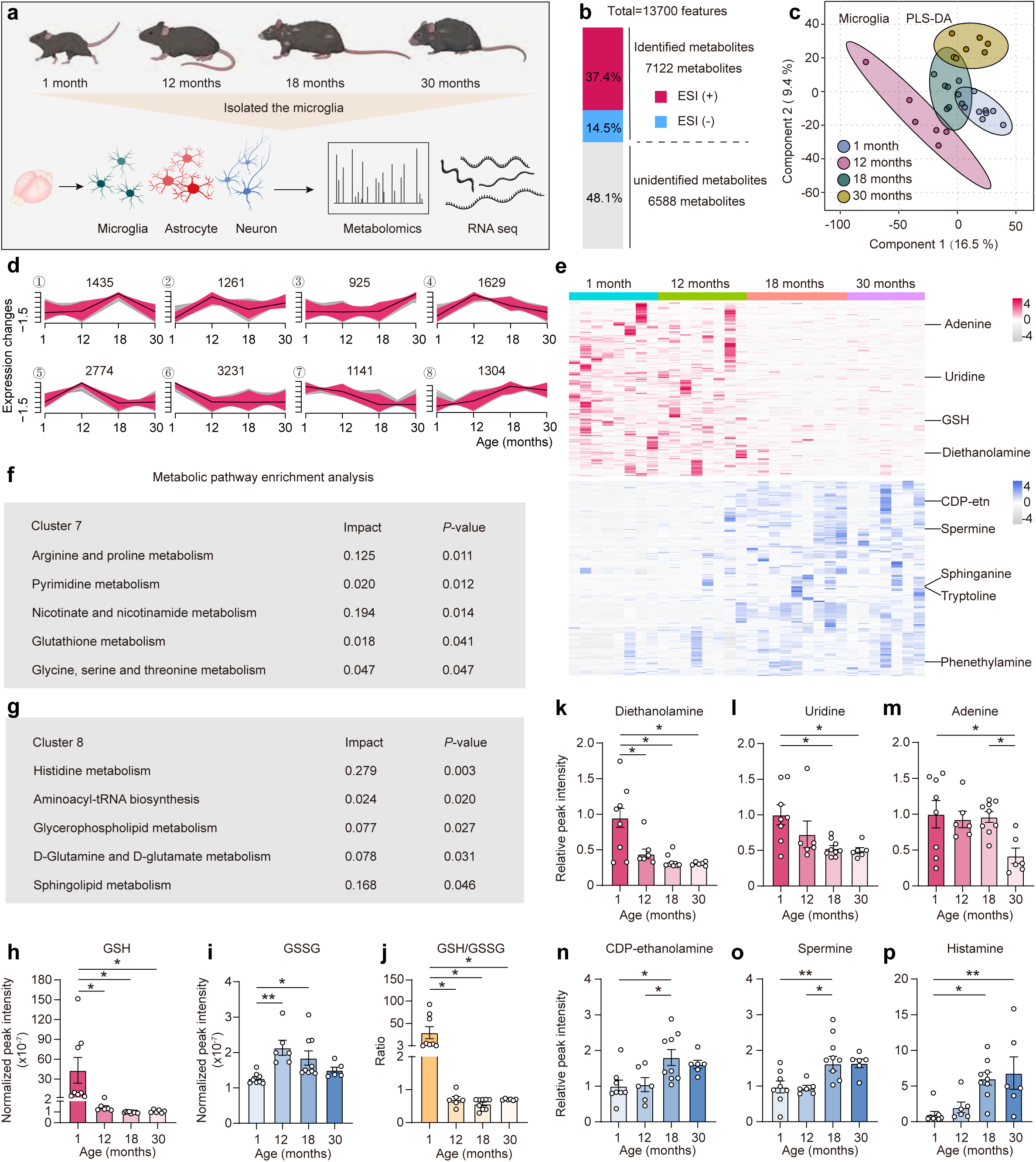
Characterizing the metabolome of microglia from aging brains. (a) Graphic illustration of the workflow to acquire the microglia metabolome from aging mouse brains. (b) Feature composition of the aging microglia metabolome using untargeted metabolomics. Relative percentages of metabolites identified by electrospray ionization positive (ESI+) and negative (ESI–) modes. (c) PLS-DA of metabolomes of aging microglia. In total, 13,700 features were identified from all samples used for PLS-DA. Circles indicate individual samples of microglia from different age groups (1, 12, 18, and 30 months). (d) Clustering of time course expression patterns in microglia from mouse brains using the fuzzy c-means algorithm. Warm (red) and cold (gray) colors indicate low and high deviation from the consensus profile, respectively. (e) Heatmap representation of top metabolites of the four groups from clusters 7 and 8, as identified by fold change, *P*-values (two-sided Student’s *t*-test), and abundance. Each column represents a sample. The red or blue color indicates greater abundance of a metabolite, whereas the gray color indicates lower abundance. (**f, g**) Top five metabolic pathways analyzed from KEGG. The analyzed metabolites were from cluster 7 (f) and cluster 8 (g) in (e). KEGG, Kyoto Encyclopedia of Genes and Genomes. Pathway enrichment was performed using a hypergeometric test and pathway topology analysis based on relative-betweenness centrality. (**h–j**) Relative peak intensity of GSH (h), GSSG (i), and their relative ratio (j) in microglia from mouse brains of different ages. GSH, glutathione (reduced); GSSG, (oxidized). (**k–p**) Relative peak intensity of diethanolamine (k, cluster 7), uridine (l, cluster 7), adenine (m, cluster 7), CDP-ethanolamine (n, cluster 8), spermine (o, cluster 8), and histamine (p, cluster 8) in microglia from mouse brains of different ages. Data are represented as the mean ± SEM; one-way ANOVA; \**P* < 0.05, \*\**P* < 0.01. ANOVA, analysis of variance. In (c, e, h–p): 1 month, n = 8 samples; 12 months, n = 6 samples; 18 months, n = 9 samples; 30 months, n = 6 samples.

For astrocytes, we used *AAV2/PHP.eB-gfaABC1D-mRuby3-WPRE-pA* to label and isolate astrocytes from the brains of mice at different ages (**Supplementary Fig. 6a-i**). The metabolomic profiles of astrocytes from these mice were **notably** separated, and many metabolites changed over time (**Fig. 6a, b**). Clusters 7 and 8 comprised metabolites (e.g., malate) with declining and enhancing trajectories over time, respectively (**Fig. 6c**). Multiple pathways from clusters 7 and 8 underwent **decreases and increases in pathway enrichment** over time (**Fig. 6d, e**). In addition, PE(O-18:0/14:0), CDP-ethanolamine, pyridoxal, and pyridoxal 5’-phosphate decreased in cluster 7 (**Fig. 6f-i**), whereas hexanoic acid, isobutyrate, nicotinamide, 2-oxoglutaric acid, and malate increased in cluster 8 (**Fig. 6j-n**).

**Fig. 6.**
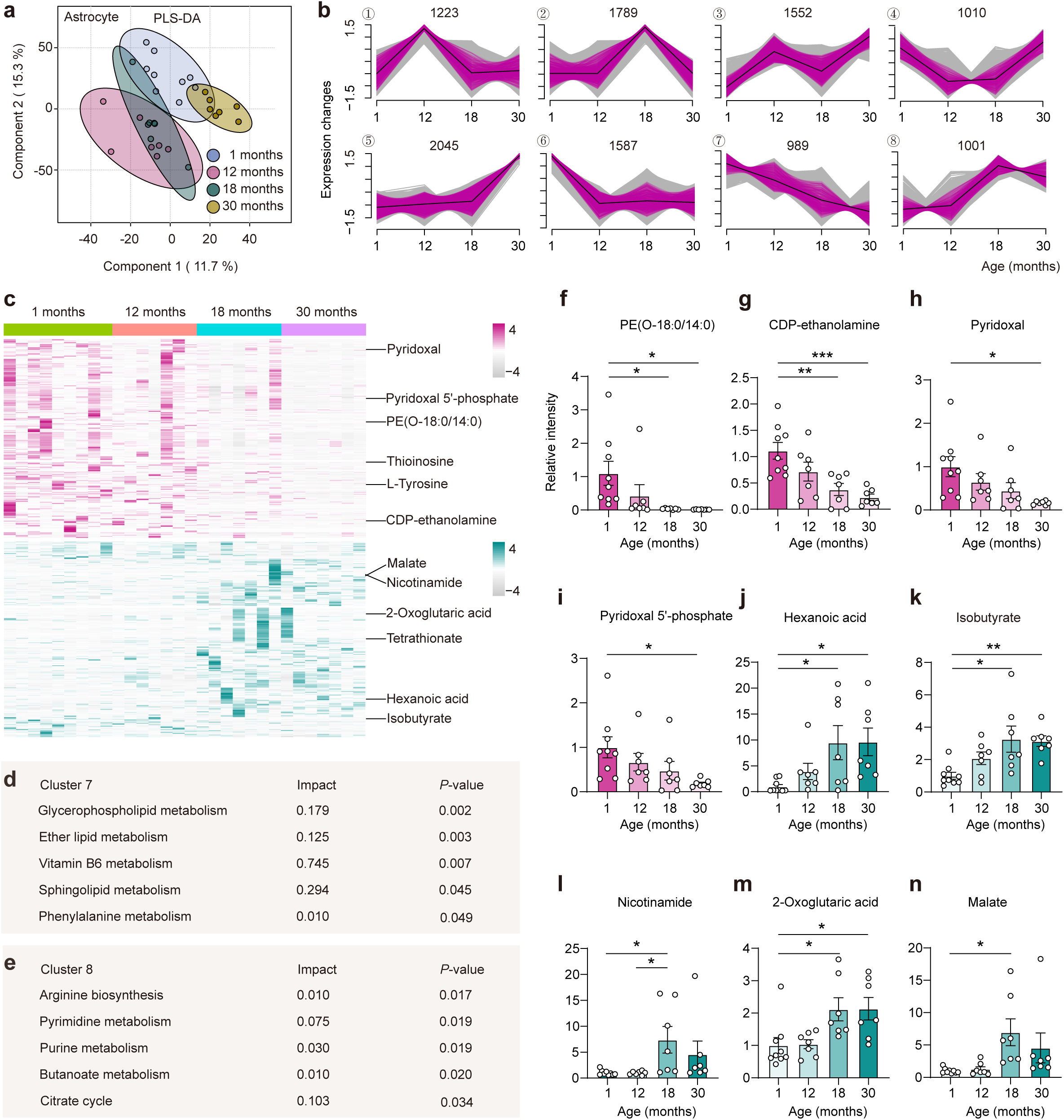
Characterizing the metabolome of astrocytes from aging brains. (a) PLS-DA of metabolomes of aging astrocytes. Two components explained 27.0% (11.7% and 15.3%) of the variance among the four groups. Component scores are indicated as percentages; circles indicate individual samples of astrocytes from different age groups (1, 12, 18, and 30 months). (b) Clustering of time course expression patterns in astrocytes from mouse brains using the fuzzy c-means algorithm. Warm (purple) and cold (gray) colors indicate low and high deviation from the consensus profile, respectively. (c) Heatmap representation of top metabolites of the four groups from clusters 7 and 8, as identified by fold change, *P*-values (two-tailed unpaired *t*-test), and abundance. Each column represents a sample. Red or blue color indicates greater abundance of a metabolite, whereas gray indicates lower abundance. (d) Top five metabolic pathways based on KEGG. The analyzed metabolites were from cluster 7 in (c). (e) Top five metabolic pathways based on KEGG. The analyzed metabolites were from cluster 8 in (c). Pathway enrichment in (d) and (e) was performed using a hypergeometric test and pathway topology analysis based on relative-betweenness centrality. (**f–n**) Relative peak intensity of PE(O-18:0/14:0) (f, cluster 7), CDP-ethanolamine (g, cluster 7), pyridoxal (h, cluster 7), pyridoxal 5’-phosphate (i, cluster 7), hexanoic acid (j, cluster 8), isobutyrate (k, cluster 8), nicotinamide (l, cluster 8), 2-oxoglutaric acid (m, cluster 8) and malate (n, cluster 8) in astrocytes from mouse brains of different ages. Data are represented as the mean ± SEM; one-way ANOVA; \**P* < 0.05, \*\**P* < 0.01, \*\*\**P* < 0.001. ANOVA, analysis of variance. In (a, c, f–n): 1 month, n = 9 samples; 12 months, n = 7 samples; 18 months, n = 7 samples; 30 months, n = 7 samples.

We also found that the metabolomic profiles of neurons from mice at different ages were **notably** separated (**Extended Data Fig. 8a**). Using the fuzzy c-means method ^76^, we clustered all measured metabolites based on their temporal trajectories and divided them into eight clusters (**Extended Data Fig. 8b**). Some of the metabolites in clusters 7 and 8 were enriched in neurons, such as taurine (**Extended Data Fig. 8c**). The top five metabolic pathways from these two clusters are also shown (**Extended Data Fig. 8d, e**). Other representative metabolites, such as SL 20:2;O/15:0, carnosine, taurine, and erucic acid, decreased in cluster 7 (**Extended Data Fig. 8f-i**), whereas cytidine 5’-*P*, crotonic acid, L-aspartic acid, 2’-deoxymugineicacid, and methylacetate increased in cluster 8 (**Extended Data Fig. 8j-n**).

### Characterizing the metabolic alterations of the microglial, astrocyte, and neuronal metabolomes in a mouse model of AD

Little is known about metabolic alterations of brain cells during AD. To determine the involvement of metabolic reprogramming of microglia, astrocytes, and neurons during AD progression, we carried out metabolomic profiling of these three brain cell types in 5xFAD mice (**Fig. 7a**), which express mutant human *PSEN1* and *APP* transgenes and show AD pathology^77^. Based on PLS-DA and clustering analysis, the metabolomes from 5xFAD mice and wild type were well separated into two distinct clusters (**Fig. 7b, 7c**). We detected a substantial number of metabolites that were down-regulated in 5xFAD mice as compared to wild-type mice (**Fig. 7d**). Notably, GSH, the microglia-enriched metabolite, which progressively decreased in abundance during aging, was lower in the 5xFAD group (**Fig. 7e**). Although there was no difference in GSSG between the 5xFAD group and wild-type group, the GSH/GSSG ratio was significantly reduced in the 5xFAD group (**Fig. 7f, 7g**). Spermine, which progressively increased in abundance during aging, was at lower levels in the 5xFAD group (**Fig. 7h**). Other metabolites, such as NADP^+^, adenine, guanine, and adenosine diphosphate (ADP), also decreased in the 5xFAD group (**Fig. 7i-l**). In contrast, some metabolites, including guanidinoacetate, indole, lysidine, and putrescine, showed higher abundance in the 5xFAD group (**Fig. 7m-p**). To identify the metabolic pathways that were altered in microglia from the 5xFAD group, we carried out a global metabolic pathway analysis using identified metabolites (fold change > 1.2, *P* < 0.01). The glutathione metabolic pathway of microglia was significantly downregulated in association with AD pathology (**Fig. 7q**). We further found that most metabolic enzymes in the glutathione metabolic pathway, such as *Chac1*, *Gstm5*, *Gss*, *Ggt5*, *Ggt7*, *Gclc*, and *Ggct*, were also significantly downregulated in microglia from the 5xFAD group (**Supplementary Fig. 7**). Consistent with our untargeted findings, GSH levels in 5xFAD microglia showed a decreasing trend relative to controls, although the difference did not reach statistical significance (*P* = 0.080), likely due to variability within groups (**Supplementary Fig. 8a, b)**. Together, these targeted analyses provide independent support for our conclusion that microglial GSH metabolism declines with aging and is impaired in AD pathology.

**Fig. 7.**
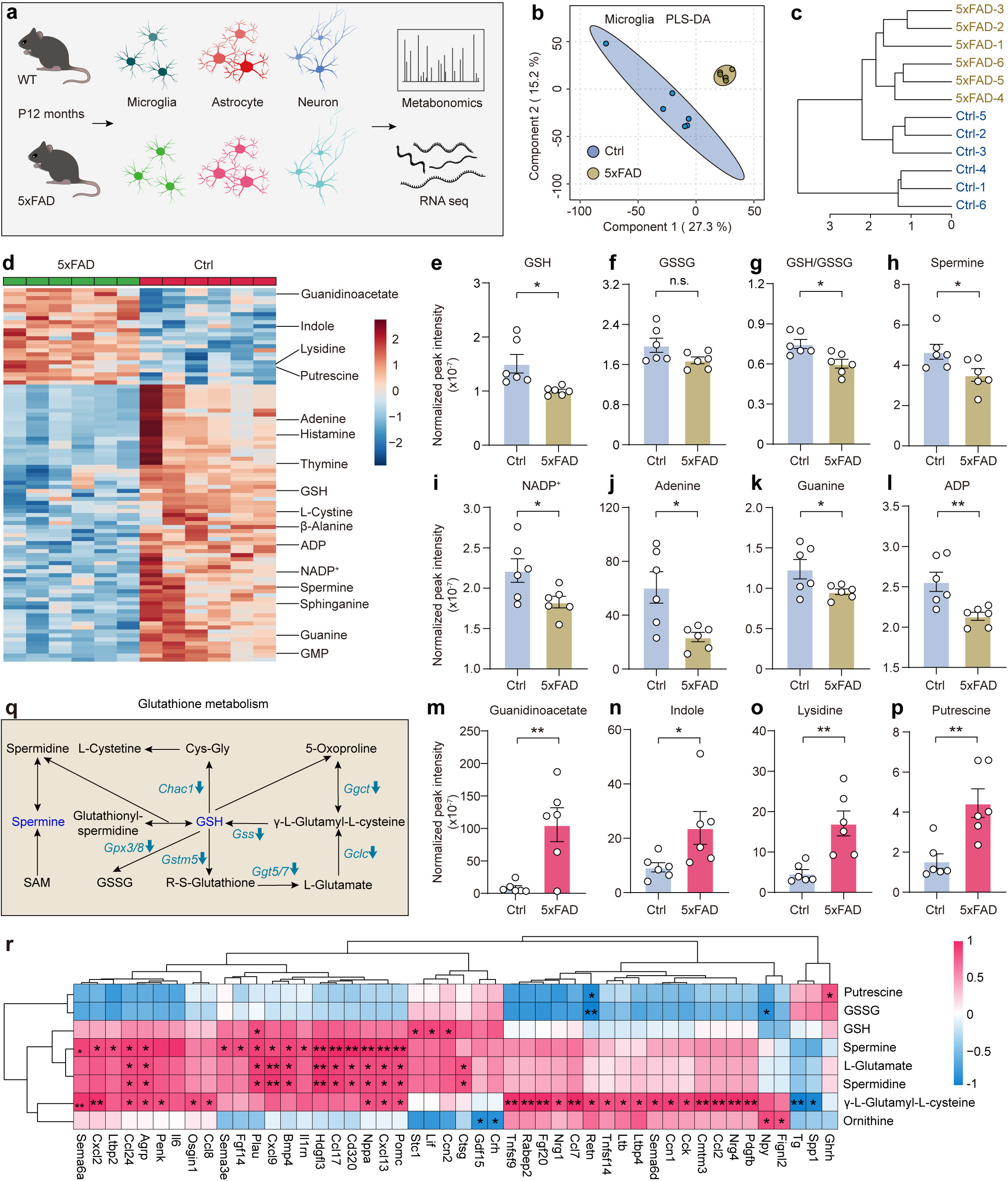
Characterizing the microglial metabolome in the brain of AD mouse models. **(a)** The workflow to acquire the metabolomes of microglia from 5xFAD and control (wild-type) mice. **(b)** PLS-DA of microglia metabolomes of mouse brains. In total, 13,700 features were identified from all samples. Two components explained 42.5% (27.3% and 15.2%) of the variance between the groups. Circles indicate individual samples of microglia from different groups. **(c)** Dendrogram showing the clustering of all microglial samples. **(d)** Heatmap representation of top metabolites (93 metabolites) in the two groups, as identified by VIP scores from a PLS-DA. Each column represents a sample from a different mouse. The red color indicates a greater abundance of a metabolite. **(e–g)** Relative peak intensity of GSH and GSSG of microglia between the 5xFAD and control group: GSH (e, *P* = 0.016), GSSG (f, *P* = 0.082). The ratio of GSH to GSSG (g, *P* = 0.010). **(h–p)** Relative peak intensities of representative metabolites from microglia between the 5xFAD group and control group, including spermine (h, *P* = 0.038), NADP^+^ (i, *P* = 0.037), adenine (j, *P* = 0.013), guanine (k, *P* = 0.048), ADP (l, *P* = 0.008), guanidinoacetate (m, *P* = 0.004), indole (n, *P* = 0.043), lysidine (o, *P* = 0.003), and putrescine (p, *P* = 0.005). NADP, nicotinamide adenine dinucleotide phosphate; ADP, adenosine diphosphate. **(q)** A portion of the glutathione metabolism pathway of microglia between the 5xFAD group and control group. Metabolites and genes in blue were present at lower levels in the microglia of the 5xFAD group compared with the control group. **(r)** A heatmap of Spearman’s rank correlation analysis between the identified metabolites in glutathione metabolism pathway and the cytokine-related gene expression in microglia between the 5xFAD group and control group. Red indicates positive correlation and blue denotes negative correlation. Data are presented as the mean ± SEM; two-tailed unpaired *t*-test *P*-values; \**P* < 0.05, \*\**P* < 0.01; n.s., no significant difference. Ctrl, control. In (a–p): Ctrl, n = 6 samples; 5xFAD, n = 6 samples.

As immune cells of the brain, microglia play a protective role by secreting a range of cytokines, facilitating the clearance of deposited Aβ through processes such as chemotaxis and phagocytosis^78^. Therefore, we conducted a combined correlation analysis using data from both the 5xFAD group and wild-type group to examine the correlation between metabolites associated with GSH metabolism and polyamine metabolism in microglia, alongside a subset of cytokines identified in the transcriptomic analysis (**Fig. 7r**). These subgroup analyses indicated distinct correlation patterns between the control and 5xFAD mice, underscoring the influence of the 5xFAD model on the interaction between metabolic and inflammatory pathways (**Supplementary Fig. 9a, b**). Notably, the levels of GSH, spermine, and spermidine were significantly positively correlated with the expression of some chemotaxis-related genes, such as *Cxcl2, Cxcl9, Ccl7, Ccl8,* and *Ccl24*^79,80^. Conversely, an inverse relationship was observed with the levels of putrescine and GSSG. This suggested that GSH and polyamine metabolism may play a crucial role in regulating the chemotactic processes of microglia.

Based on PLS-DA and clustering analysis, the metabolomes of both astrocytes and neurons from 5xFAD and wild-type mice were well separated into two distinct clusters (**Extended Data Fig. 9 and 10**). We detected a substantial number of metabolites that were down-regulated or up-regulated in neurons or astrocytes in 5xFAD mice compared to wild-type mice (**Extended Data Fig. 9 and 10**). Notably, malate, glutamine, cytidine, and 4-acetmidobutanoate were higher in astrocytes of the 5xFAD group relative to the control (**Extended Data Fig. 9d**), whereas tyrosocholic acid, sphingosine (d18:1), *O*-adipoylcarnitine, and hexanedioic acid were at lower levels in astrocytes of the 5xFAD group (**Extended Data Fig. 9e)**.

Oxamate, indole, L-2-amino-3-oxobutanoic acid, and hexyl isothiocyanate were higher in neurons of the 5xFAD group relative to the control (**Extended Data Fig. 10d**), whereas PC 18:1_18:1, arachidonic acid, 6-methyiflavone, and β-alanine were at lower levels in neurons of the 5xFAD group (**Extended Data Fig.10e**). To identify the metabolic pathways that were altered in neurons or astrocytes from the 5xFAD group, we carried out a global metabolic pathway analysis using identified metabolites (fold change > 1, *P* < 0.01). The arginine and proline metabolism pathway of astrocytes was significantly downregulated in mice with AD pathology relative to the normally aging mice (**Extended Data Fig. 9f).** The β-alanine metabolism pathway of neurons was significantly downregulated in mice with AD pathology (**Extended Data Fig. 10f**).

### Contribution of GSH and polyamine metabolism to microglial morphogenesis and Aβ deposition

Because GSH is converted to GSSG by enzymes encoded by the glutathione peroxidase (*Gpx*) family ^81,82^, we looked for differences in the expression of *Gpx* family members between the two groups. The expression of *Gpx3* and *Gpx8* in microglia was significantly decreased, whereas that of *Gpx1* and *Gpx4* was increased in the 5xFAD group relative to wild type (**Supplementary Fig. 10**), which may compensate for increased GSH consumption in the 5xFAD group. *Gpx1* is the most highly expressed enzyme in the brain. It converts GSH to GSSG (**Fig. 8a**). From our transcriptome results, it was highly enriched in microglia (**Fig. 8b**). To determine the contribution of the GSH and polyamine metabolism to microglia, we generated a transgenic mouse with global *Gpx1* knockout and investigated alterations in the metabolome of its microglia (**Fig. 8c**). The metabolome of microglia from the *Gpx*^−/–^ mice was well separated from that of control mice according to a PLS-DA analysis (**Fig. 8d**). *Gpx1* knockout led to the disruption of the GSH metabolic pathway in microglia of *Gpx1^−/–^* mice, which was characterized by a significant decrease in GSH, L-glutamate, spermine, and spermidine and a significant increase in GSSG, putrescine, and γ-L-glutamyl-L-cysteine (**Fig. 8e-g**).

**Fig. 8.**
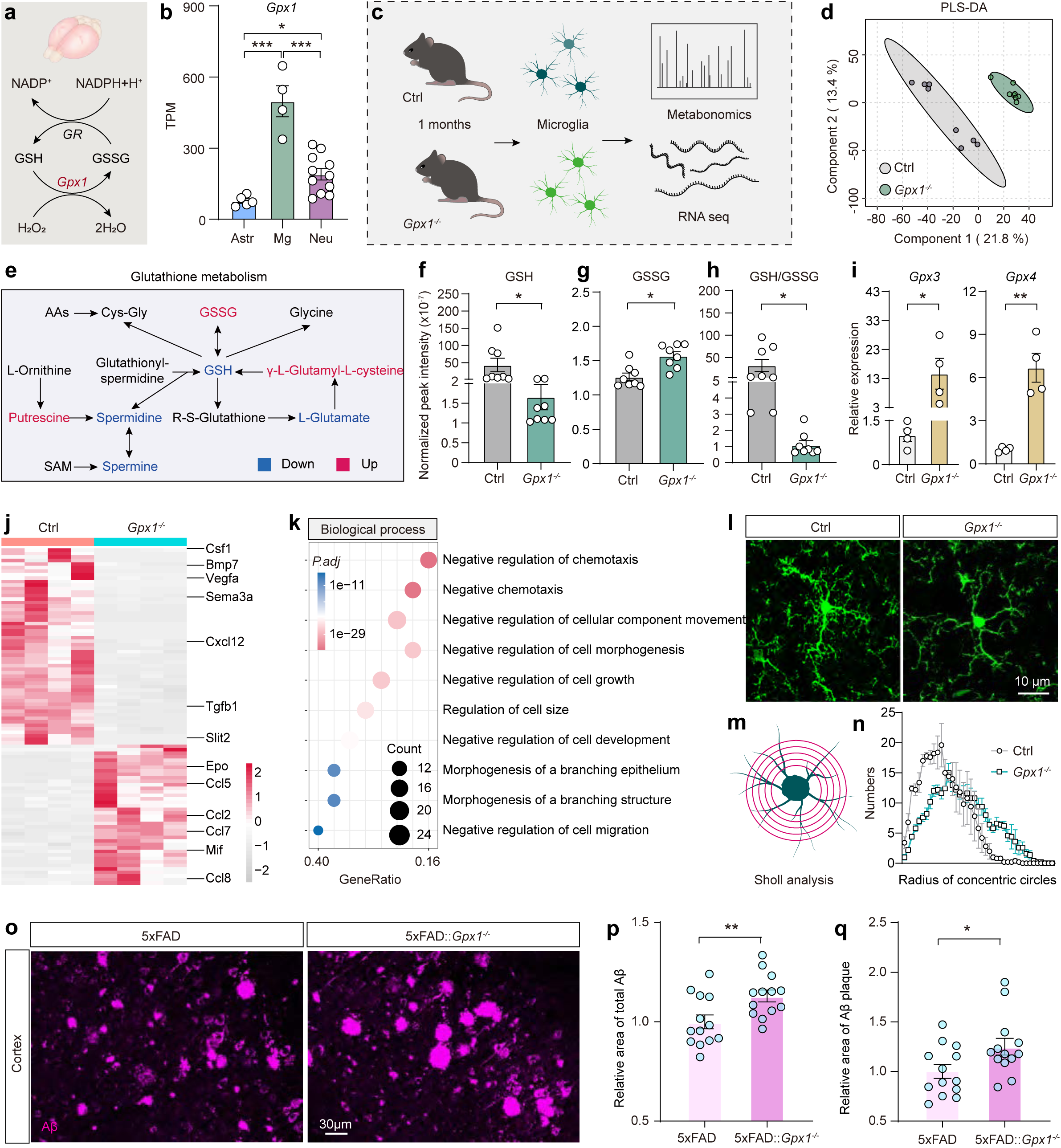
Disruption of glutathione metabolism and polyamine metabolism accelerates Aβ deposition in 5xFAD mice. **(a)** Mutual conversion of GSH and GSSG. GR, glutathione reductase. **(b)** *Gpx1* expression among neurons (Neu, n = 11), astrocytes (Astr, n = 5), and microglia (Mg, n = 4; astrocyte vs microglia, *P* < 0.001; astrocyte vs neuron, *P* = 0.039; neuron vs microglia, *P* < 0.001). TPM: transcripts per kilobase million. **(c)** The workflow to acquire the microglial metabolome. **(d)** PLS-DA of microglia metabolomes of control and *Gpx1*^−/–^ mouse brains (n = 8 samples each). **(e)** A portion of the glutathione metabolism pathway in microglia between the *Gpx1*^−/–^ group and control group. Metabolites in blue (or red) were present at lower (or higher) levels in microglia in the *Gpx1*^−/–^group. **(f–h)** Relative peak intensity of GSH and GSSG in control and *Gpx1*^−/–^ microglia (n = 8 samples each): GSH (f, *P* = 0.048), GSSG (g, *P* = 0.021). The ratio of GSH to GSSG (h, *P* = 0.034). **(i)** Comparisons of *Gpx3* and *Gpx4* expression in control and *Gpx1*^−/–^ microglia (*Gpx3*, *P* = 0.042; *Gpx4*, *P* = 0.001; n = 4 samples each). **(j)** Heatmap representation of cytokine-related gene expression in control and *Gpx1*^−/–^ microglia (n = 4 samples each). The red (or gray) color indicates a greater (or lower) abundance. **(k)** Gene Ontology (GO) analysis of differentially expressed cytokine-related genes in microglia (dot, the representative 10 enriched GO terms within ‘biological process’). Dot size, gene counts; dot color, significance of enrichment. **(l)** Images of microglia (green, anti-Iba1) in the cerebral cortex of a *Gpx1*^−/–^ and a control mouse. **(m, n)** Sholl analysis of reconstructed microglia in the *Gpx1*^−/–^ (n = 5 slices) and control (n = 5 slices) groups. **(o)** Image of Aβ in the cerebral cortex of a 5xFAD and a 5xFAD;*Gpx1*^−/–^ mouse. **(p, q)** Relative area of total Aβ (p) and Aβ plaque (q) (n = 13 for each group, *P* = 0.009). Data are represented as the mean ± SEM; \**P* < 0.05, \*\**P* < 0.01, \*\*\**P <* 0.001; n.s., no significant difference. In (b), one-way ANOVA; in (f–i, p, q), two-tailed unpaired *t*-test.

We also observed that the ratio of GSH to GSSG dramatically decreased in the *Gpx1*^−/–^ group (**Fig. 8h**), indicating that a substantial amount of the remaining GSH was converted to GSSG. Therefore, we looked for differences in other *Gpx* family members between these two groups. We noted a compensatory increase in *Gpx3* and *Gpx4* expression in the *Gpx1*^−/–^ group as compared to the control group (**Fig. 8i**). Further analysis of the transcriptomes of the *Gpx1*^−/–^ and control mice revealed a significant decrease in the expression of chemotaxis-related genes, such *Csf1*, *Bmp7*, *Vegfa*, *Sema3a*, *Cxcl12*, *Tgfb1*, and *Slit2*, in the *Gpx1*^−/–^ group (**Fig. 8j**). The majority of enriched genes in microglia of *Gpx1*^−/–^ mice encoded biological processes related to regulation of chemotaxis, cell morphogenesis, and cell migration (**Fig. 8k**), indicating that GSH and polyamine metabolism may influence the morphology and function of microglia through the regulation of associated cytokines.

To further verify this, we then stained brain sections with anti-Iba1. The microglia in the *Gpx1*^−/–^ group were abnormal, with fewer spines along their branches, and the total length of their branches was also substantially decreased (**Fig. 8l-n**), which is consistent with our analysis of the transcriptomes (**Fig. 8k**). During the progression of AD, microglia are able to clear extracellular deposits of Aβ ^83,84^. To further elucidate the impact of GSH and polyamine metabolism on microglial function, as well as their roles in the pathological progression of AD, we compared the amount of Aβ between the 5xFAD;*Gpx1*^−/–^ mice and the 5xFAD mice at 3 months after birth (**Fig. 8o**). The relative area of Aβ plaque in the brains of 5xFAD;*Gpx1*^−/–^ mice was significantly higher than that in the 5xFAD group (**Fig. 8p-q**). These results suggest that disruption of the GSH and polyamine metabolic pathway in microglia decreases their ability to clear Aβ and accelerates the progression of AD.

## Discussion

We systematically identified the metabolites and the corresponding metabolic enzymes that were enriched in three different brain cell types. A similar analysis was applied to these brain cells during aging and in a mouse model of AD. In particular, GSH, a typical enriched metabolite in microglia, plays a dominant role in the regulation of energy metabolism in microglia, and the disruption of this metabolism exacerbates the pathological progression of AD.

The investigation of different brain cell metabolomes is critical for understanding the cellular properties and functions of various cell types in the brain. For example, choline, an essential nutrient for humans and other mammals ^85^, is mainly used to synthesize acetylcholine, membrane phospholipids, and betaine ^86^. It is found at high levels in neurons. Unexpectedly, we observed that choline and its downstream metabolites, e.g., acetylcholine, betaine, and glycerophosphocholine, were also highly enriched in microglia. Other metabolites enriched in microglia, such as spermidine and spermine, the two ubiquitous biogenic amines, had dramatically higher levels in microglia than in neurons and astrocytes. In our RNA-seq results, *Smox*, *Sat1*, and *Paox*, the genes responsible for the metabolic pathway that connects spermine and spermidine, were expressed at higher levels in microglia, suggesting that the primary source of polyamines in the brain is their production from the metabolism of spermine in microglia. Moreover, several studies have implicated spermine and spermidine as inhibitors of microglial activation that thus prevent the production of pro-inflammatory mediators (NO and PGE2) and cytokines (TNF-α and IL-6)^87–89^. We did not observe glutamate enrichment in astrocytes. This may be attributed to the highly dynamic nature of astrocytic glutamate uptake, recycling, and interaction with neurons, which may not be fully captured in a steady-state metabolomic analysis.

The exchange of metabolites contributes to energy metabolism between astrocytes and neurons ^2^. For example, mitochondrial reactive oxygen species (ROS), lactate, and L-serine generated from astrocytes are shuttled to neurons to maintain neuronal redox status and support their energy needs ^90^. Likewise, lactate and pyruvate can be released from astrocytes and taken up by neurons to maintain their energy metabolism ^46,91^. Our study found that metabolites and related metabolic enzymes of the TCA cycle are present at a higher level in astrocytes than in neurons. This may be due to their high glycolytic activity or the large amount of acetyl-CoA produced from fatty acid β-oxidation in astrocytes ^92^. In summary, the regulation of energy metabolism in astrocytes and neurons mainly depends on glycolysis and related downstream metabolites. However, these metabolic regulatory patterns do not dominate energy metabolism in microglia.

We detected many metabolites enriched in microglia, with GSH and polyamine metabolism being the prominent examples. In the physiological state, GSH can be converted to GSSG and can be chelated with iron ions or copper ions ^93,94^; GSH can also enter mitochondria and be converted to glutamate to participate in the TCA cycle ^95,96^. When cells are stimulated by inflammation, the massive consumption of GSH inhibits the TCA cycle, triggering a reprogramming of energy metabolism and initiating mitophagy to protect cells ^97,98^. As upstream regulators of GSH metabolism, spermine and spermidine directly contribute to antioxidant processes to reduce GSH consumption^99^. They may also enhance GSH synthesis by promoting glutathione synthase activity and facilitate GSH regeneration via increased glutathione reductase activity^100^. In our study, we observed that the reduction in GSH and polyamine metabolism with aging and during AD progression occurred primarily in microglia. Interestingly, instead of being activated, microglia with disrupted GSH and polyamine metabolism showed smooth branches and reduced complexity, along with a downregulation of several genes implicated in chemotaxis, suggesting that this may affect the development and function of microglia, hindering their ability to clear Aβ in the early pathological stages of AD, rather than causing their overactivation and subsequent toxic effects on neurons, thereby worsening the progression of AD.

Nevertheless, only a fraction of the metabolites in brain cells have been identified. As the annotation of metabolomic databases continues to advance, those currently unidentified metabolites could provide critical insights and potentially reveal pathways underlying cell type–specific metabolism. Additionally, only three types of brain cells were investigated in our study. Metabolic profiling of other neural cells such as oligodendrocytes, NG2 glia, and vascular cells (including endothelial cells, pericytes, and smooth muscle cells) will also be critical for understanding metabolism in the brain.

## Acknowledgments

We thank lab members in the Ge laboratory for their advice and feedback and B. Samuels for critical reading of the manuscript. We thank Drs. Qingchun Guo, Xinwei Gao, and Mingyue Jia in the imaging core of CIBR for image analysis; Drs. Wenlong Li, Ruirui Shen, and Shufang Huang in the animal core facility of CIBR for animal care; Jinmei Chen and Xuefang Zhang in the genomics center of CIBR for assistance with FACS; and Drs. Li Zhang, Xinshuang Zhang, and Hanzong Liu of the genomics center of CIBR for support for RNA sequencing and analysis. This work received support from STI2030-Major Projects 2022ZD0204700; the CAMS Innovation Fund for Medical Sciences (CIFMS, 2024-I2M-ZD-012); the Youth Beijing Scholar Program (no. 065); the Natural Science Foundation of China (32170964); funds from the CIBR to W.G.; a grant from the National Natural Science Foundation of China to X.H. (82171367); a grant from the Beijing Natural Science Foundation Program and Scientific Research Key Program of the Beijing Municipal Commission of Education, China, to X.H. (KZ201710025016); and a grant from the China Postdoctoral Science Foundation to J.Y. (2024M752878).

## Author Contributions Statement

W.G. conceived and supervised the project. W.G., J.Y., X.H., and F.L. designed the experiments. F.L., X.H., J.Y., and Z.F. provided transgenic animals and performed animal experiments. F.L. and X.H. purified various brain cells. J.Y., F.L., and X.H. extracted metabolites. X.H., W.G., and L.G.Z. performed the targeted metabolomics. J.Y., D.Y., X.Y., and F.L. performed the untargeted metabolomics and analysis. J.Y., X.H., F.L., and W.G. completed the targeted metabolomics analysis. F.L., J.Y., X.Z. and L.Z. extracted RNA and constructed a cDNA library. J.Y. completed the RNA-seq analysis. J.Y., X.C., and C.M. performed immunostaining and imaging. Z.B. assisted in online data visualization and website development. L.D. assisted in GSH measurement. W.G., B.L., Z.L., and R.J.D. provided reagents. R.J.D. advised on the data interpretation. J.Y. prepared all figures F.L. assisted in some figure preparation. J.Y., F.L., and W.G. wrote the manuscript. All authors discussed, reviewed, and edited the manuscript.

---

**Extended Data Fig. 1.**
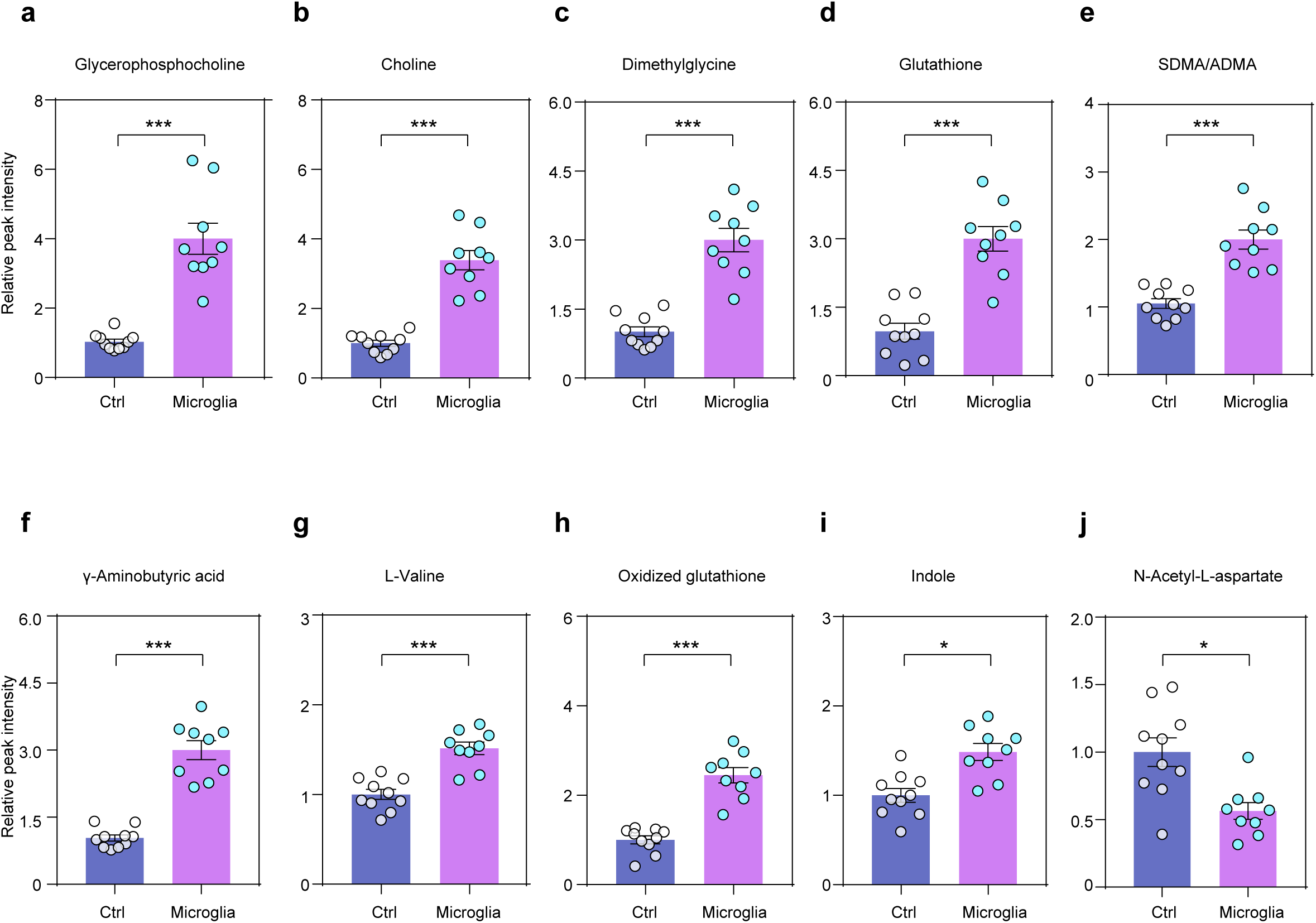
Representative metabolites in microglia. (a-j) Relative peak intensities of representative metabolites between microglia and the control samples from mouse brains, including glycerophosphocholine (a), choline (b), dimethylglycine (c), GSH (d), SDMA/ADMA (e), γ-aminobutyric acid (f), L-valine (g), oxidized glutathione (h), indole (i), *N*-acetyl-L-aspartate (j). Ctrl (n = 10 samples), Microglia (n = 9 samples). Data are presented as the mean ± SEM; two-tailed unpaired *t*-test; \**P* < 0.05, \*\*\**P* < 0.001. SDMA/ADMA, symmetric dimethylarginine/asymmetric dimethylarginine. GSH, glutathione (reduced); Ctrl, control.

**Extended Data Fig. 2.**
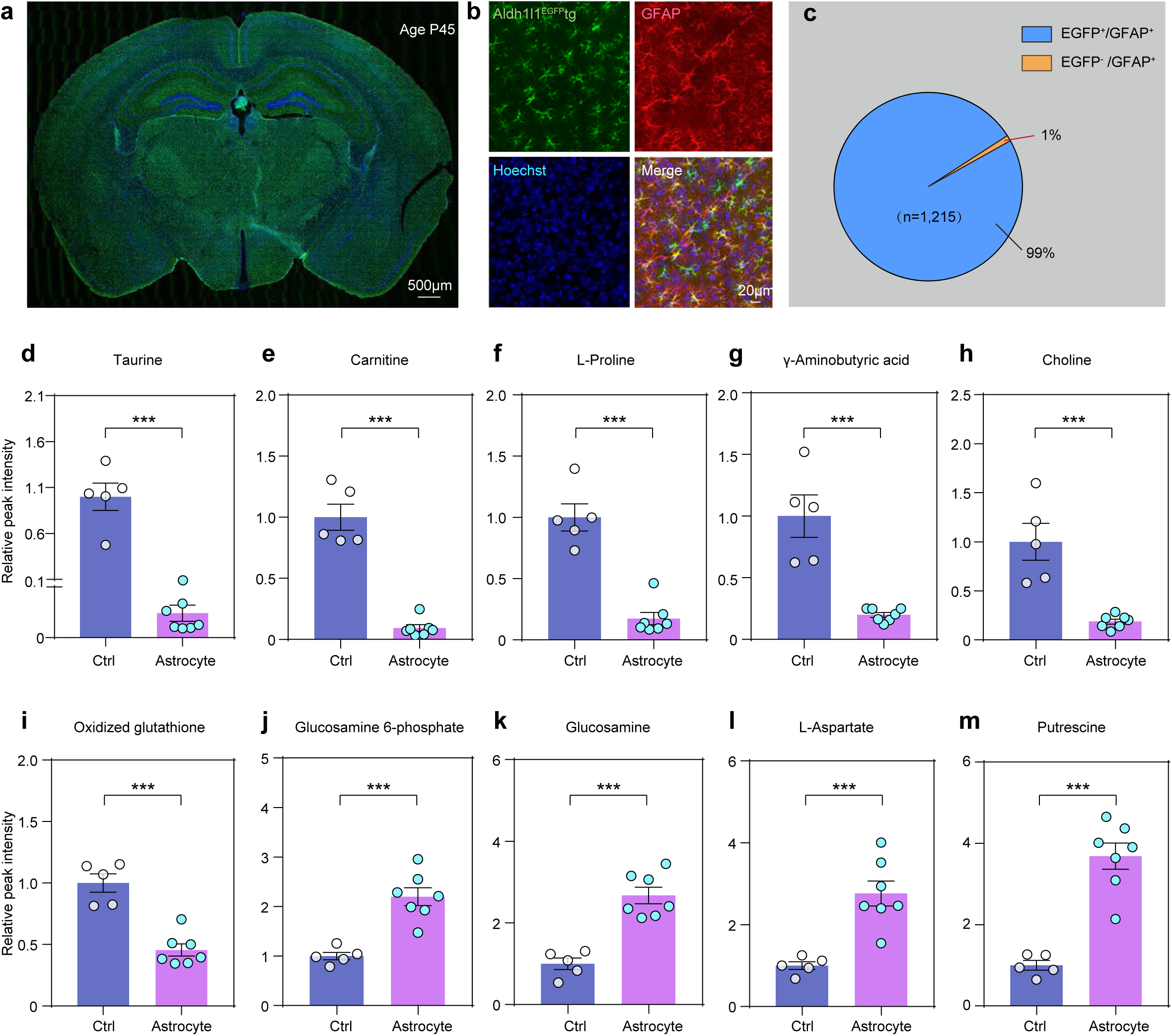
Astrocyte identification and representative metabolites in *Aldh1l1^EGFP^tg* mice. **(a)** Image of a coronal brain section from the brain of an *Aldh1l1^EGFP^tg* mouse. Mouse age, postnatal day 45. **(b)** Image of astrocytes in the hippocampus of an *Aldh1l1^EGFP^*tg mouse. Astrocytes (GFAP^+^, red) with EGFP (EGFP^+^GFAP^+^) or without EGFP (EGFP^−^GFAP^+^) expression in the hippocampal region were included for analysis in (c). Nuclei were stained with Hoechst 33342 (Hoechst, blue). **(c)** Percentage of astrocytes labeled with EGFP expression in *Aldh1l1^EGFP^tg* mice. Percentage was calculated as 100% × (EGFP^+^GFAP^+^ cells/GFAP^+^ cells). **(d–m)** Comparison of the relative peak intensities of representative metabolites in astrocytes (n = 7 samples) and the control samples (n = 5 samples) in *Aldh1l1^EGFP^tg* mice, including taurine (d), carnitine (e), L-proline (f), γ-aminobutyric acid (g), choline (h), oxidized glutathione (i), glucosamine 6-phosphate (j), glucosamine (k), L-aspartate (l), and putrescine (m). Data are shown as the mean ± SEM; two-tailed unpaired *t*-test; \*\*\**P* < 0.001.

**Extended Data Fig. 3.**
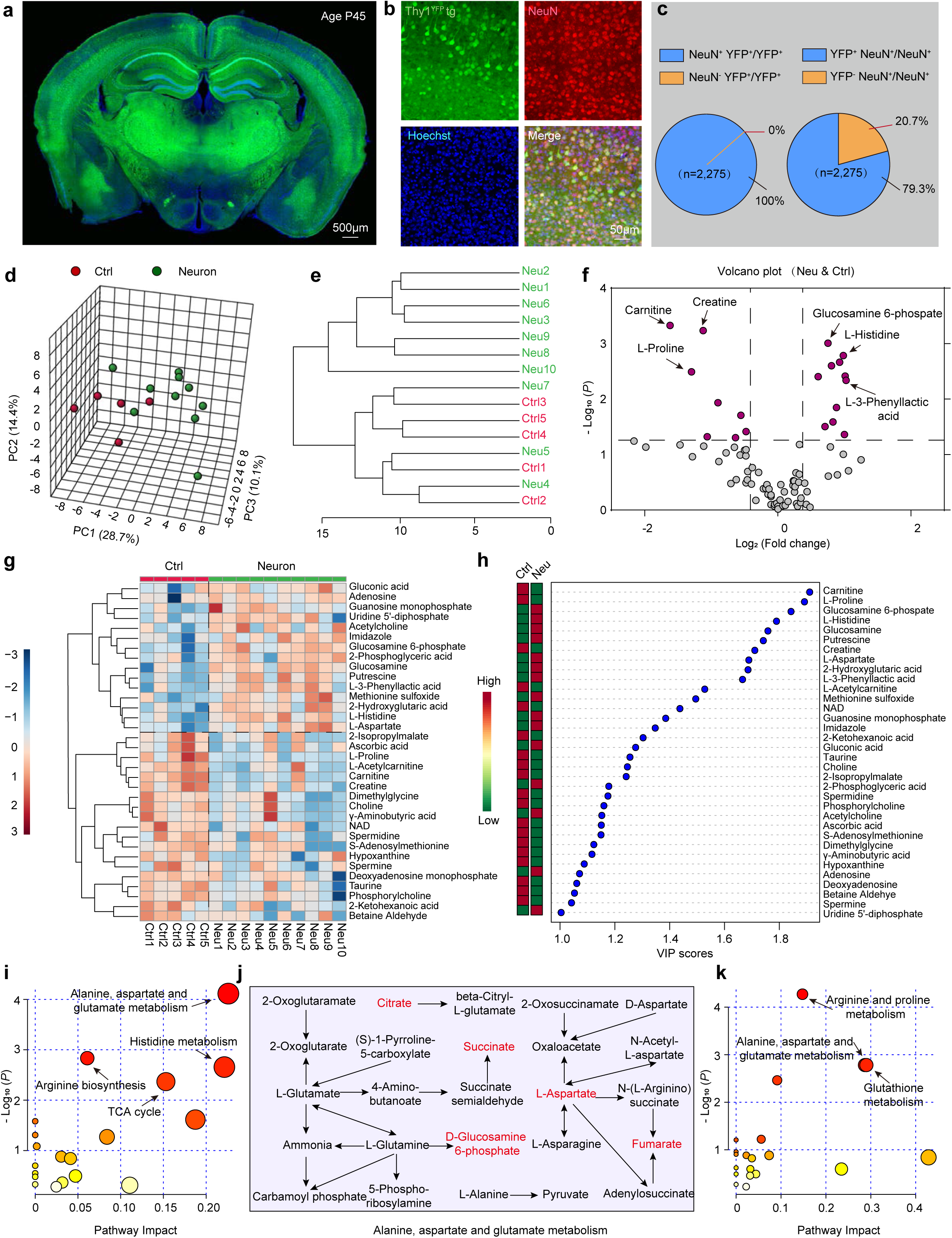
The metabolome of neurons identified by targeted metabolomics. **(a)** Image of a coronal brain section from a P45 *Thy1^YFP^tg* mouse. **(b)** Image of YFP^+^ neurons in the cerebral cortex of a *Thy1^YFP^tg* mouse. Neuron, NeuN^+^(red), nuclei, Hoechst 33342 (Hoechst, blue). **(c)** Percentage of neurons with YFP expression in *Thy1^YFP^tg* mice was calculated as 100% × (NeuN^+^YFP^+^ cells/YFP^+^ cells) and 100% × (NeuN^−^YFP^+^ cells/YFP^+^ cells). Percentage of neurons with YFP expression in *Thy1^YFP^tg* mice was calculated as 100% × (YFP^+^NeuN^+^ cells/NeuN^+^ cells) and 100% × (YFP^−^NeuN^+^ cells/NeuN^+^ cells). **(d)** PCA of the metabolomes of neurons (n = 10 samples) and the control group (i.e., YFP^−^ cells, n = 5 samples). In total, 82 metabolites were identified with targeted metabolomics from each sample and used for PCA.. **(e)** Dendrogram showing the clustering of all samples. **(f)** Volcano plot of differential metabolites between the two groups. Differential (purple) and non-differential (gray) metabolites were defined by the criteria of fold change > 1.2 and *P* < 0.05 (two-tailed unpaired *t*-test). A data point with a positive value indicates a high level of a metabolite in neurons. **(g)** Heatmap representation of 34 of 82 metabolites in the two groups, as identified by VIP scores from a PLS-DA. Red indicates a greater abundance of the metabolite. **(h)** Top 34 metabolites (VIP scores > 1) for differentiating between neurons and the control group. The colored boxes on the left indicate the relative concentrations of the corresponding metabolite averaged across each group. Ctrl, control; Neu, neurons. **(i–k)** A summary of the pathway analysis using metabolites found at high levels (fold change > 1.2) in neurons as compared with the control group (i) or with high levels in the control group as compared with neurons (k). The size of the circle represents pathway impact, and the color represents the *P*-value. A portion of the Alanine, aspartate and glutamate metabolism pathway in neurons and the control group. Metabolites in red were present at high levels in neurons as compared with the control group. TCA, tricarboxylic acid; GSH, glutathione (reduced); Ctrl, control; Neu, neurons.

**Extended Data Fig. 4.**
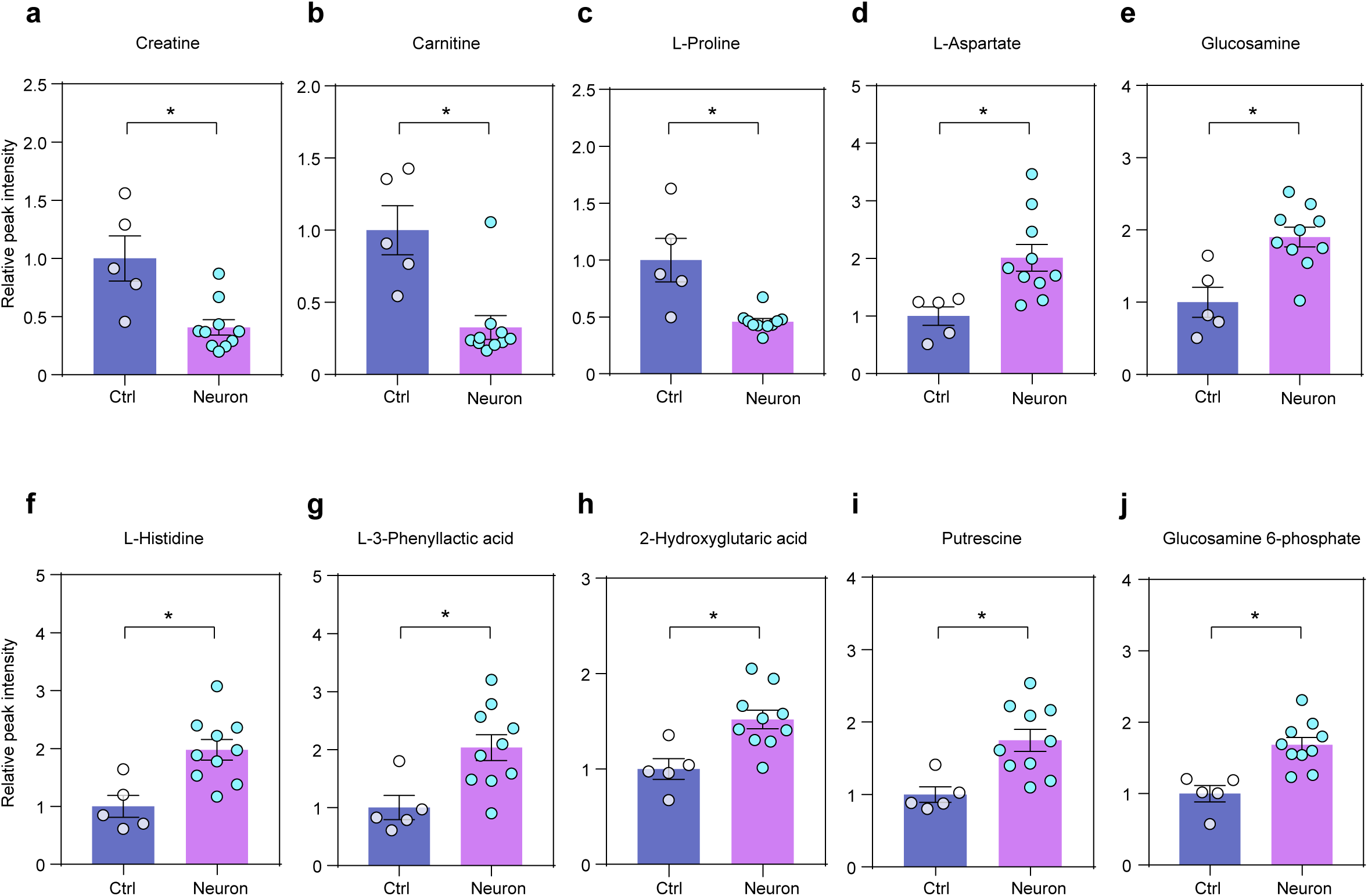
Representative metabolites in neurons. (a-j) Comparison of the relative intensities of representative metabolites in neurons and the control samples in mice, including creatine (a), carnitine (b), L-proline (c), L-aspartate (d), glucosamine (e), L-histidine (f), L-3-phenylacetic acid (g), 2-hydroxyglutaric acid (h), putrescine (i), and glucosamine 6-phosphate (j). Data are represented as the mean ± SEM; two-tailed unpaired *t*-test; \**P* < 0.05; Ctrl, control. Ctrl (n = 5 samples), neuron (n = 10 samples).

**Extended Data Fig. 5.**
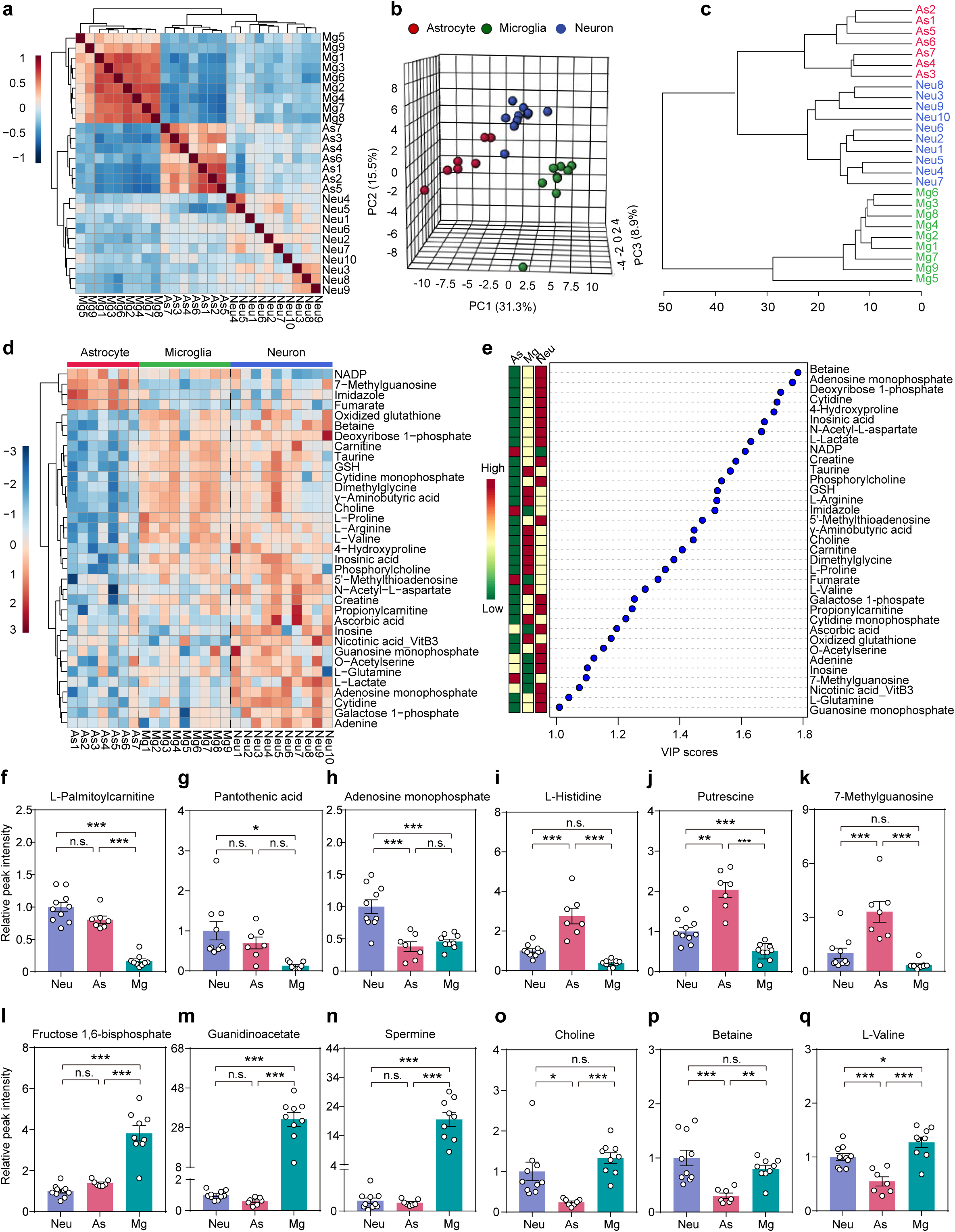
Comparisons of the metabolomes among neurons, astrocytes, and microglia from targeted metabolomics. **(a)** A heatmap showing the correlation coefficients of the metabolomes of neurons, astrocytes, and microglia. **(b)** Principal component analysis (PCA) of the metabolomes of neurons, astrocytes, and microglia. In total, 95 metabolites were identified from all cell samples and used for PCA. Three PCs explained 55.7% (31.3%, 15.5%, and 8.9%) of variance among the three groups. PC scores are indicated as percentages; circles indicate individual samples from neurons, astrocytes, and microglia. **(c)** Dendrogram showing the clustering of all samples. **(d)** Heatmap representation of 35 of 95 metabolites in the three groups, as identified by VIP scores from a PLS-DA. Each column represents a sample from a different mouse. The red color indicates a greater abundance of a metabolite. PLS-DA, partial least-squares discriminant analysis; VIP, variable importance in projection. **(e)** Top 35 metabolites (based on VIP scores) for differentiating among the three cell types. The colored boxes on the left indicate the relative concentrations of corresponding metabolites averaged across each group. **(f–q)** Relative peak intensity of representative metabolites among the neurons, astrocytes, and microglia, including L-palmitoylcarnitine (f), pantothenic acid (g), adenosine monophosphate (h), L-histidine (i), putrescine (j), 7-methylguanosine (k), fructose 1,6-bisphosphate (l), guanidinoacetate (m), spermine (n), choline (o), betaine (p), and L-valine (q). Data are represented as the mean ± SEM; one-way ANOVA; \**P* < 0.05, \*\**P <* 0.01, \*\*\**P <* 0.001; n.s., no significant difference. ANOVA, analysis of variance; Neu, neurons; As, astrocytes; Mg, microglia. In (a–d, f–q): As, n = 7 samples; Mg, n = 9 samples; Neu, n = 10 samples.

**Extended Data Fig. 6.**
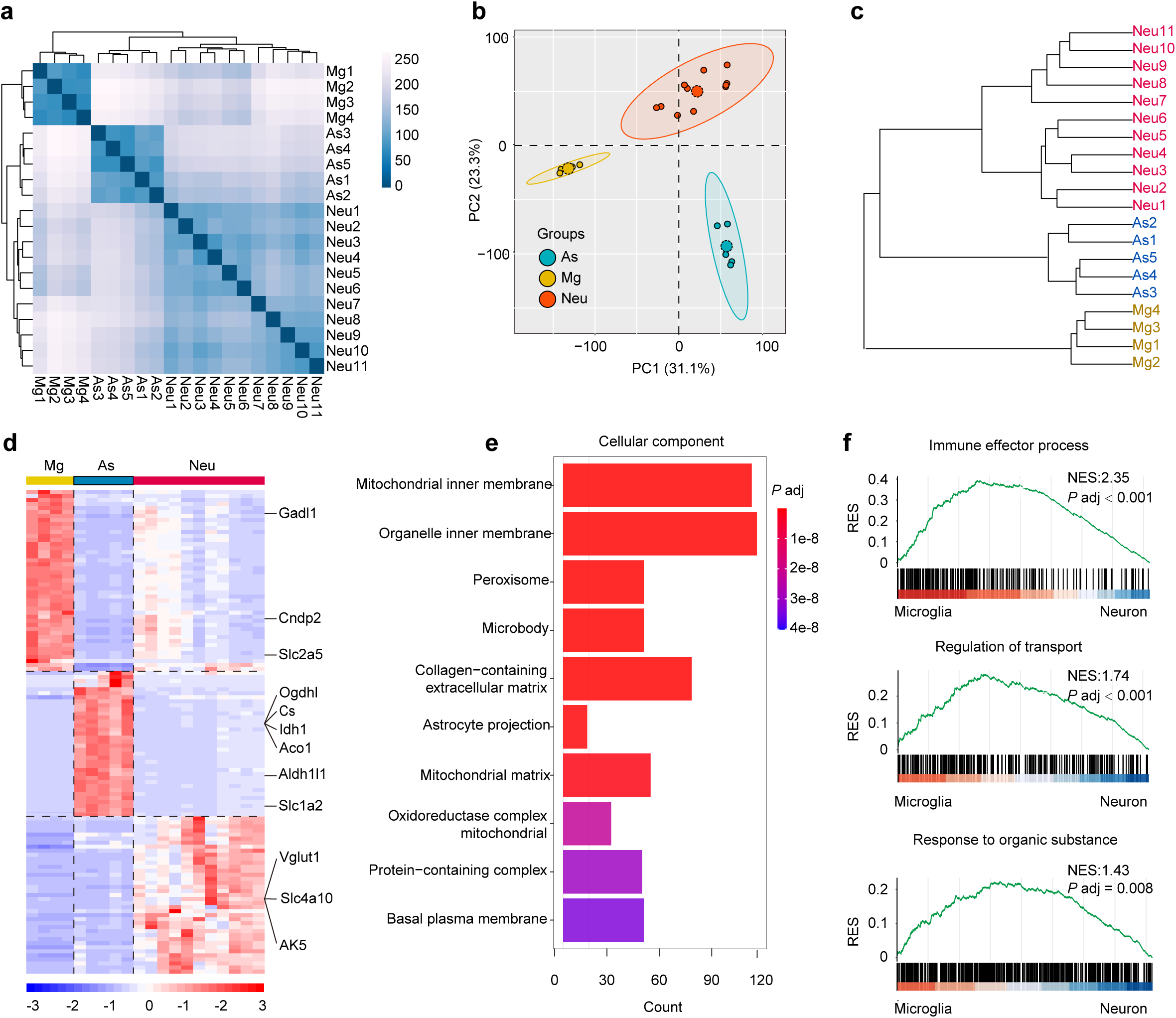
Comparison of the transcriptome among neurons, astrocytes, and microglia. **(a)** Spearman’s rank correlation of all transcriptomes obtained from neurons (n = 11 samples), astrocytes (n = 5 samples), and microglia (n = 4 samples). Each sample was obtained from a different mouse. In addition, gene expression profiles of samples from the same cell type showed a high degree of correlation. **(b)** Principal component analysis (PCA) of the transcriptomes of the samples from neurons, astrocytes, and microglia. In total, 16,649 genes were used for the PCA. Two PCs explained 54.4% (31.1% and 23.3%) of the variance among the three groups. PC scores are indicated as percentages; circles indicate individual samples from neurons, astrocytes, and microglia. **(c)** Dendrogram analysis showing the clustering of all samples. In (a–c): Ast, n = 5 samples; Mg, n = 4 samples; Neu, n = 11 samples. **(d)** Heatmap representation of the top genes from the three groups, as identified by fold change, *P*-values, and transcript abundance. Each column represents a sample: Ast, n = 5; Mg, n = 4; Neu, n = 11. The red color indicates the greater transcript abundance of a metabolic enzyme among all samples. **(e)** Gene Ontology (GO) analysis between astrocytes and neurons. The bar plot depicts the top 10 enriched GO terms within categories corresponding to the term ‘cellular component’, based on a hypergeometric test. The length of the bar represents the gene counts; the color of the bar represents the significance of enrichment, with red representing lower *P*-values and purple representing higher *P*-values **(f)** Representative gene set enrichment analysis (GSEA) between microglia and neurons. Three functional gene sets are shown with enrichment significance assessed using permutation-based testing. Black bars in the GSEA indicate the hits in gene sets represented among all genes pre-ranked by ranking metrics. RES, running enrichment score; NES, normalized enrichment score.

**Extended Data Fig. 7.**
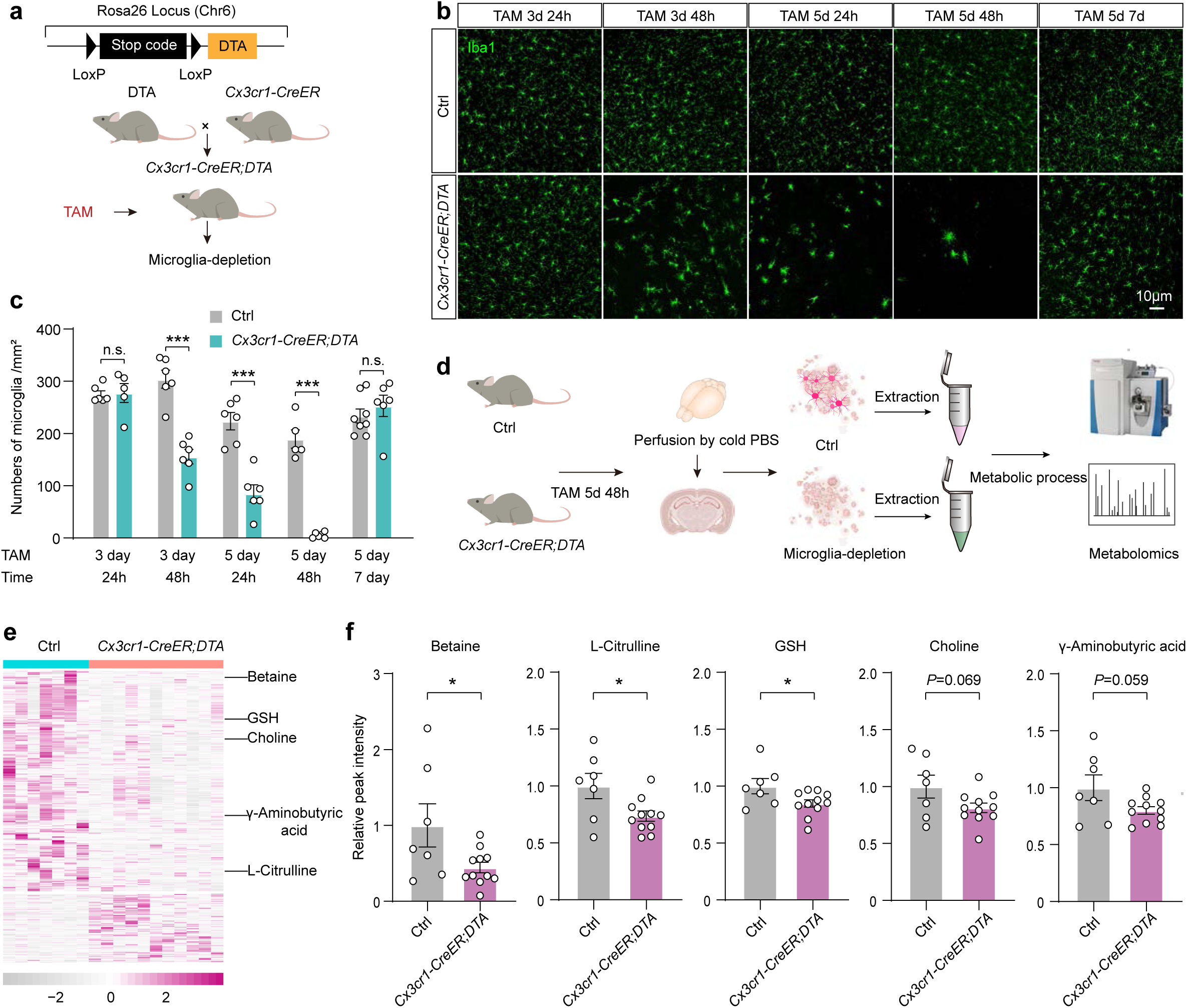
Genetic ablation of microglia confirms microglia-dependent metabolic alterations in the brain. (a) Schematic of the genetic microglial depletion strategy using *Cx3cr1-CreER;DTA* mice. (b) Immunohistochemical validation of microglial depletion at different time points after tamoxifen injection. (c) Quantification of microglial density across different time points (24 h, 48 h, and 7 d) following a 3- or 5-d tamoxifen treatment. 3 d, 24 h: Ctrl, n = 5 samples; *Cx3cr1-CreER;DTA*, n = 5 samples. 3 d, 48 h: Ctrl, n = 6 samples; *Cx3cr1-CreER*, n = 6 samples. 5 d, 24 h: Ctrl, n = 6 samples; *Cx3cr1-CreER;DTA*, n = 6 samples. 5 d, 48 h: Ctrl, n = 5 samples; *Cx3cr1-CreER;DTA*, n = 5 samples. 5 d, 7 d: Ctrl, n = 8 samples; *Cx3cr1-CreER;DTA*, n = 6 samples. (d) Experimental workflow for brain metabolomic profiling 48 h after microglial ablation. (e) Comparison of key metabolites (choline, betaine, GSH, L-citrulline, and γ-aminobutyric acid) between *Cx3cr1-CreER;DTA* mice and controls. The purple color indicates the greater abundance of a metabolite among all samples. Ctrl, n = 7 samples; *Cx3cr1-CreER;DTA*, n = 11 samples. (f) Relative peak intensity of representative metabolites between the control (n = 7 samples) and *Cx3cr1-CreER;DTA* (n = 11 samples) groups, including betaine, L-citrulline, GSH, choline, and γ-aminobutyric acid. Data are presented as the mean ± SEM; two-tailed unpaired *t*-test; \**P* < 0.05; \*\*\**P* < 0.001.

**Extended Data Fig. 8.**
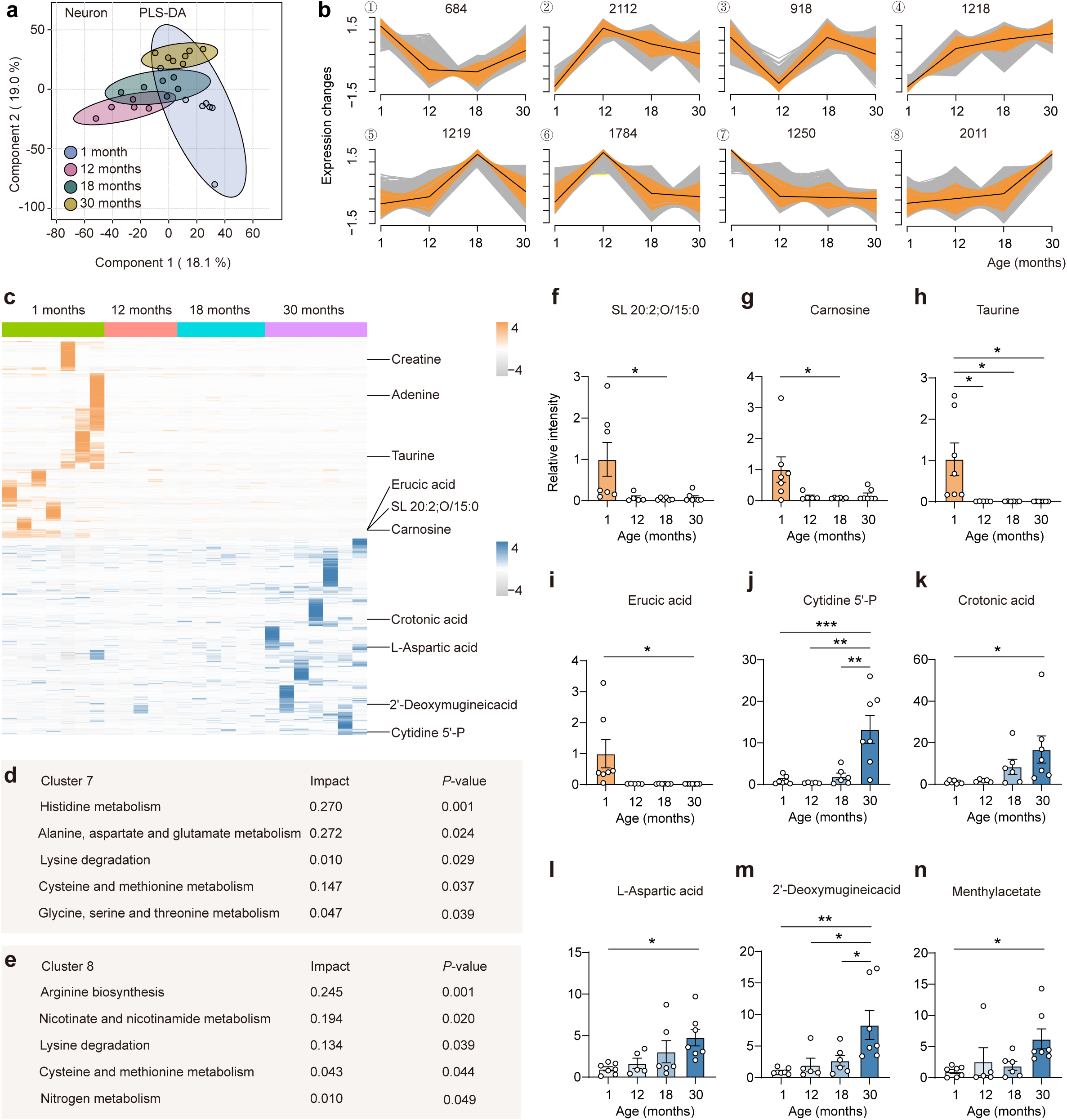
Characterizing the metabolome of neurons from aging brains. (a) Partial least-squares discriminant analysis (PLS-DA) of metabolomes of aging neurons. Two components explained 37.1% (18.1% and 19.0%) of the variance among the four groups. Component scores are indicated as percentages; circles indicate individual samples of neurons from different age groups (1, 12, 18, and 30 months). (b) Clustering of time course expression patterns in neurons from mouse brains using the fuzzy c-means algorithm. Warm (orange) and cold (gray) colors indicate low and high deviation from the consensus profile, respectively. (c) Heatmap representation of top metabolites of the four groups from clusters 7 and 8, as identified by fold change, *P*-values (two-tailed unpaired *t*-test), and abundance. Each column represents a sample. Orange or blue color indicates greater abundance of a metabolite, whereas gray indicates lower abundance. (d) Top five metabolic pathways based on KEGG. The analyzed metabolites were from cluster 7 in (c). KEGG, Kyoto Encyclopedia of Genes and Genomes. Pathway enrichment was performed using a hypergeometric test and pathway topology analysis based on relative-betweenness centrality. (e) Top five metabolic pathways based on KEGG. The analyzed metabolites were from cluster 8 in (c). KEGG, Kyoto Encyclopedia of Genes and Genomes. Pathway enrichment was performed using a hypergeometric test and pathway topology analysis based on relative-betweenness centrality. (**f–n**) Relative peak intensity of SL 20:2;O/15:0 (f, cluster 7), carnosine (g, cluster 7), taurine (h, cluster 7), erucic acid (i, cluster 7), cytidine 5’-P (j, cluster 8), crotonic acid (k, cluster 8), L-aspartic acid (l, cluster 8), 2’-deoxymugineicacid (m, cluster 8) and methylacetate (n, cluster 8) in neurons from mouse brains of different ages. Data are represented as the mean ± SEM; one-way ANOVA; \**P* < 0.05, \*\**P* < 0.01, \*\*\**P* < 0.001. ANOVA, analysis of variance. In (a, c, f–n): 1 month, n = 7 samples; 12 months, n =5 samples; 18 months, n =6 samples; 30 months, n = 7 samples.

**Extended Data Fig. 9.**
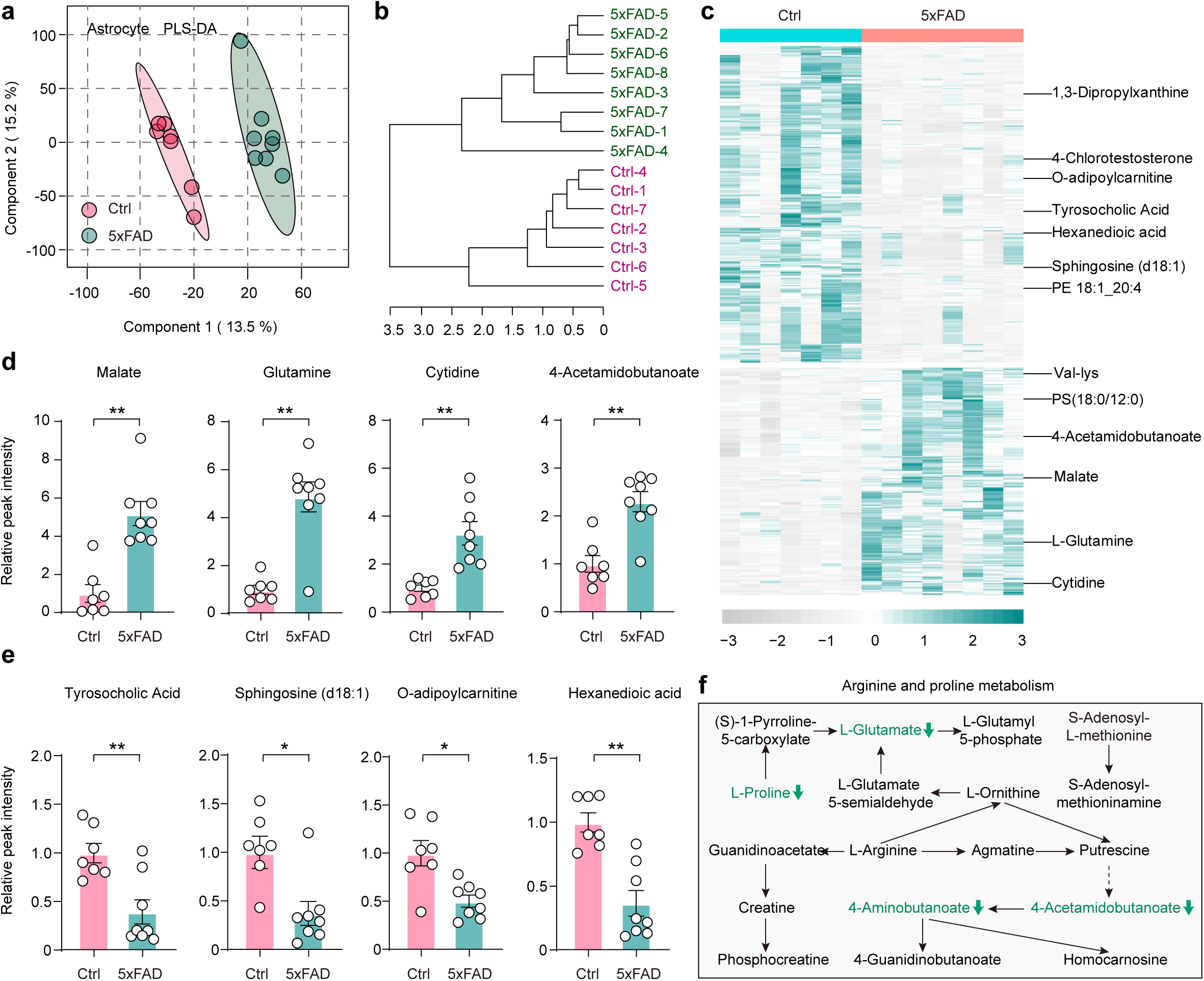
Characterizing the astrocyte metabolome in the brain of AD mouse models. **(a)** Partial least-squares discriminant analysis (PLS-DA) of astrocyte metabolomes of 5xFAD and wild type mouse brains. Two components explained 28.7% (13.5% and 15.2%) of the variance between the two groups. Component scores are indicated as percentages; circles indicate individual samples of astrocytes from different groups. **(b)** Dendrogram showing the clustering of all astrocyte samples. **(c)** Heatmap representation of top metabolites (top 727, *P* < 0.05) in the two groups. Each column represents a sample from a different mouse. Green color indicates a greater abundance of a metabolite. **(d)** Relative peak intensities of representative metabolites increased in astrocytes of the 5xFAD group compared to the control group, including malate, glutamine, cytidine and 4-acetmidobutanoate. **(e)** Relative peak intensities of representative metabolites decreased in astrocytes of the 5xFAD group compared to control group, including tyrosocholic acid, sphingosine (d18:1), O-adipoylcarnitine and hexanedioic acid. **(f)** A portion of the arginine and proline metabolism pathway of astrocytes differed between the 5xFAD group and control group. Metabolites in green were present at lower levels in the astrocytes of the 5xFAD group compared with the control group. In (a–e): Ctrl, n = 7 samples; 5xFAD, n = 8 samples. Data are presented as the mean ± SEM; two-tailed unpaired *t*-test; \**P* < 0.05, \*\**P* < 0.01.

**Extended Data Fig. 10.**
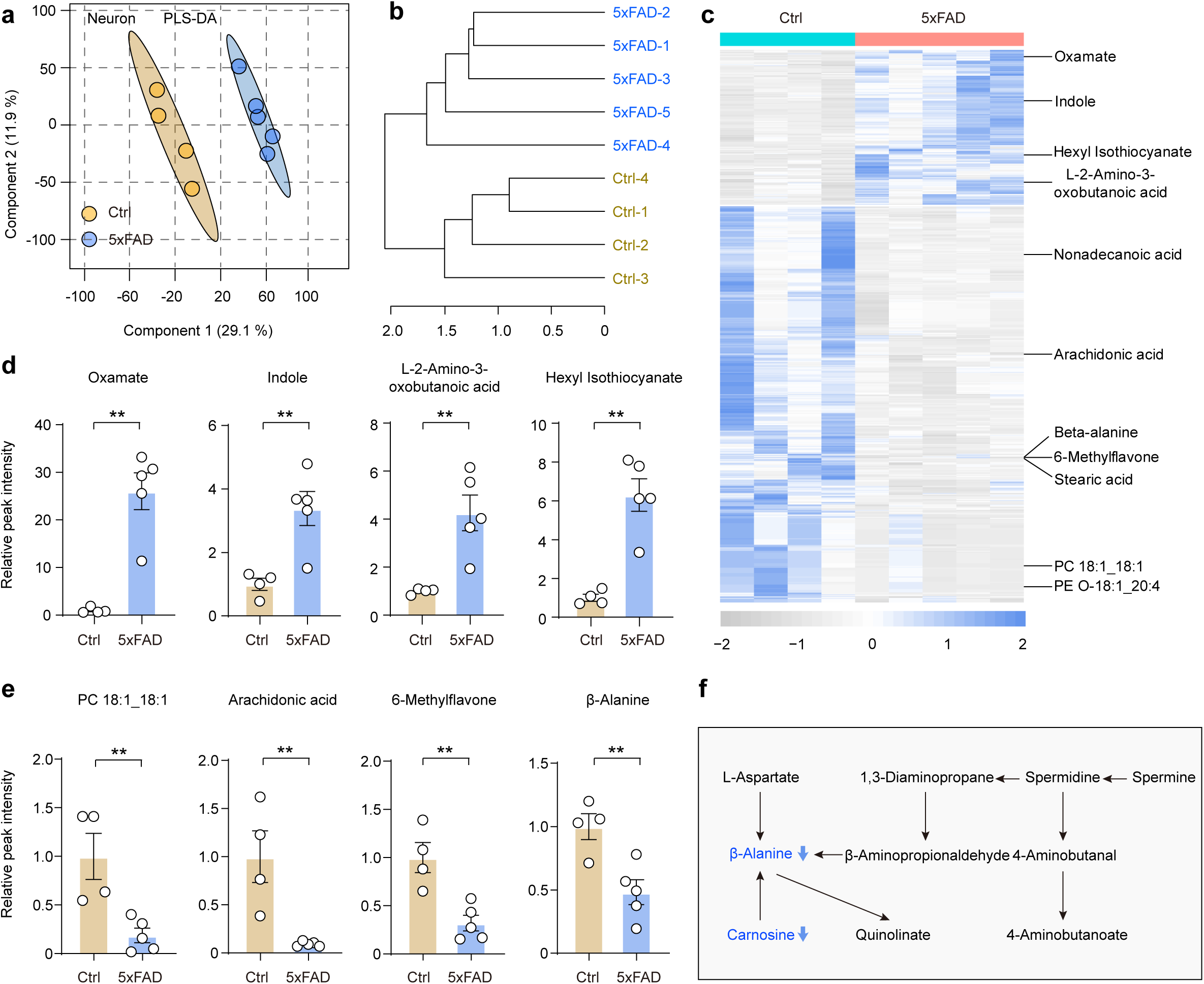
Characterizing the neuronal metabolome in the brain of AD mouse models. **(a)** Partial least-squares discriminant analysis (PLS-DA) of neuronal metabolomes of mouse brains. Two components explained 41% (29.1% and 11.9%) of the variance between the two groups. Component scores are indicated as percentages; circles indicate individual samples of neurons from different groups. **(b)** Dendrogram showing the clustering of all neuronal samples. **(c)** Heatmap representation of top metabolites (top 619, *P* < 0.05) in the two groups. Each column represents a sample from a different mouse. Blue color indicates a greater abundance of a metabolite. **(d)** Relative peak intensities of representative metabolites increased in neurons of the 5xFAD group compared to the control group, including oxamate, indole, L-2-amino-3-oxobutanoic acid and hexyl isothiocyanate. **(e)** Relative peak intensities of representative metabolites decreased in neurons of the 5xFAD group compared to the control group, including PC 18:1_18:1, arachidonic acid, 6-methyiflavone and β-alanine. **(f)** A portion of the β-alanine metabolism pathway of neurons differed between the 5xFAD group and control group. Metabolites in blue were present at lower levels in the neurons of the 5xFAD group compared with the control group. In (a–e): Ctrl, n = 4 samples; 5xFAD, n = 5 samples. Data are presented as the mean ± SEM; two-tailed unpaired *t*-test; \*\**P* < 0.01.

**Supplementary Fig. 1.**
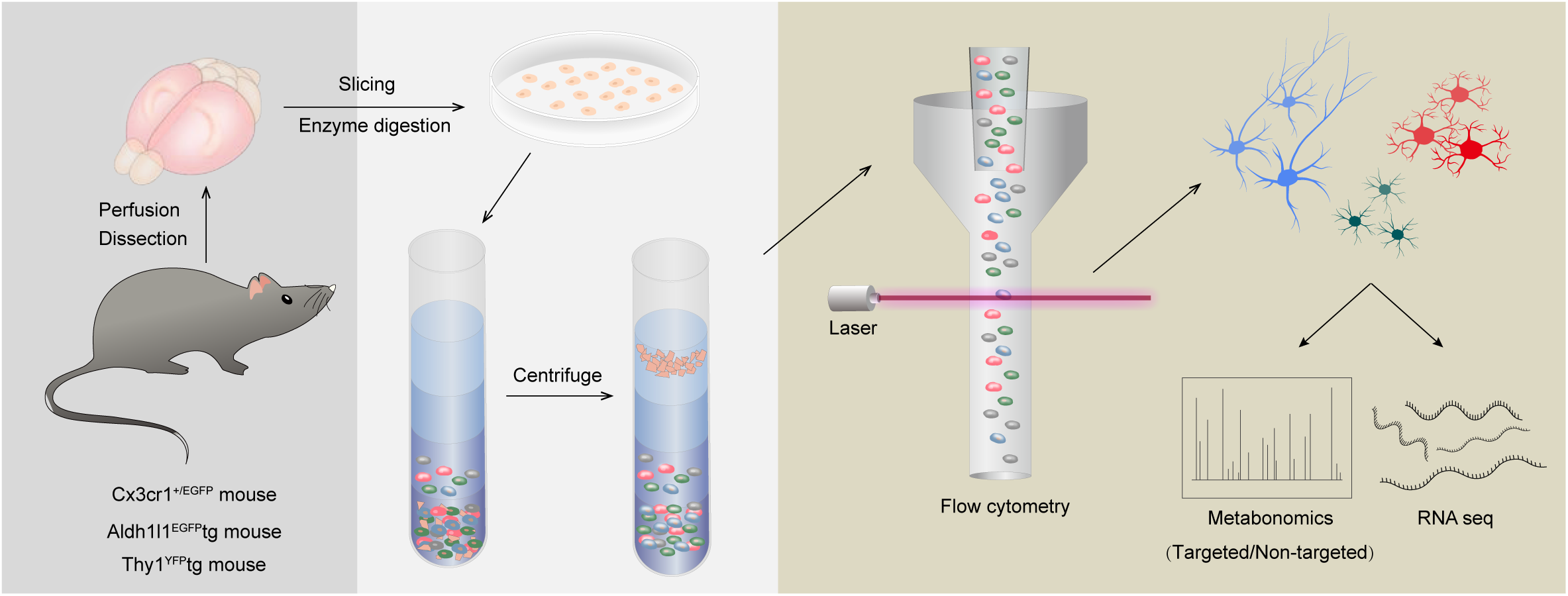
Flowchart of targeted and non-targeted analyses. Flowchart outlining the process of purifying neurons, astrocytes, and microglia for metabolomics and RNA sequencing (RNA-seq).

**Supplementary Fig. 2.**
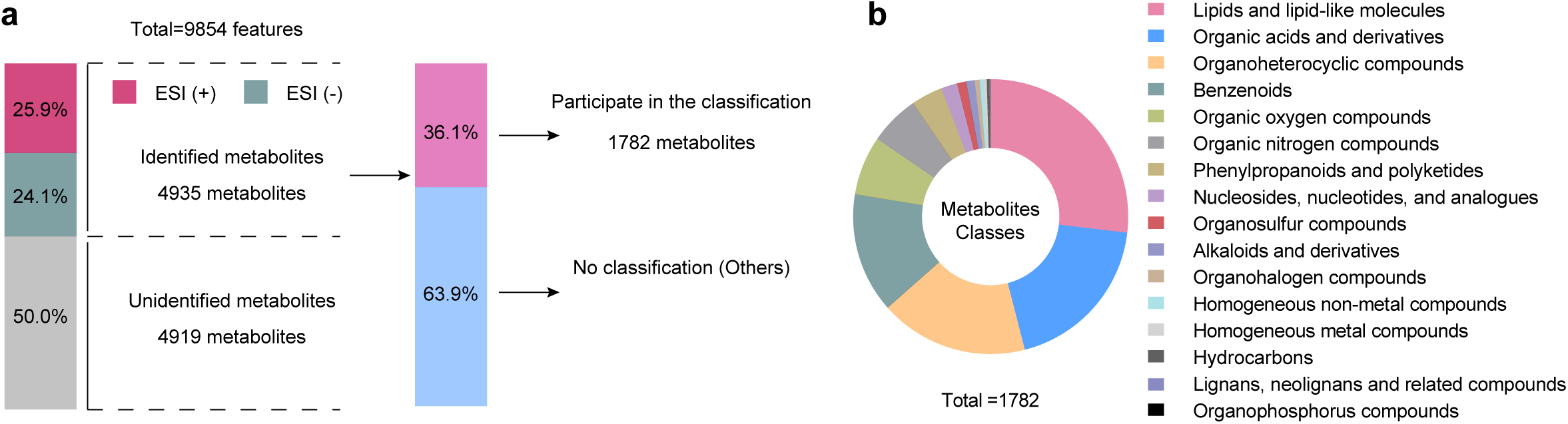
Comprehensive overview of metabolites in neurons, astrocytes, and microglia from untargeted metabolomics. **(a)** Total number of features detected in neurons, astrocytes, and microglia by untargeted metabolomics. ESI (+), electrospray ionization in positive mode; ESI (–), electrospray ionization in negative mode. Metabolomic analysis detected a total of 9,854 metabolite features, of which 4,935 (50.0%) were successfully annotated from the mzCloud libraries. The identified metabolites were classified using the Human Metabolome Database (HMDB), resulting in the successful categorization of 1,782 metabolites (36.1%). **(b)** The categorization of the identified metabolites. Metabolites were grouped into 16 major classes(Lipids and lipid-like molecules; Organic acids and derivatives; Organoheterocyclic compounds; Benzenoids; Organic oxygen compounds; Organic nitrogen compounds; Phenylpropanoids and polyketides; Nucleosides, nucleotides, and analogues; Organosulfur compounds; Alkaloids and derivatives; Organohalogen compounds; Homogeneous non-metal compounds; Homogeneous metal compounds; Hydrocarbons; Lignans, neolignans and related compounds; Organophosphorus compounds, with the relative abundance of each category quantified). Among them, the categories Lipids and lipid-like molecules, Organic acids and derivatives, Organoheterocyclic compounds, Benzenoids and Organic oxygen compounds represented the most predominant groups, ranking among the top five. The remaining 3,153 metabolite features (63.9%) lacked corresponding classifications and were therefore designated as “Others”.

**Supplementary Fig. 3.**
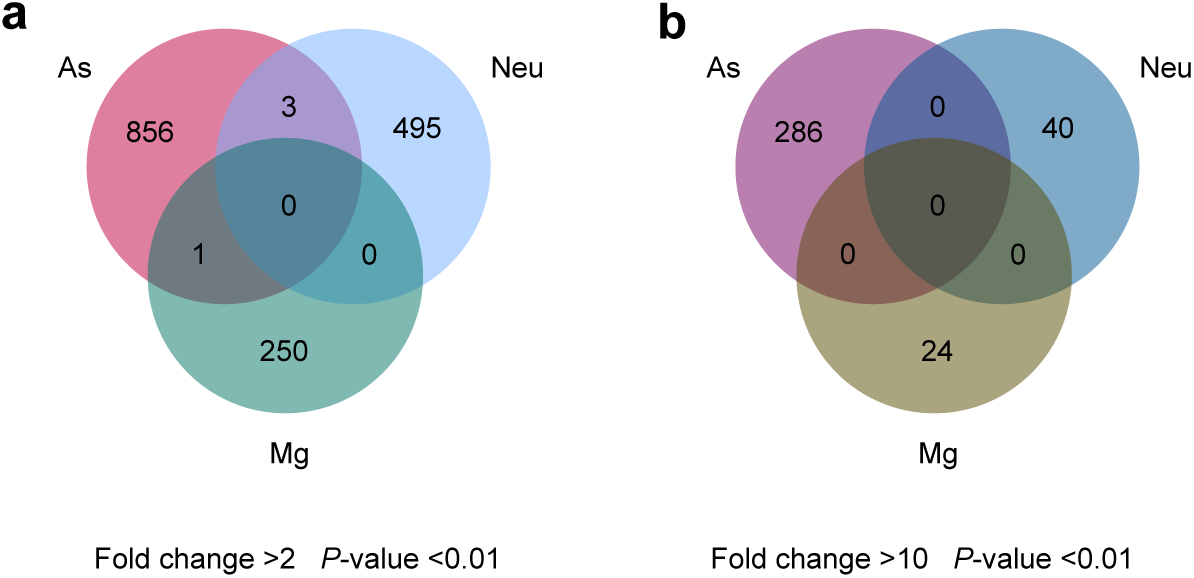
Identification of cell-type-enriched features among microglia, astrocytes, and neurons. **(a)** The numbers of features enriched in microglia, astrocytes, and neurons under the stringent criteria (fold change > 2; *P* < 0.01, two-tailed unpaired *t*-test) in a Venn diagram. Each circle represents one cell type. **(b)** The numbers of features enriched in microglia, astrocytes, and neurons under the stringent criteria (fold change > 10; *P* < 0.01, two-tailed unpaired *t*-test) in a Venn diagram. Each circle represents one cell type.

**Supplementary Fig. 4.**
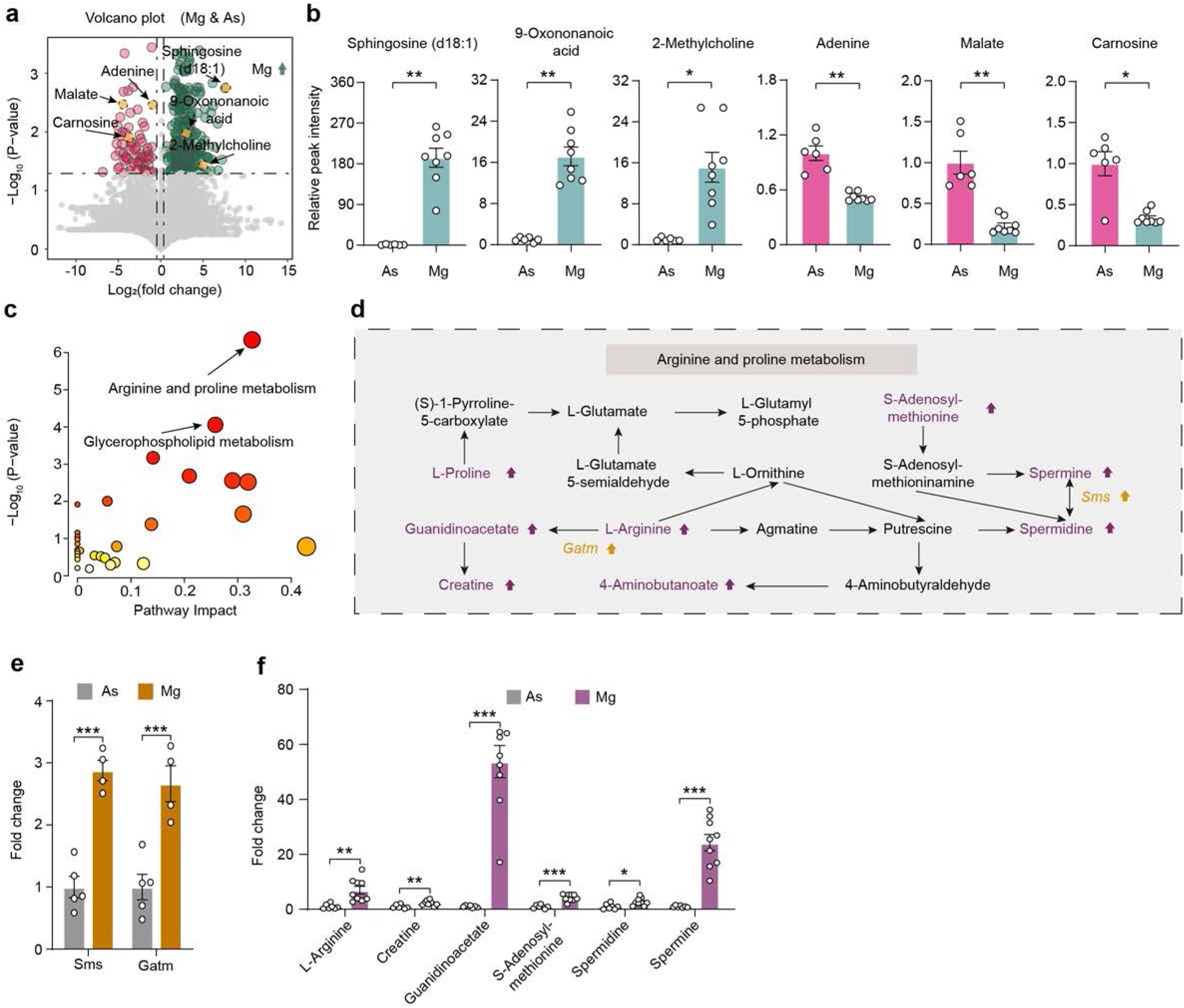
Comparison of metabolome between microglia and astrocytes. **(a)** Volcano plot of differential metabolites between microglia and astrocytes. Differential (red and green) and non-differential (gray) metabolites were defined by the criteria of fold change >1.5 and *P* adj < 0.05. A data point with a negative fold change indicates a high level of a metabolite (fold change < –1.5 and *P* adj < 0.05) in astrocytes, and a data point with a positive value indicates a high level of a metabolite in microglia. **(b)** Relative peak intensities of representative identified metabolites between astrocytes (n = 6 samples) and microglia (n = 8 samples), including sphingosine (d18:1) (b), 9-oxononanoic acid (c), 2-methylcholine (d), adenine (e), adenine (f), and carnosine (g). **(c)** A summary of the pathway analysis using metabolites found at high levels (fold change > 1.2) in microglia as compared with the astrocytes. The size of the circle represents pathway impact, and the color represents the *P*-value. Pathway enrichment was performed using a hypergeometric test and pathway topology analysis based on relative-betweenness centrality. **(d)** A portion of the Arginine and proline metabolism pathway in microglia and astrocytes. Genes in yellow and metabolites in purple were expressed at a high level in microglia when compared with astrocytes. **(e)** Comparisons of genes expression in the Arginine and proline metabolism pathway in microglia (n = 4 samples) and astrocytes (n = 5 samples). **(f)** Comparison of the abundance of metabolites in the Arginine and proline metabolism pathway in microglia (n = 9 samples) and astrocytes (n = 7 samples). Data are represented as the mean ± SEM; two-tailed unpaired *t*-test; \**P* < 0.05, \*\**P* < 0.01, \*\*\**P* < 0.001.

**Supplementary Fig. 5.**
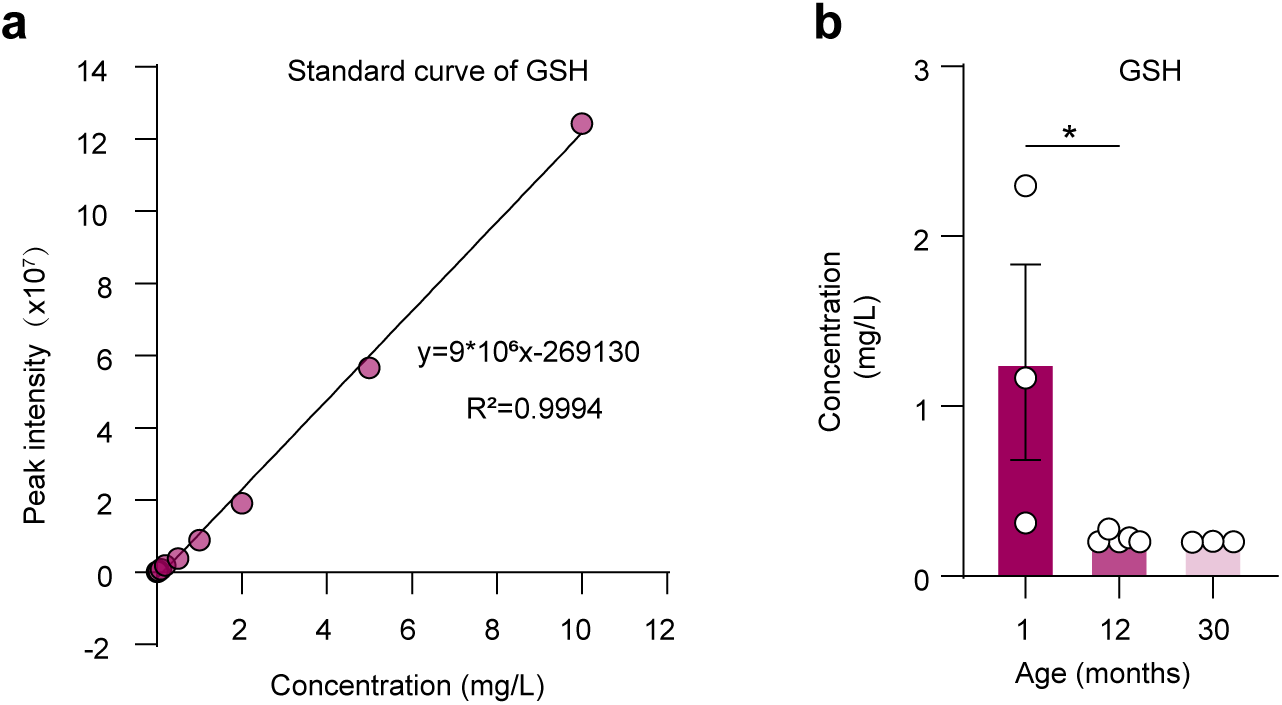
Targeted metabolomic validation of age- and AD-associated decline in microglial GSH metabolism. (a) Standard curve for absolute quantification of GSH in targeted metabolomics across different ages of mice. (b) Absolute GSH levels in microglia isolated from mice at 1 (n = 3 samples), 12 (n = 5 samples), and 30 (n = 3 samples) months of age (∼60,000 cells per sample). Data are represented as the mean ± SEM; one-way ANOVA; \**P* < 0.05.

**Supplementary Fig. 6.**
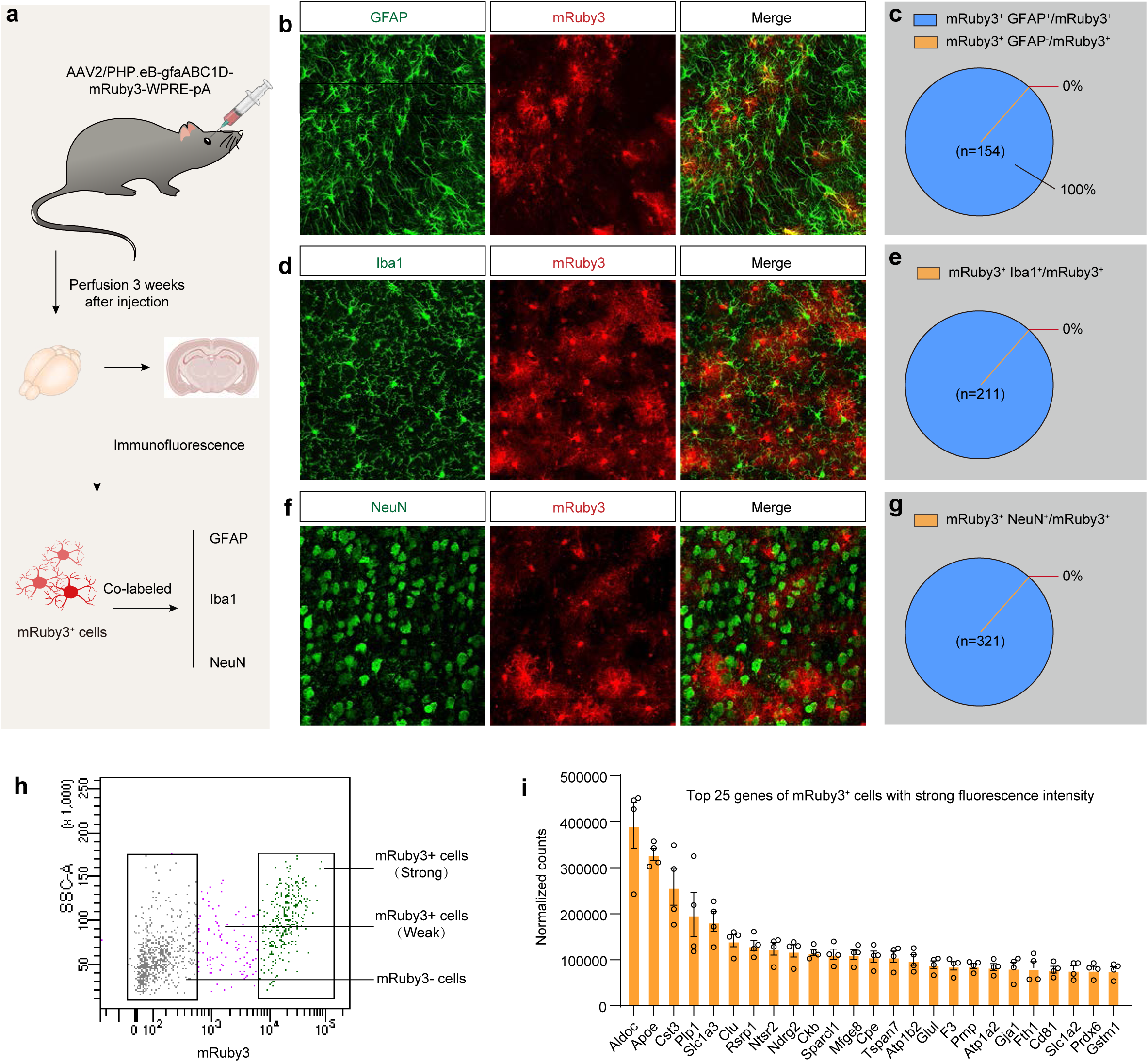
Verification of purity of *AAV2/PHP.eB-gfaABC1D-mRuby3-WPRE-pA*-labeled astrocytes. (a) Flowchart outlining the process of adeno-associated virus injection and immunofluorescence. (b) Image of astrocytes in the hippocampus labeled with *AAV2/PHP.eB-gfaABC1D-mRuby3-WPRE-pA*. mRuby3^+^ (red) cells were stained with anti-GFAP (green). (c) Percentage of mRuby3^+^ cells with strong fluorescence intensity in astrocytes from the hippocampus. The percentage was calculated as 100% × (mRuby3^+^GFAP^+^ cells/mRuby3^+^ cells). Percentage of mRuby3^+^ cells with strong fluorescence intensity in the other brain cells from the hippocampus. Percentage was calculated as 100% × (mRuby3^+^GFAP^−^ cells/mRuby3^+^ cells). (d) Image of microglia in cerebral cortex labeled with *AAV2/PHP.eB-gfaABC1D-mRuby3-WPRE-pA*. mRuby3^+^ (red) cells were stained with anti-Iba1 (green). (e) Percentage of mRuby3^+^ cells with strong fluorescence intensity in microglia from cerebral cortex. The percentage was calculated as 100% × (mRuby3^+^Iba1^+^ cells/mRuby3^+^ cells). (f) Image of neurons in cerebral cortex labeled with *AAV2/PHP.eB-gfaABC1D-mRuby3-WPRE-pA*. mRuby3^+^ (red) cells were stained with anti-NeuN (green). (g) Percentage of mRuby3^+^ cells with strong fluorescence intensity in neurons from cerebral cortex. Percentage was calculated as 100% × (mRuby3^+^NeuN^+^ cells/mRuby3^+^ cells). (h) The clustering of mRuby3^+^ cells during fluorescence-activated cell sorting. mRuby3^−^ cells (gray); mRuby3^+^ cells with strong fluorescence intensity (green); mRuby3^+^ cells with weak fluorescence intensity (purple). (i) Normalized counts of top 25 genes in mRuby3^+^ cells with strong fluorescence intensity, including *Aldoc*, *Apoe*, *Cst3*, *Plp1*, *Slc1a3*, *Clu*, *Rsrp1*, *Ntsr2*, *Ndrg2*, *Ckb*, *Sparcl1*, *Mfge8*, *Cpe*, *Tspan7*, *Atp1b2*, *Glul*, *F3*, *Prnp*, *Atp1a2*, *Gja1*, *Fth1*, *Cd81*, *Slc1a2*, *Prdx6*, and *Gstm1*.

**Supplementary Fig. 7.**
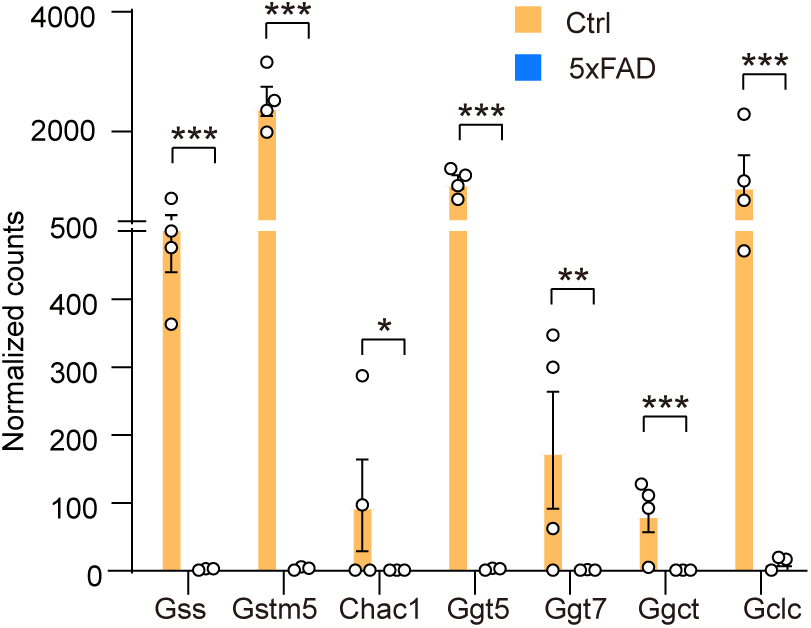
Comparisons of gene expression in microglia between the 5xFAD group and control group. Comparisons of gene expression in the glutathione metabolic pathway of microglia between the 5xFAD group (n = 3 samples) and control group (n = 4 samples). Data are presented as the mean ± SEM; two-tailed unpaired *t*-test; \**P* < 0.05, \*\**P* < 0.01, \*\*\**P* < 0.001.

**Supplementary Fig. 8.**
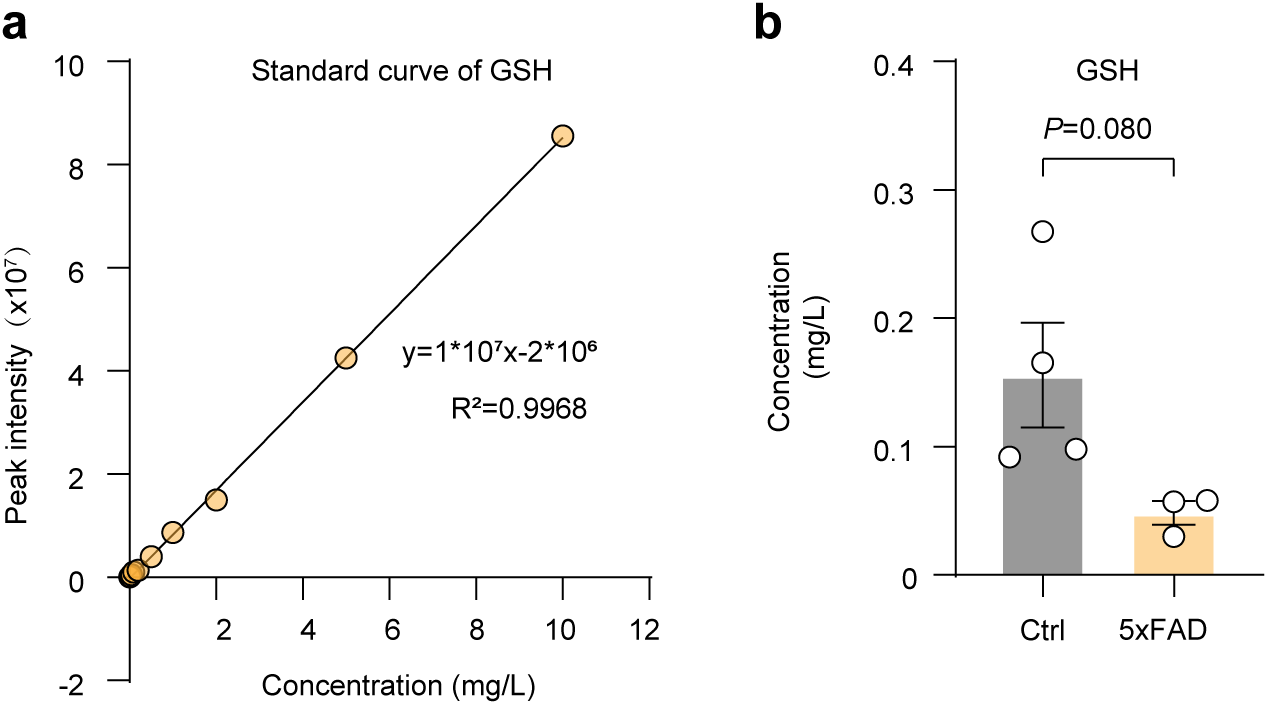
Targeted metabolomic validation of AD-associated decline in microglial GSH metabolism. (a) Standard curve for absolute quantification of GSH in targeted metabolomics in 5xFAD mice. (b) Absolute GSH levels in microglia from 12-month-old wild-type (n = 4 samples) and 5xFAD (n = 3 samples) mice (∼60,000 cells per sample). Data are presented as the mean ± SEM; two-tailed unpaired *t*-test; Ctrl, control. GSH, glutathione (reduced).

**Supplementary Fig. 9.**
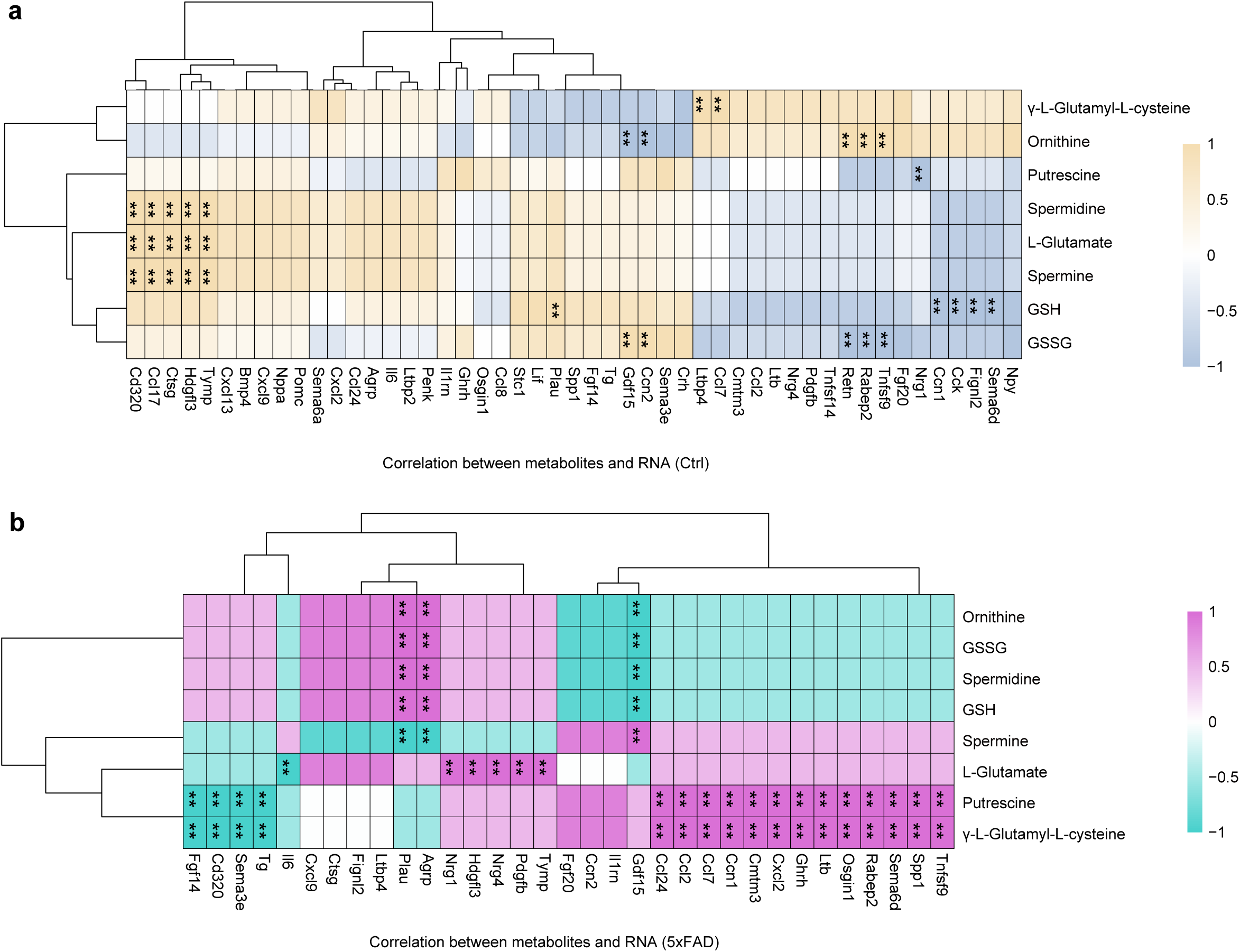
Correlation between glutathione-related metabolites and cytokine gene expression in microglia. **(a)** A heatmap of Spearman’s rank correlation analysis between the identified metabolites in glutathione metabolism pathway and the cytokine-related gene expression in microglia in control group. Yellow indicates positive correlation and blue denotes negative correlation. Data are presented as the mean ± SEM; two-tailed unpaired *t*-test; \*\**P* < 0.01; n.s., no significant difference. **(b)** A heatmap of Spearman’s rank correlation analysis between the identified metabolites in glutathione metabolism pathway and the cytokine-related gene expression in microglia in 5xFAD group. Purple indicates positive correlation and green denotes negative correlation. Data are presented as the mean ± SEM; two-tailed unpaired *t*-test; \*\**P* < 0.01; n.s., no significant difference.

**Supplementary Fig. 10.**
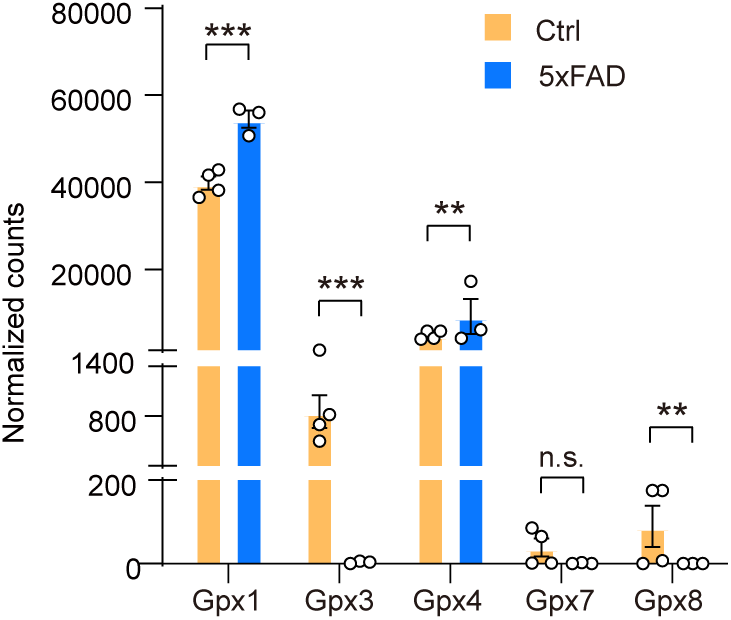
Comparisons of gene expression in microglia between the 5xFAD group and control group. Comparisons of gene expression in the Gpx family of microglia between the 5xFAD group (n = 3 samples) and control group (n = 4 samples). Data are presented as the mean ± SEM; two-tailed unpaired t-test; \*\**P* < 0.01, \*\*\**P* < 0.001; n.s., no significant difference.

**Supplementary Fig. 11.**
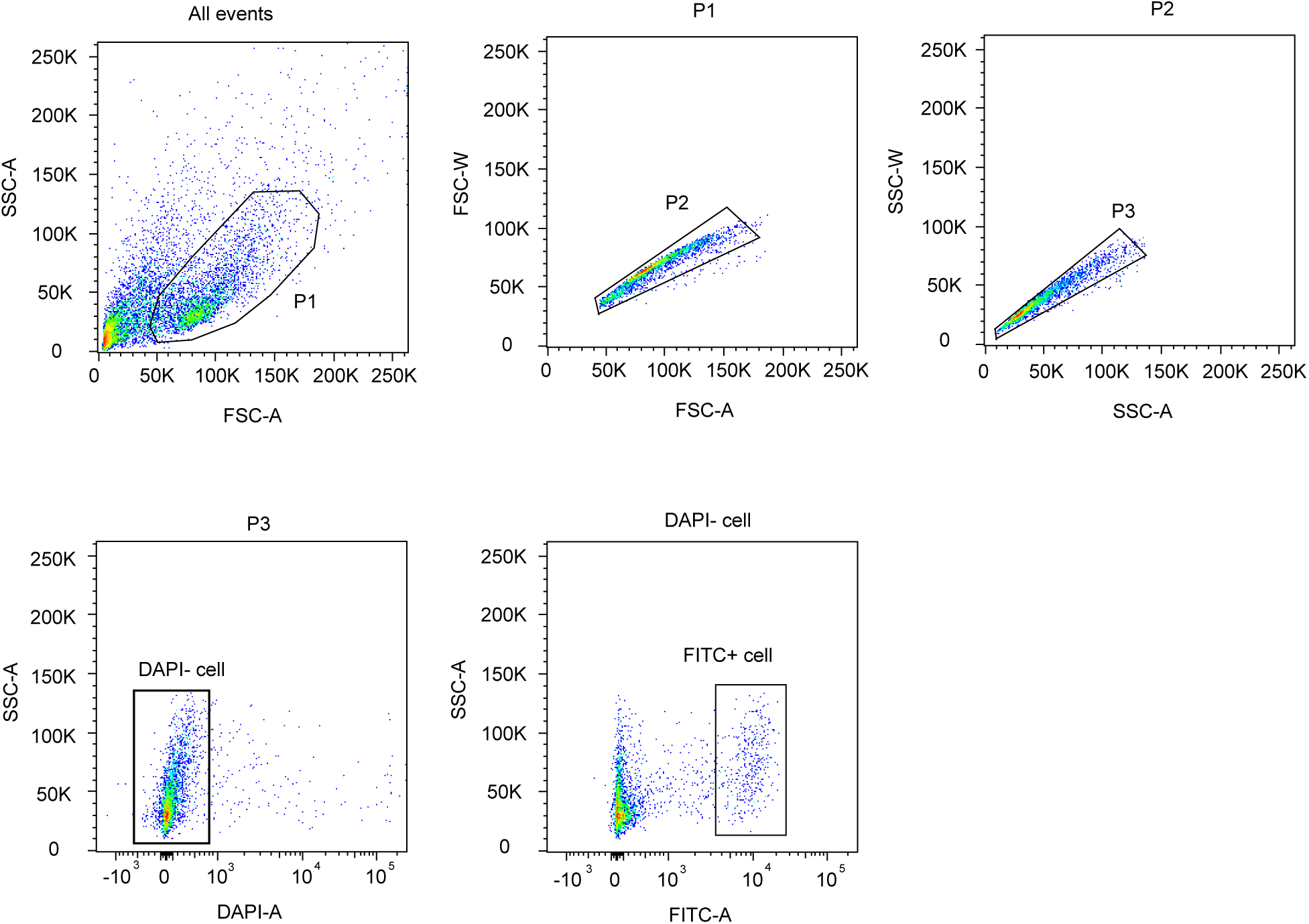
Flow cytometric gating strategy for isolation of three brain cell types. Representative gating strategy is shown. All events were first gated based on forward scatter area (FSC-A) and side scatter area (SSC-A) to exclude debris. P1 and P2 gates were applied sequentially using FSC-A/FSC-W and SSC-A/SSC-W to select single cells and exclude doublets. W, width. P3 cells were further gated to exclude dead cells based on DAPI staining (DAPI⁻ cells). Finally, FITC-positive cells were selected for downstream analyses.

## Methods

### Animals

All animal experiments were conducted in accordance with protocols approved by the Institutional Animal Care and Use Committee at Chinese Institute for Brain Research, Beijing (CIBR) (Number: LARC-T025). *Thy1^YFP^tg* (stock no.: 003709, n = 10), *Aldh1l1^EGFP^tg* (stock no.: 030247, n = 7), *Cx3cr1^+/EGFP^* (stock no.: 005582, n = 9), 5xFAD (Tg6799, stock no.: 006554, n = 6), *Cx3cr1-CreER* (stock no.: 021160, n = 11), *DTA* (stock no. 009669) mice were purchased from the Jackson Laboratory. *Gpx1* knockout mice (n = 8) were developed with assistance from Cyagen Biosciences Co., Ltd. Genotyping was performed with the following primers: 5’-TCTGAGTGGCAAAGGACCTTA GG-3’ (forward) and 5’-CGCTGAACTTGTGGCCGTTTACG-3’ (reverse) for *Thy1^YFP^tg* mice; 5’-CTGTCATCAGTCTTCACATCCAC-3’ (forward) and 5’-CTGAACTTGTGGCCGTTTAC-3’ (reverse) for *Aldh1l1^EGFP^tg* mice; 5’-CTCCCCCTGAACCTGAAAC-3’ (forward) and 5’-CCCAGACACTCGTTGTCCTT-3’ (reverse) for *Cx3cr1^+/EGFP^* mice; 5’-CGGGCCTCTTCGCTATTAC-3’, 5’-ACCCCCATGTCAGAGTTCCT-3’, and 5’-TATACAACCTTGGGGGATGG-3’ for 5xFAD mice. 5’-TTACACAATATAAGGGAGCTGTGC-3’ (forward), 5’-TTAAGGCACTGAGTAGCAGTGTG-3’ (reverse), 5’-TTACACAATATAAGGGAGCTGTGC-3’ (forward) and 5’-ATACCTGGTGTCCGAACTGATTG-3’ (reverse) for *Gpx1^+/–^* mice. 5’-AAGACTCACGTGGACCTGCT-3’ (forward), 5’-AGGATGTTGACT TCCGAGTTG-3’ (reverse), 5’-AAGACTCACGTGGACCTGCT-3’ (forward) (forward) and 5’-CGGTTATTCAACTTGCACCA-3’ (reverse) for *Cx3cr1-CreER* mice. 5’-CCAAAGTCGCTCTGAGTTGTTATC-3’ (forward), 5’-GAGCGGGAGAAATGGATATG-3’ (reverse), 5’- CGACCTGCAGGTCCTCG-3’ (forward) and 5’-CGACCTGCAGGTCCTCG-3’ (reverse) for *DTA* mice. C57BL/6 mice were purchased from Beijing Vital River Laboratory Animal Technology Co., Ltd. For Alzheimer’s disease model experiments, 5xFAD mice were studied at 12 months of age to capture established pathology. The others were used at 35–45 days of age to ensure consistency across astrocyte, microglia, and neuron comparisons. In our study, the majority of the animals used for cell isolation and downstream analyses (including metabolomic profiling of microglia, astrocytes, and neurons) were male mice. A small number of female mice (1–2 per cell type) were included in the sorting process, primarily due to availability constraints at the time of experimentation (e.g., aging mice). However, the number of female samples per cell type was below the generally accepted minimum for statistical comparison (n ≥ 3), particularly in the context of metabolomic analysis where biological variability is high and sex-stratified analysis typically requires larger cohorts to yield meaningful insights. Therefore, we did not perform sex-specific comparisons in this study. All animals were housed in the facility of the Animal Resource Center at CIBR under a light/dark cycle (12/12 h) with food and water *ad libitum*.

### Microglial ablation

To ablate brain microglia, we fed mice (4 weeks old) a diet formulated with 2 g PLX3397 (MCE)/kg diet (formulated by Jiangsu Xietong Pharmaceutical Bio-Engineering, China) for 14 d *ad libitum* as described^101^. The control group of mice was fed *ad libitum* with the same diet without the addition of PLX3397.

When we used a genetic approach, *Cx3cr1-CreER;DTA* to deplete microglia. Because microglia can rapidly repopulate after depletion, we optimized the tamoxifen regimen by injecting it for 3 or 5 consecutive days and examining microglial abundance 1 day (24 h), 2 days (48 h), and 7 days after the final injection. Immunohistochemical staining confirmed that 5 consecutive days of tamoxifen treatment resulted in near-complete depletion of microglia 48 h later, with microglia repopulation by day 7 after the cessation of treatment. We therefore used this optimized condition, 5-day tamoxifen induction with 48 h recovery, for subsequent metabolomic analyses.

### Tissue preparation for histological experiments

Mice were anesthetized with avertin (100 mg/kg of body weight; cat no. 75-80-9, Sigma). Animals were sequentially perfused with phosphate-buffered saline (PBS) and 4% paraformaldehyde in PBS. Brains were harvested, followed by overnight post-fixation in 4% paraformaldehyde at 4°C. The brains were then dehydrated in 30% sucrose in PBS at 4°C for 2 d. After being embedded in optimal cutting temperature compound (OCT, Tissue-Tek), the brain samples were frozen before sectioning (40 μm thick) with a cryostat (CM3050S, Leica)^102,103^.

### Immunohistochemistry

Brain sections were rinsed with PBS three times for 10 min each, followed by incubation with a blocking solution containing 4% normal goat serum (C0265, Beyotime). The samples were then treated with 0.3% Triton X-100 (T8787, Sigma) in 3% bovine serum albumin (BSA; Beyotime) at room temperature for 2 h. Samples were incubated with primary antibodies with blocking solution at 4°C overnight. After being rinsed with PBS three times, samples were reacted with secondary antibodies and Hoechst 33342 (1:1,000; 62249, Thermo Scientific) at room temperature for 2 h. Samples were rinsed three times with PBS before being mounted for subsequent imaging as previously reported^103^. Immunostaining results from Fig. 1a, 1b, 2a, 4b, 8l, 8o and Extended Data Fig. 2a, 2b, 3a, 3b, and 7b were independently repeated at least three times (from three mice). Sholl analysis of reconstructed microglia was performed in the *knockout* group and control group. Analysis was performed by placing a series of concentric circles spaced at 2-μm intervals and centered on the soma.

### Astrocyte-specific labeling in mice using systemic AAV delivery

For astrocyte labeling, mice were injected with the adeno-associated viral vector AAV2/PHP.eB-gfaABC1D-mRuby3-WPRE-pA (Shanghai Taitool Bioscience Co., Ltd) via retro-orbital injection. Each mouse received a total viral dose of 2 × 10¹¹ genome copies^104^. Viral expression was allowed to proceed for 3 weeks before subsequent experiments were performed.

### Tissue preparation for metabolomics

Mice were anesthetized with avertin. Animals were then perfused with cold PBS. Brains were quickly harvested and frozen in liquid nitrogen and stored at –80°C. Before the samples were prepared, the brains were rewarmed at –20°C for 2 h. The region of interest used for metabolomic profiling was a ∼1 mm-thick cortical region located above the hippocampus, corresponding approximately to coronal sections spanning from Bregma −1.46 mm to −2.46 mm, based on the Paxinos and Franklin mouse brain atlas (4th edition). Tissue was collected from a similar position of the cerebral cortex from each brain. After being weighed with a precision balance, all samples were stored at –80℃ until metabolite extraction.

### Cell isolation

Mice at postnatal day (P)30–45 were anesthetized with avertin. We chose avertin because of its rapid onset, reproducible depth of anesthesia, and minimal interference with core physiological and metabolic parameters, which is particularly important for metabolomic studies. Previous studies have shown that tribromoethanol (TBE, i.e., avertin) produces stable anesthesia with no significant alterations in arterial blood gas (pCO₂, pO₂), pH, mean arterial pressure, heart rate, SpO₂ (peripheral oxygen saturation), or body temperature during the anesthetic period, and does not induce histological or biochemical damage to major organs, even at high doses^105^. They were then perfused with cold PBS through the heart. Brains were isolated and immediately immersed in cold hibernation medium (5 mM KCl, 5 mM NaOH, 5 mM NaH_2_PO_4_, 0.5 mM MgCl_2_, 20 mM Na pyruvate, 5.5 mM glucose, 200 mM sorbitol, pH 7.2–7.4; filtered with a 0.22-μm filter; BrainBits). Dissected cerebral cortex and hippocampus regions from isolated brains were cut into small pieces and incubated in 15-ml tubes containing cold hibernation medium with papain (2 mg/ml; Sigma) and DNase I (100 U**/**ml; Sigma) at 37°C for 30 min with intermittent flicking of the tubes. The tissue was triturated into an individual cell suspension with a siliconized Pasteur pipette with a fire-polished tip (BrainBits). Cell suspensions were added to an Optiprep gradient ^106^ and centrifuged at 900*g* for 15 min at 4°C. We then removed and discarded the uppermost 5 ml of the gradient (which contains debris) and collected specific Optiprep layers (neurons and astrocytes are enriched in upper layers, whereas microglia are enriched in pellets; Brewer and Torricelli, 2007a). Cells from the collected layers were washed with cold hibernation medium without Ca^2+^ and Mg^2+^ (BrainBits) and centrifuged at 600*g* for 10 min at 4°C. Cell pellets were resuspended with 500 µl cold hibernation medium for sorting.

### Fluorescence-activated cell sorting (FACS)

FACS was performed on a BD FACS Aria II sorter using a 100-μm nozzle at 13.1 psi. Cell gates were defined as follows: neurons, YFP^+^DAPI^−^; control for neurons, YFP^−^DAPI^−^; astrocytes, EGFP^+^DAPI^−^; control for astrocytes, EGFP^−^DAPI^−^; microglia, EGFP^+^DAPI^−^; control for microglia, EGFP^−^DAPI^−^. Cells were sorted into 200 µl PBS and then spun down at 1,600*g* at 4°C for 8 min. After washing the cells with PBS and centrifuging them again, we resuspended the cells in 80% methanol/20% water (vol/vol) or 80% acetonitrile/20% water (vol/vol). Fluorescent Anti-CD11b Rat Monoclonal Antibody (Brilliant Violet 421; 101236, BioLegend) was used for sorting microglia from the cortices of aging mice and 5xFAD and *Gpx*^−/–^ mice (**Supplementary Fig. 11**). From each adult mouse, approximately 50,000 microglia were isolated to establish a single sample for targeted or untargeted metabolomic analysis.

### Targeted metabolomic analysis

The cell pellet was added to 800 µl of ice-cold 80% methanol/20% water for metabolite extraction. After vigorous agitation, samples were centrifuged at 15,000*g* for 15 min at 4°C. The metabolite-containing supernatant was dried in a SpeedVac concentrator (35°C, vacuum pressure: 5.1 mTorr, Thermo Savant) for 5–7 h. Targeted metabolomic measurements were performed as described ^28,107,108^. Briefly, metabolites were reconstituted in 100 μl of 0.03% formic acid in water. The samples were vortexed and centrifuged to remove debris. LC-MS (Liquid chromatography–mass spectrometry) and data acquisition were performed with an AB QTRAP 5500 liquid chromatography/triple quadrupole mass spectrometer (Applied Biosystems SCIEX) with an injection volume of 20 μl. Separation was achieved on a Phenomenex Synergi Polar-RP HPLC column (150 × 2 mm, 4 μm, 80 Å) using a Nexera Ultra High-Performance Liquid Chromatograph (UHPLC) system (Shimadzu Corporation, Kyoto, Japan). The mobile phases employed were 0.03% formic acid in water (A) and 0.03% formic acid in acetonitrile (B). The column was maintained at 35°C and the samples were kept in the autosampler at 4°C. The flow rate was 0.5 ml/min, and injection volume 20 μl. The mass spectrometer was used with the electrospray ionization (ESI) source in multiple reaction monitoring (MRM) mode. The MRM MS/MS detector conditions were set as follows: curtain gas 30 psi; ion spray voltages 1200 V (positive) and -1500 V (negative); temperature 650°C; ion source gas 1 with 50 psi; ion source gas 2 with 50 psi; interface heater on; entrance potential 10 V. Dwell time for each transition was set at 3 msec. Samples were analyzed in a randomized order, and MRM data was acquired using Analyst 1.6.3 software. Chromatogram review and peak area integration were performed using MultiQuant software version 2.1 (Applied Biosystems SCIEX). The peak area of a detected metabolite was normalized against the total ion counts of each sample and the mean for each metabolite within the sample batch of each run to correct for variations introduced by sample handling.

### Untargeted metabolomic measurement and data analysis

The cell pellet from tissue or isolated brain cells was added to 300 µl of ice-cold 80% acetonitrile/20% water for metabolite extraction. Brain tissues were added to 400 µl of ice-cold 80% acetonitrile/20% water and were homogenized with a grinder (1 ml, Wheaton Dounce) for metabolite extraction. After mixing and centrifuging, the metabolite-containing supernatant was dried in a SpeedVac concentrator as described above for 3 h. The cell pellets were dissolved in 35 µl of 80% acetonitrile/20% water. The brain slice samples were dissolved in a volume of 80% acetonitrile/20% water according to their initial tissue weight (100 µl/mg brain tissue). Untargeted metabolomic measurements were performed on a Q Exactive mass spectrometer (Thermo Scientific) as reported^4,29^.

The HILIC method used an Agilent Infinity Lab Poroshell 120 HILIC-Z column (2.1 × 100 mm, 2.7 μm) with a PEEK liner. Each sample (in a volume of 5 µl) was injected into the column. For the positive ion mode, mobile phase A was water containing 10 mM ammonium formate, and mobile phase B was acetonitrile containing 10 mM ammonium formate. For the negative ion mode, mobile phase A was water containing 10 mM ammonium acetate, and mobile phase B was acetonitrile containing 10 mM ammonium acetate. Gradient separation proceeded as follows: from 0 to 4 min, mobile phase B was ramped linearly from 100% to 84%; from 4 min to 11 min, mobile phase B was ramped linearly from 84% to 40%; from 11 min to 12 min, mobile phase B was held at 40%; from 12 min to 13 min, mobile phase B was ramped linearly from 40% to 100%; mobile phase B was then held at 100% from 13 min to 17 min to regenerate the initial chromatographic environment. The solvent flow rate was kept steady throughout at 400 μl/minute, and the column temperature was maintained at 35°C. For the MS1 method, the resolution was 70,000 in full-scan MS. The automatic gain control (AGC) target was set to 3,000,000 with a maximum injection time of 100 ms. The scan range was set to 60–900 *m/z*. MS2 was only used to pool the sample. The resolution was 17,500, and the AGC target was set to 50,000 with a maximum injection time of 80 ms. A top-10 data-dependent MS scheme was used with an isolation window of 1.6 *m/z*. Analytes were fragmented with stepped collision energies of 20, 30, and 40 normalized collision energy (NCE) units. The minimum AGC target was 1,000 with a dynamic exclusion of 6 s. The scan range was set to 60–400 *m/z* and 350–900 *m/z*^4^.

Metabolomic analysis detected a total of 9,854 metabolite features. They were annotated from the mzCloud libraries (https://www.mzcloud.org/). The identified metabolites were classified using the Human Metabolome Database (HMDB) (https://www.hmdb.ca/), resulting in the successful categorization of 1,782 metabolites.

### RNA-seq library construction and sequencing

Total RNA was extracted from brain cells using a RNeasy Micro kit (cat no. 74004, QIAGEN). RNA quality was evaluated with an Agilent RNA 6000 Pico kit. We used samples with The RNA integrity number (RIN) ≥ 8 for cDNA library preparation. The cDNA libraries were constructed using SMARTer Stranded Total RNA-Seq kit v2 (cat no. 634411, Takara). Preliminary quantification of the library was assessed with Qubit 4.0 (Thermo Fisher Scientific). An Agilent 2100 Bioanalyzer was used to detect DNA fragment integrity and insert fragment size. High-quality libraries were sequenced with an Illumina NovaSeq 6000 sequencer to obtain 150-bp paired-end reads.

For the preprocessing of bulk RNA-seq, we carried out read quality control using FastQC with the paired-end parameter ^109^. Then, the trimming and filtering procedure was implemented using fastp ^110^. We ran STAR to map the filtered reads to the mouse genome and quantify the mapped gene number ^111^. To further compare different cell types at the mRNA level, we used the transcripts per kilobase million (TPM) method to normalize gene counts and isolated differentially expressed genes with DESeq2 ^112^.

### Pattern recognition analysis and statistical analysis

MultiQuant software (AB SCIEX) was used to integrate chromatogram peaks of each metabolite detected by targeted metabolomics. A total ion chromatogram was used to normalize single ion values against the total ion value of the entire chromatogram, and the ion intensity was calculated as described ^28^. Compound Discoverer 3.2 (Thermo Fisher Scientific) and XCMS^Plus^ were used to analyze data from untargeted metabolomics. The mass-to-charge ratio of metabolites was calculated with Xcalibur software ^113,114^. For the unidentified metabolites, they often correspond to multiple possible chemical structures, so classification without definitive structural identification would be speculative and potentially misleading. Moreover, due to the limited quantity of starting material, the MS/MS fragmentation data necessary for reliable structural annotation could not be acquired for these specific features. The low abundance of ions did not meet the threshold required for confident MS/MS acquisition, resulting in insufficient fragment of ion data. Thus, we represent them using *m/z*. For both targeted and untargeted metabolomics, raw LC-MS data were processed to generate peak intensity tables, which were normalized by the sum of intensities across all detected features per sample prior to downstream analyses. Pathway analysis and unsupervised and supervised multivariate data analysis approaches, including principal component analysis (PCA), hierarchical clustering, and partial least squares discriminant analysis (PLS-DA), were performed using Metaboanalyst 5.0 ^115^. Feature selection in PLS-DA was used to identify metabolites that maximized the separation between targeted cells and control groups. The variable importance in projection (VIP) of each variable in the PLS-DA model was calculated to determine its contribution to the classification. By convention, a VIP score > 1 is used as a criterion for variable selection ^116–118^. Thus, metabolites with a VIP score > 1 were reported. Comparison of gene expression and pathway analyses of differentially expressed genes were performed using R software. NIH ImageJ software was used to quantify the imaging data obtained from confocal microscopy. Statistical analyses were performed by SPSS version 23.0. Continuous variables are presented as the mean ± standard error of the mean (SEM). The statistical significance (raw *P* < 0.05) of differences in gene expression and metabolite levels was assessed using a two-tailed Student’s t-test for comparing two groups or a one-way analysis of variance (ANOVA) followed by a post hoc Tukey’s multiple comparisons test for comparing three groups, where appropriate.

## Data availability

Data generated and/or analyzed in the current study are available from the website (https://metabolismocean.org/braincell), or from the corresponding authors upon reasonable request. Sequencing data that support the findings of this study have been deposited in the National Center for Biotechnology Information GSE316928. There are no restrictions on data availability. Data supporting the findings of this study are available from the lead contact on reasonable request. Source data are provided with this paper.

## References

1 Somel, M., Liu, X. & Khaitovich, P. Human brain evolution: transcripts, metabolites and their regulators. Nat Rev Neurosci 14, 112–127, doi:10.1038/nrn3372 (2013).

2 Belanger, M., Allaman, I. & Magistretti, P. J. Brain energy metabolism: focus on astrocyte-neuron metabolic cooperation. Cell Metab 14, 724–738, doi:10.1016/j.cmet.2011.08.016 (2011).

3 Magistretti, P. J. & Allaman, I. A cellular perspective on brain energy metabolism and functional imaging. Neuron 86, 883–901, doi:10.1016/j.neuron.2015.03.035 (2015).

4 Wang, Y., Zhou, L., Wang, N., Qiu, B., Yao, D., Yu, J., He, M., Li, T., Xie, Y., Yu, X., Bi, Z., Sun, X., Ji, X., Li, Z., Mo, D. & Ge, W. P. Comprehensive characterization of metabolic consumption and production by the human brain. Neuron 113, 1708–1722 e1705, doi:10.1016/j.neuron.2025.03.003 (2025).

5 Araque, A. & Navarrete, M. Glial cells in neuronal network function. Philos Trans R Soc Lond B Biol Sci 365, 2375–2381, doi:10.1098/rstb.2009.0313 (2010).

6 Zhang, Z. G., Bower, L., Zhang, R. L., Chen, S., Windham, J. P. & Chopp, M. Three-dimensional measurement of cerebral microvascular plasma perfusion, glial fibrillary acidic protein and microtubule associated protein-2 immunoreactivity after embolic stroke in rats: a double fluorescent labeled laser-scanning confocal microscopic study. Brain Res 844, 55–66, doi:10.1016/s0006-8993(99)01886-7 (1999).

7 Simard, M., Arcuino, G., Takano, T., Liu, Q. S. & Nedergaard, M. Signaling at the gliovascular interface. J Neurosci 23, 9254–9262 (2003).

8 Tsacopoulos, M. & Magistretti, P. J. Metabolic coupling between glia and neurons. J Neurosci 16, 877–885 (1996).

9 Simpson, I. A., Carruthers, A. & Vannucci, S. J. Supply and demand in cerebral energy metabolism: the role of nutrient transporters. J Cereb Blood Flow Metab 27, 1766–1791, doi:10.1038/sj.jcbfm.9600521 (2007).

10 Yang, A. C., Vest, R. T., Kern, F., Lee, D. P., Agam, M., Maat, C. A., Losada, P. M., Chen, M. B., Schaum, N., Khoury, N., Toland, A., Calcuttawala, K., Shin, H., Palovics, R., Shin, A., Wang, E. Y., Luo, J., Gate, D., Schulz-Schaeffer, W. J., Chu, P., Siegenthaler, J. A., McNerney, M. W., Keller, A. & Wyss-Coray, T. A human brain vascular atlas reveals diverse mediators of Alzheimer’s risk. Nature 603, 885–892, doi:10.1038/s41586-021-04369-3 (2022).

11 Salter, M. W. & Stevens, B. Microglia emerge as central players in brain disease. Nat Med 23, 1018–1027, doi:10.1038/nm.4397 (2017).

12 Liu, Y. U., Ying, Y., Li, Y., Eyo, U. B., Chen, T., Zheng, J., Umpierre, A. D., Zhu, J., Bosco, D. B., Dong, H. & Wu, L. J. Neuronal network activity controls microglial process surveillance in awake mice via norepinephrine signaling. Nat Neurosci 22, 1771–1781, doi:10.1038/s41593-019-0511-3 (2019).

13 Song, W. M. & Colonna, M. The identity and function of microglia in neurodegeneration. Nat Immunol 19, 1048–1058, doi:10.1038/s41590-018-0212-1 (2018).

14 Distefano-Gagne, F., Bitarafan, S., Lacroix, S. & Gosselin, D. Roles and regulation of microglia activity in multiple sclerosis: insights from animal models. Nat Rev Neurosci 24, 397–415, doi:10.1038/s41583-023-00709-6 (2023).

15 Pocock, J. M. & Piers, T. M. Modelling microglial function with induced pluripotent stem cells: an update. Nat Rev Neurosci 19, 445–452, doi:10.1038/s41583-018-0030-3 (2018).

16 Borst, K., Dumas, A. A. & Prinz, M. Microglia: Immune and non-immune functions. Immunity 54, 2194–2208, doi:10.1016/j.immuni.2021.09.014 (2021).

17 Wu, J., Wang, Y., Li, X., Ouyang, P., Cai, Y., He, Y., Zhang, M., Luan, X., Jin, Y., Wang, J., Xiao, Y., Liang, Y., Xie, F., Shu, Y., Hu, J., Chang, C., Jiang, J., Wu, D., Zhao, Y., Liu, T., Li, Y., Huang, X., Li, Y., Zhang, J., Cao, Y., Cheng, X., Mao, Y., Rao, Y., Cao, L. & Peng, B. Microglia replacement halts the progression of microgliopathy in mice and humans. Science 389, eadr1015, doi:10.1126/science.adr1015 (2025).

18 Marschallinger, J., Iram, T., Zardeneta, M., Lee, S. E., Lehallier, B., Haney, M. S., Pluvinage, J. V., Mathur, V., Hahn, O., Morgens, D. W., Kim, J., Tevini, J., Felder, T. K., Wolinski, H., Bertozzi, C. R., Bassik, M. C., Aigner, L. & Wyss-Coray, T. Lipid-droplet-accumulating microglia represent a dysfunctional and proinflammatory state in the aging brain. Nat Neurosci 23, 194–208, doi:10.1038/s41593-019-0566-1 (2020).

19 Badimon, A., Strasburger, H. J., Ayata, P., Chen, X., Nair, A., Ikegami, A., Hwang, P., Chan, A. T., Graves, S. M., Uweru, J. O., Ledderose, C., Kutlu, M. G., Wheeler, M. A., Kahan, A., Ishikawa, M., Wang, Y. C., Loh, Y. E., Jiang, J. X., Surmeier, D. J., Robson, S. C., Junger, W. G., Sebra, R., Calipari, E. S., Kenny, P. J., Eyo, U. B., Colonna, M., Quintana, F. J., Wake, H., Gradinaru, V. & Schaefer, A. Negative feedback control of neuronal activity by microglia. Nature 586, 417–423, doi:10.1038/s41586-020-2777-8 (2020).

20 Consortium, H. D. i. Bioenergetic deficits in Huntington’s disease iPSC-derived neural cells and rescue with glycolytic metabolites. Hum Mol Genet 29, 1757–1771, doi:10.1093/hmg/ddy430 (2020).

21 Cunnane, S. C., Trushina, E., Morland, C., Prigione, A., Casadesus, G., Andrews, Z. B., Beal, M. F., Bergersen, L. H., Brinton, R. D., de la Monte, S., Eckert, A., Harvey, J., Jeggo, R., Jhamandas, J. H., Kann, O., la Cour, C. M., Martin, W. F., Mithieux, G., Moreira, P. I., Murphy, M. P., Nave, K. A., Nuriel, T., Oliet, S. H. R., Saudou, F., Mattson, M. P., Swerdlow, R. H. & Millan, M. J. Brain energy rescue: an emerging therapeutic concept for neurodegenerative disorders of ageing. Nat Rev Drug Discov 19, 609–633, doi:10.1038/s41573-020-0072-x (2020).

22 Duscha, A., Gisevius, B., Hirschberg, S., Yissachar, N., Stangl, G. I., Eilers, E., Bader, V., Haase, S., Kaisler, J., David, C., Schneider, R., Troisi, R., Zent, D., Hegelmaier, T., Dokalis, N., Gerstein, S., Del Mare-Roumani, S., Amidror, S., Staszewski, O., Poschmann, G., Stuhler, K., Hirche, F., Balogh, A., Kempa, S., Trager, P., Zaiss, M. M., Holm, J. B., Massa, M. G., Nielsen, H. B., Faissner, A., Lukas, C., Gatermann, S. G., Scholz, M., Przuntek, H., Prinz, M., Forslund, S. K., Winklhofer, K. F., Muller, D. N., Linker, R. A., Gold, R. & Haghikia, A. Propionic Acid Shapes the Multiple Sclerosis Disease Course by an Immunomodulatory Mechanism. Cell 180, 1067–1080 e1016, doi:10.1016/j.cell.2020.02.035 (2020).

23 Sada, N., Lee, S., Katsu, T., Otsuki, T. & Inoue, T. Epilepsy treatment. Targeting LDH enzymes with a stiripentol analog to treat epilepsy. Science 347, 1362–1367, doi:10.1126/science.aaa1299 (2015).

24 Morrison, B. M. Surprising New Players in Glia-Neuron Crosstalk: Role in CNS Regeneration. Cell Metab 32, 695–696, doi:10.1016/j.cmet.2020.10.009 (2020).

25 Bartels, T., De Schepper, S. & Hong, S. Microglia modulate neurodegeneration in Alzheimer’s and Parkinson’s diseases. Science 370, 66–69, doi:10.1126/science.abb8587 (2020).

26 Filipello, F., Morini, R., Corradini, I., Zerbi, V., Canzi, A., Michalski, B., Erreni, M., Markicevic, M., Starvaggi-Cucuzza, C., Otero, K., Piccio, L., Cignarella, F., Perrucci, F., Tamborini, M., Genua, M., Rajendran, L., Menna, E., Vetrano, S., Fahnestock, M., Paolicelli, R. C. & Matteoli, M. The Microglial Innate Immune Receptor TREM2 Is Required for Synapse Elimination and Normal Brain Connectivity. Immunity 48, 979–991 e978, doi:10.1016/j.immuni.2018.04.016 (2018).

27 Savini, M., Folick, A., Lee, Y. T., Jin, F., Cuevas, A., Tillman, M. C., Duffy, J. D., Zhao, Q., Neve, I. A., Hu, P. W., Yu, Y., Zhang, Q., Ye, Y., Mair, W. B., Wang, J., Han, L., Ortlund, E. A. & Wang, M. C. Lysosome lipid signalling from the periphery to neurons regulates longevity. Nat Cell Biol, doi:10.1038/s41556-022-00926-8 (2022).

28 Yu, J., Meng, F., X., He, F., P., Chen, F., Bao, W., X., Yu, Y., M., Zhou, J., T., Gao, J., Li, J., Q., Yao, Y., Ge, W., P. & Luo, B., Y. Metabolic Abnormalities in Patients with Chronic Disorders of Consciousness. Aging and disease 12, doi:10.14336/AD.12020.10812 (2020) (2020).

29 He, W., Gao, S., Du, L., Han, T., Yao, D., Wu, H., Li, Q., Li, F., Ge, W. P. & Wang, Y. Cisternostomy Facilitates Clearance of Metabolic Waste from Cerebrospinal Fluid in Patients with Severe Brain Injury. Aging Dis, doi:10.14336/AD.2025.0738 (2025).

30 Hu, Y., Cao, K., Wang, F., Wu, W., Mai, W., Qiu, L., Luo, Y., Ge, W. P., Sun, B., Shi, L., Zhu, J., Zhang, J., Wu, Z., Xie, Y., Duan, S. & Gao, Z. Dual roles of hexokinase 2 in shaping microglial function by gating glycolytic flux and mitochondrial activity. Nat Metab 4, 1756–1774, doi:10.1038/s42255-022-00707-5 (2022).

31 Ulland, T. K., Song, W. M., Huang, S. C., Ulrich, J. D., Sergushichev, A., Beatty, W. L., Loboda, A. A., Zhou, Y., Cairns, N. J., Kambal, A., Loginicheva, E., Gilfillan, S., Cella, M., Virgin, H. W., Unanue, E. R., Wang, Y., Artyomov, M. N., Holtzman, D. M. & Colonna, M. TREM2 Maintains Microglial Metabolic Fitness in Alzheimer’s Disease. Cell 170, 649–663 e613, doi:10.1016/j.cell.2017.07.023 (2017).

32 Ulland, T. K. & Colonna, M. TREM2 - a key player in microglial biology and Alzheimer disease. Nat Rev Neurol 14, 667–675, doi:10.1038/s41582-018-0072-1 (2018).

33 Nugent, A. A., Lin, K., van Lengerich, B., Lianoglou, S., Przybyla, L., Davis, S. S., Llapashtica, C., Wang, J., Kim, D. J., Xia, D., Lucas, A., Baskaran, S., Haddick, P. C. G., Lenser, M., Earr, T. K., Shi, J., Dugas, J. C., Andreone, B. J., Logan, T., Solanoy, H. O., Chen, H., Srivastava, A., Poda, S. B., Sanchez, P. E., Watts, R. J., Sandmann, T., Astarita, G., Lewcock, J. W., Monroe, K. M. & Di Paolo, G. TREM2 Regulates Microglial Cholesterol Metabolism upon Chronic Phagocytic Challenge. Neuron 105, 837–854 e839, doi:10.1016/j.neuron.2019.12.007 (2020).

34 Hammer, J., Alvestad, S., Osen, K. K., Skare, O., Sonnewald, U. & Ottersen, O. P. Expression of glutamine synthetase and glutamate dehydrogenase in the latent phase and chronic phase in the kainate model of temporal lobe epilepsy. Glia 56, 856–868, doi:10.1002/glia.20659 (2008).

35 Coulter, D. A. & Eid, T. Astrocytic regulation of glutamate homeostasis in epilepsy. Glia 60, 1215–1226, doi:10.1002/glia.22341 (2012).

36 Liang, S. L., Carlson, G. C. & Coulter, D. A. Dynamic regulation of synaptic GABA release by the glutamate-glutamine cycle in hippocampal area CA1. J Neurosci 26, 8537–8548, doi:10.1523/JNEUROSCI.0329-06.2006 (2006).

37 Kaddurah-Daouk, R. & Krishnan, K. R. Metabolomics: a global biochemical approach to the study of central nervous system diseases. Neuropsychopharmacology 34, 173–186, doi:10.1038/npp.2008.174 (2009).

38 Ivanisevic, J. & Siuzdak, G. The Role of Metabolomics in Brain Metabolism Research. J Neuroimmune Pharmacol 10, 391–395, doi:10.1007/s11481-015-9621-1 (2015).

39 Ivanisevic, J., Epstein, A. A., Kurczy, M. E., Benton, P. H., Uritboonthai, W., Fox, H. S., Boska, M. D., Gendelman, H. E. & Siuzdak, G. Brain region mapping using global metabolomics. Chem Biol 21, 1575–1584, doi:10.1016/j.chembiol.2014.09.016 (2014).

40 Varma, V. R., Oommen, A. M., Varma, S., Casanova, R., An, Y., Andrews, R. M., O’Brien, R., Pletnikova, O., Troncoso, J. C., Toledo, J., Baillie, R., Arnold, M., Kastenmueller, G., Nho, K., Doraiswamy, P. M., Saykin, A. J., Kaddurah-Daouk, R., Legido-Quigley, C. & Thambisetty, M. Brain and blood metabolite signatures of pathology and progression in Alzheimer disease: A targeted metabolomics study. PLoS Med 15, e1002482, doi:10.1371/journal.pmed.1002482 (2018).

41 Filiou, M. D., Zhang, Y., Teplytska, L., Reckow, S., Gormanns, P., Maccarrone, G., Frank, E., Kessler, M. S., Hambsch, B., Nussbaumer, M., Bunck, M., Ludwig, T., Yassouridis, A., Holsboer, F., Landgraf, R. & Turck, C. W. Proteomics and metabolomics analysis of a trait anxiety mouse model reveals divergent mitochondrial pathways. Biol Psychiatry 70, 1074–1082, doi:10.1016/j.biopsych.2011.06.009 (2011).

42 Montaner, J., Ramiro, L., Simats, A., Tiedt, S., Makris, K., Jickling, G. C., Debette, S., Sanchez, J. C. & Bustamante, A. Multilevel omics for the discovery of biomarkers and therapeutic targets for stroke. Nat Rev Neurol 16, 247–264, doi:10.1038/s41582-020-0350-6 (2020).

43 Lee, J., d’Aigle, J., Atadja, L., Quaicoe, V., Honarpisheh, P., Ganesh, B. P., Hassan, A., Graf, J., Petrosino, J., Putluri, N., Zhu, L., Durgan, D. J., Bryan, R. M., Jr., McCullough, L. D. & Venna, V. R. Gut Microbiota-Derived Short-Chain Fatty Acids Promote Poststroke Recovery in Aged Mice. Circ Res 127, 453–465, doi:10.1161/CIRCRESAHA.119.316448 (2020).

44 Zhang, Y., Lu, W., Wang, Z., Zhang, R., Xie, Y., Guo, S., Jiao, L., Hong, Y., Di, Z., Wang, G. & Aa, J. Reduced Neuronal cAMP in the Nucleus Accumbens Damages Blood-Brain Barrier Integrity and Promotes Stress Vulnerability. Biol Psychiatry 87, 526–537, doi:10.1016/j.biopsych.2019.09.027 (2020).

45 Lan, M. J., McLoughlin, G. A., Griffin, J. L., Tsang, T. M., Huang, J. T., Yuan, P., Manji, H., Holmes, E. & Bahn, S. Metabonomic analysis identifies molecular changes associated with the pathophysiology and drug treatment of bipolar disorder. Mol Psychiatry 14, 269–279, doi:10.1038/sj.mp.4002130 (2009).

46 Machler, P., Wyss, M. T., Elsayed, M., Stobart, J., Gutierrez, R., von Faber-Castell, A., Kaelin, V., Zuend, M., San Martin, A., Romero-Gomez, I., Baeza-Lehnert, F., Lengacher, S., Schneider, B. L., Aebischer, P., Magistretti, P. J., Barros, L. F. & Weber, B. In Vivo Evidence for a Lactate Gradient from Astrocytes to Neurons. Cell Metab 23, 94–102, doi:10.1016/j.cmet.2015.10.010 (2016).

47 Beyer, B. A., Fang, M., Sadrian, B., Montenegro-Burke, J. R., Plaisted, W. C., Kok, B. P. C., Saez, E., Kondo, T., Siuzdak, G. & Lairson, L. L. Metabolomics-based discovery of a metabolite that enhances oligodendrocyte maturation. Nat Chem Biol 14, 22–28, doi:10.1038/nchembio.2517 (2018).

48 Haney, M. S., Palovics, R., Munson, C. N., Long, C., Johansson, P. K., Yip, O., Dong, W., Rawat, E., West, E., Schlachetzki, J. C. M., Tsai, A., Guldner, I. H., Lamichhane, B. S., Smith, A., Schaum, N., Calcuttawala, K., Shin, A., Wang, Y. H., Wang, C., Koutsodendris, N., Serrano, G. E., Beach, T. G., Reiman, E. M., Glass, C. K., Abu-Remaileh, M., Enejder, A., Huang, Y. & Wyss-Coray, T. APOE4/4 is linked to damaging lipid droplets in Alzheimer’s disease microglia. Nature 628, 154–161, doi:10.1038/s41586-024-07185-7 (2024).

49 Guma, M., Tiziani, S. & Firestein, G. S. Metabolomics in rheumatic diseases: desperately seeking biomarkers. Nat Rev Rheumatol 12, 269–281, doi:10.1038/nrrheum.2016.1 (2016).

50 Hempel, C. M., Sugino, K. & Nelson, S. B. A manual method for the purification of fluorescently labeled neurons from the mammalian brain. Nat Protoc 2, 2924–2929, doi:10.1038/nprot.2007.416 (2007).

51 Brewer, G. J. & Torricelli, J. R. Isolation and culture of adult neurons and neurospheres. Nat Protoc 2, 1490–1498, doi:10.1038/nprot.2007.207 (2007).

52 Barres, B. A. Designing and troubleshooting immunopanning protocols for purifying neural cells. Cold Spring Harb Protoc 2014, 1342–1347, doi:10.1101/pdb.ip073999 (2014).

53 Zhu, C., Wang, X., Huang, Z., Qiu, L., Xu, F., Vahsen, N., Nilsson, M., Eriksson, P. S., Hagberg, H., Culmsee, C., Plesnila, N., Kroemer, G. & Blomgren, K. Apoptosis-inducing factor is a major contributor to neuronal loss induced by neonatal cerebral hypoxia-ischemia. Cell Death Differ 14, 775–784, doi:10.1038/sj.cdd.4402053 (2007).

54 Wang, H., Yu, S. W., Koh, D. W., Lew, J., Coombs, C., Bowers, W., Federoff, H. J., Poirier, G. G., Dawson, T. M. & Dawson, V. L. Apoptosis-inducing factor substitutes for caspase executioners in NMDA-triggered excitotoxic neuronal death. J Neurosci 24, 10963–10973, doi:10.1523/JNEUROSCI.3461-04.2004 (2004).

55 Sun, L. O., Mulinyawe, S. B., Collins, H. Y., Ibrahim, A., Li, Q., Simon, D. J., Tessier-Lavigne, M. & Barres, B. A. Spatiotemporal Control of CNS Myelination by Oligodendrocyte Programmed Cell Death through the TFEB-PUMA Axis. Cell 175, 1811–1826 e1821, doi:10.1016/j.cell.2018.10.044 (2018).

56 Zhang, Y., Chen, K., Sloan, S. A., Bennett, M. L., Scholze, A. R., O’Keeffe, S., Phatnani, H. P., Guarnieri, P., Caneda, C., Ruderisch, N., Deng, S., Liddelow, S. A., Zhang, C., Daneman, R., Maniatis, T., Barres, B. A. & Wu, J. Q. An RNA-sequencing transcriptome and splicing database of glia, neurons, and vascular cells of the cerebral cortex. J Neurosci 34, 11929–11947, doi:10.1523/JNEUROSCI.1860-14.2014 (2014).

57 Zhang, Y., Sloan, S. A., Clarke, L. E., Caneda, C., Plaza, C. A., Blumenthal, P. D., Vogel, H., Steinberg, G. K., Edwards, M. S., Li, G., Duncan, J. A., 3rd, Cheshier, S. H., Shuer, L. M., Chang, E. F., Grant, G. A., Gephart, M. G. & Barres, B. A. Purification and Characterization of Progenitor and Mature Human Astrocytes Reveals Transcriptional and Functional Differences with Mouse. Neuron 89, 37–53, doi:10.1016/j.neuron.2015.11.013 (2016).

58 Clarke, L. E., Liddelow, S. A., Chakraborty, C., Munch, A. E., Heiman, M. & Barres, B. A. Normal aging induces A1-like astrocyte reactivity. Proc Natl Acad Sci U S A 115, E1896–E1905, doi:10.1073/pnas.1800165115 (2018).

59 Bennett, M. L., Bennett, F. C., Liddelow, S. A., Ajami, B., Zamanian, J. L., Fernhoff, N. B., Mulinyawe, S. B., Bohlen, C. J., Adil, A., Tucker, A., Weissman, I. L., Chang, E. F., Li, G., Grant, G. A., Hayden Gephart, M. G. & Barres, B. A. New tools for studying microglia in the mouse and human CNS. Proc Natl Acad Sci U S A 113, E1738–1746, doi:10.1073/pnas.1525528113 (2016).

60 Jung, S., Aliberti, J., Graemmel, P., Sunshine, M. J., Kreutzberg, G. W., Sher, A. & Littman, D. R. Analysis of fractalkine receptor CX(3)CR1 function by targeted deletion and green fluorescent protein reporter gene insertion. Mol Cell Biol 20, 4106–4114, doi:10.1128/MCB.20.11.4106-4114.2000 (2000).

61 Yang, Y., Vidensky, S., Jin, L., Jie, C., Lorenzini, I., Frankl, M. & Rothstein, J. D. Molecular comparison of GLT1+ and ALDH1L1+ astrocytes in vivo in astroglial reporter mice. Glia 59, 200–207, doi:10.1002/glia.21089 (2011).

62 Srinivasan, R., Lu, T. Y., Chai, H., Xu, J., Huang, B. S., Golshani, P., Coppola, G. & Khakh, B. S. New Transgenic Mouse Lines for Selectively Targeting Astrocytes and Studying Calcium Signals in Astrocyte Processes In Situ and In Vivo. Neuron 92, 1181–1195, doi:10.1016/j.neuron.2016.11.030 (2016).

63 Feng, G., Mellor, R. H., Bernstein, M., Keller-Peck, C., Nguyen, Q. T., Wallace, M., Nerbonne, J. M., Lichtman, J. W. & Sanes, J. R. Imaging neuronal subsets in transgenic mice expressing multiple spectral variants of GFP. Neuron 28, 41–51, doi:10.1016/s0896-6273(00)00084-2 (2000).

64 Lautrup, S., Sinclair, D. A., Mattson, M. P. & Fang, E. F. NAD(+) in Brain Aging and Neurodegenerative Disorders. Cell Metab 30, 630–655, doi:10.1016/j.cmet.2019.09.001 (2019).

65 Nakajima, S. & Kunugi, H. Lauric acid promotes neuronal maturation mediated by astrocytes in primary cortical cultures. Heliyon 6, e03892, doi:10.1016/j.heliyon.2020.e03892 (2020).

66 Nonaka, Y., Takagi, T., Inai, M., Nishimura, S., Urashima, S., Honda, K., Aoyama, T. & Terada, S. Lauric Acid Stimulates Ketone Body Production in the KT-5 Astrocyte Cell Line. J Oleo Sci 65, 693–699, doi:10.5650/jos.ess16069 (2016).

67 Saito, M. & Saito, M. Incorporation of very-long-chain fatty acids into sphingolipids of cultured neural cells. J Neurochem 57, 465–469, doi:10.1111/j.1471-4159.1991.tb03774.x (1991).

68 Turner, C. E., Byblow, W. D. & Gant, N. Creatine supplementation enhances corticomotor excitability and cognitive performance during oxygen deprivation. J Neurosci 35, 1773–1780, doi:10.1523/JNEUROSCI.3113-14.2015 (2015).

69 San Martin, A., Arce-Molina, R., Galaz, A., Perez-Guerra, G. & Barros, L. F. Nanomolar nitric oxide concentrations quickly and reversibly modulate astrocytic energy metabolism. J Biol Chem 292, 9432–9438, doi:10.1074/jbc.M117.777243 (2017).

70 Itoh, Y., Esaki, T., Shimoji, K., Cook, M., Law, M. J., Kaufman, E. & Sokoloff, L. Dichloroacetate effects on glucose and lactate oxidation by neurons and astroglia in vitro and on glucose utilization by brain in vivo. Proc Natl Acad Sci U S A 100, 4879–4884, doi:10.1073/pnas.0831078100 (2003).

71 Herrero-Mendez, A., Almeida, A., Fernandez, E., Maestre, C., Moncada, S. & Bolanos, J. P. The bioenergetic and antioxidant status of neurons is controlled by continuous degradation of a key glycolytic enzyme by APC/C-Cdh1. Nat Cell Biol 11, 747–752, doi:10.1038/ncb1881 (2009).

72 Wang, C., Yue, H., Hu, Z., Shen, Y., Ma, J., Li, J., Wang, X. D., Wang, L., Sun, B., Shi, P., Wang, L. & Gu, Y. Microglia mediate forgetting via complement-dependent synaptic elimination. Science 367, 688–694, doi:10.1126/science.aaz2288 (2020).

73 Cao, P., Chen, C., Liu, A., Shan, Q., Zhu, X., Jia, C., Peng, X., Zhang, M., Farzinpour, Z., Zhou, W., Wang, H., Zhou, J. N., Song, X., Wang, L., Tao, W., Zheng, C., Zhang, Y., Ding, Y. Q., Jin, Y., Xu, L. & Zhang, Z. Early-life inflammation promotes depressive symptoms in adolescence via microglial engulfment of dendritic spines. Neuron 109, 2573–2589 e2579, doi:10.1016/j.neuron.2021.06.012 (2021).

74 Camandola, S. & Mattson, M. P. Brain metabolism in health, aging, and neurodegeneration. EMBO J 36, 1474–1492, doi:10.15252/embj.201695810 (2017).

75 Vilchez, D., Ros, S., Cifuentes, D., Pujadas, L., Valles, J., Garcia-Fojeda, B., Criado-Garcia, O., Fernandez-Sanchez, E., Medrano-Fernandez, I., Dominguez, J., Garcia-Rocha, M., Soriano, E., Rodriguez de Cordoba, S. & Guinovart, J. J. Mechanism suppressing glycogen synthesis in neurons and its demise in progressive myoclonus epilepsy. Nat Neurosci 10, 1407–1413, doi:10.1038/nn1998 (2007).

76 Chen, Y., Zhou, S., Zhang, X., Li, D. & Fu, C. Improved fuzzy c-means clustering by varying the fuzziness parameter. Pattern Recognition Letters 157, 60–66, 10.1016/j.patrec.2022.03.017 (2022).

77 Oakley, H., Cole, S. L., Logan, S., Maus, E., Shao, P., Craft, J., Guillozet-Bongaarts, A., Ohno, M., Disterhoft, J., Van Eldik, L., Berry, R. & Vassar, R. Intraneuronal beta-amyloid aggregates, neurodegeneration, and neuron loss in transgenic mice with five familial Alzheimer’s disease mutations: potential factors in amyloid plaque formation. J Neurosci 26, 10129–10140, doi:10.1523/JNEUROSCI.1202-06.2006 (2006).

78 Leng, F. & Edison, P. Neuroinflammation and microglial activation in Alzheimer disease: where do we go from here? Nat Rev Neurol 17, 157–172, doi:10.1038/s41582-020-00435-y (2021).

79 Nibbs, R. J. & Graham, G. J. Immune regulation by atypical chemokine receptors. Nat Rev Immunol 13, 815–829, doi:10.1038/nri3544 (2013).

80 Islam, S. A., Chang, D. S., Colvin, R. A., Byrne, M. H., McCully, M. L., Moser, B., Lira, S. A., Charo, I. F. & Luster, A. D. Mouse CCL8, a CCR8 agonist, promotes atopic dermatitis by recruiting IL-5+ T(H)2 cells. Nat Immunol 12, 167–177, doi:10.1038/ni.1984 (2011).

81 Lu, S. C. Glutathione synthesis. Biochim Biophys Acta 1830, 3143–3153, doi:10.1016/j.bbagen.2012.09.008 (2013).

82 Forman, H. J., Zhang, H. & Rinna, A. Glutathione: overview of its protective roles, measurement, and biosynthesis. Mol Aspects Med 30, 1–12, doi:10.1016/j.mam.2008.08.006 (2009).

83 Wilcock, D. M., Rojiani, A., Rosenthal, A., Levkowitz, G., Subbarao, S., Alamed, J., Wilson, D., Wilson, N., Freeman, M. J., Gordon, M. N. & Morgan, D. Passive amyloid immunotherapy clears amyloid and transiently activates microglia in a transgenic mouse model of amyloid deposition. J Neurosci 24, 6144–6151, doi:10.1523/JNEUROSCI.1090-04.2004 (2004).

84 Ulrich, J. D., Ulland, T. K., Mahan, T. E., Nystrom, S., Nilsson, K. P., Song, W. M., Zhou, Y., Reinartz, M., Choi, S., Jiang, H., Stewart, F. R., Anderson, E., Wang, Y., Colonna, M. & Holtzman, D. M. ApoE facilitates the microglial response to amyloid plaque pathology. J Exp Med 215, 1047–1058, doi:10.1084/jem.20171265 (2018).

85 Aoyama, C., Liao, H. & Ishidate, K. Structure and function of choline kinase isoforms in mammalian cells. Prog Lipid Res 43, 266–281, doi:10.1016/j.plipres.2003.12.001 (2004).

86 Tzika, A. A., Astrakas, L., Cao, H., Mintzopoulos, D., Andronesi, O. C., Mindrinos, M., Zhang, J., Rahme, L. G., Blekas, K. D., Likas, A. C., Galatsanos, N. P., Carroll, R. S. & Black, P. M. Combination of high-resolution magic angle spinning proton magnetic resonance spectroscopy and microscale genomics to type brain tumor biopsies. Int J Mol Med 20, 199–208 (2007).

87 Southan, G. J., Szabo, C. & Thiemermann, C. Inhibition of the induction of nitric oxide synthase by spermine is modulated by aldehyde dehydrogenase. Biochem Biophys Res Commun 203, 1638–1644, doi:10.1006/bbrc.1994.2374 (1994).

88 Kaczmarek, L., Kaminska, B., Messina, L., Spampinato, G., Arcidiacono, A., Malaguarnera, L. & Messina, A. Inhibitors of polyamine biosynthesis block tumor necrosis factor-induced activation of macrophages. Cancer Res 52, 1891–1894 (1992).

89 Choi, Y. H. & Park, H. Y. Anti-inflammatory effects of spermidine in lipopolysaccharide-stimulated BV2 microglial cells. J Biomed Sci 19, 31, doi:10.1186/1423-0127-19-31 (2012).

90 Bonvento, G. & Bolanos, J. P. Astrocyte-neuron metabolic cooperation shapes brain activity. Cell Metab 33, 1546–1564, doi:10.1016/j.cmet.2021.07.006 (2021).

91 Lerchundi, R., Fernandez-Moncada, I., Contreras-Baeza, Y., Sotelo-Hitschfeld, T., Machler, P., Wyss, M. T., Stobart, J., Baeza-Lehnert, F., Alegria, K., Weber, B. & Barros, L. F. NH4(+) triggers the release of astrocytic lactate via mitochondrial pyruvate shunting. Proc Natl Acad Sci U S A 112, 11090–11095, doi:10.1073/pnas.1508259112 (2015).

92 Ioannou, M. S., Jackson, J., Sheu, S. H., Chang, C. L., Weigel, A. V., Liu, H., Pasolli, H. A., Xu, C. S., Pang, S., Matthies, D., Hess, H. F., Lippincott-Schwartz, J. & Liu, Z. Neuron-Astrocyte Metabolic Coupling Protects against Activity-Induced Fatty Acid Toxicity. Cell 177, 1522–1535 e1514, doi:10.1016/j.cell.2019.04.001 (2019).

93 Chen, X. & Gan, B. SLC25A39 links mitochondrial GSH sensing with iron metabolism. Mol Cell 84, 616–618, doi:10.1016/j.molcel.2023.12.037 (2024).

94 Aliaga, M. E., Lopez-Alarcon, C., Bridi, R. & Speisky, H. Redox-implications associated with the formation of complexes between copper ions and reduced or oxidized glutathione. J Inorg Biochem 154, 78–88, doi:10.1016/j.jinorgbio.2015.08.005 (2016).

95 Wang, Y., Yen, F. S., Zhu, X. G., Timson, R. C., Weber, R., Xing, C., Liu, Y., Allwein, B., Luo, H., Yeh, H. W., Heissel, S., Unlu, G., Gamazon, E. R., Kharas, M. G., Hite, R. & Birsoy, K. SLC25A39 is necessary for mitochondrial glutathione import in mammalian cells. Nature 599, 136–140, doi:10.1038/s41586-021-04025-w (2021).

96 Zhang, T., Liu, Q., Gao, W., Sehgal, S. A. & Wu, H. The multifaceted regulation of mitophagy by endogenous metabolites. Autophagy 18, 1216–1239, doi:10.1080/15548627.2021.1975914 (2022).

97 Mak, T. W., Grusdat, M., Duncan, G. S., Dostert, C., Nonnenmacher, Y., Cox, M., Binsfeld, C., Hao, Z., Brustle, A., Itsumi, M., Jager, C., Chen, Y., Pinkenburg, O., Camara, B., Ollert, M., Bindslev-Jensen, C., Vasiliou, V., Gorrini, C., Lang, P. A., Lohoff, M., Harris, I. S., Hiller, K. & Brenner, D. Glutathione Primes T Cell Metabolism for Inflammation. Immunity 46, 675–689, doi:10.1016/j.immuni.2017.03.019 (2017).

98 Chen, Y., Manna, S. K., Golla, S., Krausz, K. W., Cai, Y., Garcia-Milian, R., Chakraborty, T., Chakraborty, J., Chatterjee, R., Thompson, D. C., Gonzalez, F. J. & Vasiliou, V. Glutathione deficiency-elicited reprogramming of hepatic metabolism protects against alcohol-induced steatosis. Free Radic Biol Med 143, 127–139, doi:10.1016/j.freeradbiomed.2019.07.025 (2019).

99 Liu, M., Zhang, H., Xie, Z., Huang, Y., Sun, G., Qi, D., Furey, A., Randell, E. W., Rahman, P. & Zhai, G. Glutathione, polyamine, and lysophosphatidylcholine synthesis pathways are associated with circulating pro-inflammatory cytokines. Metabolomics 18, 76, doi:10.1007/s11306-022-01932-5 (2022).

100 Gulenturk, C., Alp-Turgut, F. N., Arikan, B., Tofan, A., Ozfidan-Konakci, C. & Yildiztugay, E. Polyamine, 1,3-diaminopropane, regulates defence responses on growth, gas exchange, PSII photochemistry and antioxidant system in wheat under arsenic toxicity. Plant Physiol Biochem 201, 107886, doi:10.1016/j.plaphy.2023.107886 (2023).

101 Hohsfield, L. A., Najafi, A. R., Ghorbanian, Y., Soni, N., Crapser, J., Figueroa Velez, D. X., Jiang, S., Royer, S. E., Kim, S. J., Henningfield, C. M., Anderson, A., Gandhi, S. P., Mortazavi, A., Inlay, M. A. & Green, K. N. Subventricular zone/white matter microglia reconstitute the empty adult microglial niche in a dynamic wave. Elife 10, doi:10.7554/eLife.66738 (2021).

102 Ai, D., Qiu, B., Chen, X. J., Li, F., Yao, D., Mi, H., Li, J. L., Zhou, B., Zuo, J., Wang, Y., Ge, W. P. & Sun, W. Inhibition of microglial Slc2a5 attenuates ischemic brain injury. Metabolism 175, 156429, doi:10.1016/j.metabol.2025.156429 (2026).

103 Zhang, T., Ai, D., Wei, P., Xu, Y., Bi, Z., Ma, F., Li, F., Chen, X. J., Zhang, Z., Zou, X., Guo, Z., Zhao, Y., Li, J. L., Ye, M., Feng, Z., Zhang, X., Zheng, L., Yu, J., Li, C., Tu, T., Zeng, H., Lei, J., Zhang, H., Hong, T., Zhang, L., Luo, B., Li, Z., Xing, C., Jia, C., Li, L., Sun, W. & Ge, W. P. The subcommissural organ regulates brain development via secreted peptides. Nat Neurosci 27, 1103–1115, doi:10.1038/s41593-024-01639-x (2024).

104 Li, J. L., Bi, Z., Chen, X. J., Ming, T., Qiu, B., Li, F., Feng, Z., Ai, D., Zhang, T., Wang, J., Lin, S., Lu, Y., Wang, Z., Huang, J., Zhao, F., Zhao, H., Wang, Y., Sun, W. & Ge, W. P. A targeted vector for brain endothelial cell gene delivery and cerebrovascular malformation modelling. Nat Biomed Eng, doi:10.1038/s41551-025-01538-x (2025).

105 Lee, M. R., Suh, H. R., Kim, M. W., Cho, J. Y., Song, H. K., Jung, Y. S., Hwang, D. Y. & Kim, K. S. Comparison of the anesthetic effects of 2,2,2-tribromoethanol on ICR mice derived from three different sources. Lab Anim Res 34, 270–278, doi:10.5625/lar.2018.34.4.270 (2018).

106 Brewer, G. J. & Torricelli, J. R. Isolation and culture of adult neurons and neurospheres. Nature Protocols 2, 1490–1498, doi:10.1038/nprot.2007.207 (2007).

107 Ni, M., Solmonson, A., Pan, C., Yang, C., Li, D., Notzon, A., Cai, L., Guevara, G., Zacharias, L. G., Faubert, B., Vu, H. S., Jiang, L., Ko, B., Morales, N. M., Pei, J., Vale, G., Rakheja, D., Grishin, N. V., McDonald, J. G., Gotway, G. K., McNutt, M. C., Pascual, J. M. & DeBerardinis, R. J. Functional Assessment of Lipoyltransferase-1 Deficiency in Cells, Mice, and Humans. Cell Rep 27, 1376–1386 e1376, doi:10.1016/j.celrep.2019.04.005 (2019).

108 Pang, H., Li, J.-L., Hu, X.-L., Chen, F., Gao, X., Zacharias, L. G., Cai, F., DeBerardinis, R. J., Sun, W., Hu, Z. & Ge, W.-p. Precision mapping of the mouse brain metabolome. bioRxiv (2020).

109 Andrews., S. FASTQC. A quality control tool for high throughput sequence data. http://www.bioinformatics.babraham.ac.uk/projects/fastqc/. (2014).

110 Chen, S., Zhou, Y., Chen, Y. & Gu, J. fastp: an ultra-fast all-in-one FASTQ preprocessor. Bioinformatics 34, i884–i890, doi:10.1093/bioinformatics/bty560 (2018).

111 Dobin, A., Davis, C. A., Schlesinger, F., Drenkow, J., Zaleski, C., Jha, S., Batut, P., Chaisson, M. & Gingeras, T. R. STAR: ultrafast universal RNA-seq aligner. Bioinformatics 29, 15–21, doi:10.1093/bioinformatics/bts635 (2013).

112 Love, M. I., Huber, W. & Anders, S. Moderated estimation of fold change and dispersion for RNA-seq data with DESeq2. Genome Biol 15, 550, doi:10.1186/s13059-014-0550-8 (2014).

113 Huan, T., Forsberg, E. M., Rinehart, D., Johnson, C. H., Ivanisevic, J., Benton, H. P., Fang, M., Aisporna, A., Hilmers, B., Poole, F. L., Thorgersen, M. P., Adams, M. W. W., Krantz, G., Fields, M. W., Robbins, P. D., Niedernhofer, L. J., Ideker, T., Majumder, E. L., Wall, J. D., Rattray, N. J. W., Goodacre, R., Lairson, L. L. & Siuzdak, G. Systems biology guided by XCMS Online metabolomics. Nat Methods 14, 461–462, doi:10.1038/nmeth.4260 (2017).

114 Tautenhahn, R., Patti, G. J., Rinehart, D. & Siuzdak, G. XCMS Online: a web-based platform to process untargeted metabolomic data. Anal Chem 84, 5035–5039, doi:10.1021/ac300698c (2012).

115 Chong, J., Soufan, O., Li, C., Caraus, I., Li, S., Bourque, G., Wishart, D. S. & Xia, J. MetaboAnalyst 4.0: towards more transparent and integrative metabolomics analysis. Nucleic Acids Res 46, W486–W494, doi:10.1093/nar/gky310 (2018).

116 Xiong, N., Gao, X., Zhao, H., Cai, F., Zhang, F. C., Yuan, Y., Liu, W., He, F., Zacharias, L. G., Lin, H., Vu, H. S., Xing, C., Yao, D. X., Chen, F., Luo, B., Sun, W., DeBerardinis, R. J., Xu, H. & Ge, W. P. Using arterial-venous analysis to characterize cancer metabolic consumption in patients. Nat Commun 11, 3169, doi:10.1038/s41467-020-16810-8 (2020).

117 Qu, P., Rom, O., Li, K., Jia, L., Gao, X., Liu, Z., Ding, S., Zhao, M., Wang, H., Chen, S., Xiong, X., Zhao, Y., Xue, C., Zhao, Y., Chu, C., Wen, B., Finney, A. C., Zheng, Z., Cao, W., Zhao, J., Bai, L., Zhao, S., Sun, D., Zeng, R., Lin, J., Liu, W., Zheng, L., Zhang, J., Liu, E. & Chen, Y. E. DT-109 ameliorates nonalcoholic steatohepatitis in nonhuman primates. Cell Metab 35, 742–757 e710, doi:10.1016/j.cmet.2023.03.013 (2023).

118 Schonberger, K., Mitterer, M., Glaser, K., Stecher, M., Hobitz, S., Schain-Zota, D., Schuldes, K., Lammermann, T., Rambold, A. S., Cabezas-Wallscheid, N. & Buescher, J. M. LC-MS-Based Targeted Metabolomics for FACS-Purified Rare Cells. Anal Chem 95, 4325–4334, doi:10.1021/acs.analchem.2c04396 (2023).

